# Novel Aromatic Polyhydroxyalkanoates from Engineered *Cupriavidus necator* H16: Expanding Bio-Polyester Synthesis from Renewable Feedstocks

**DOI:** 10.1101/2021.12.12.472320

**Authors:** Nils J.H. Averesch, Vince E. Pane, Frauke Kracke, Sulogna Chatterjee, Aaron J. Berliner, Marika Ziesack, Shannon N. Nangle, Robert M. Waymouth, Craig S. Criddle

**Affiliations:** Center for the Utilization of Biological Engineering in Space (CUBES), Berkeley, CA, USA; Department of Civil and Environmental Engineering, Stanford University, CA, USA; Department of Chemistry, Stanford University, CA, USA; Department of Bioengineering, University of California, Berkeley, Berkeley, CA, USA; Wyss Institute for Biologically Inspired Engineering, MA, USA

**Author notes:** Corresponding authors NJHA.

**Keywords:** Polyhydroxyalkanoates, aromatic polyesters, bioplastics, single-carbon (C-1) feedstocks, microbial electrosynthesis

## Abstract

Synthetic materials are integral components of consumable and durable goods and are indispensable in the modern world. Polyesters are the most versatile bulk- and specialty-polymers but their production is not sustainable and their fate at end-of-life is of great concern. Bioplastics, though highly regarded, often lag behind conventional plastics due to poor material properties and a competitive market. This has limited the success of sustainable replacements at scale. Enabling the production of bioplastics with superior properties from waste-derived feedstocks could change that.

To this end, we created a synthetic entry into the metabolic pathway of bio-polyester synthesis of *Cupriavidus necator* H16 employing heterologous hydroxyacyl-CoA transferase and mutant PHA synthase. The resulting microbial cell factories enabled the co-polymerization of a range of aliphatic and aromatic hydroxy carboxylates, including a hydroxyphenylic and a hydroxyfuranoic acid, for the first time incorporating aromatic rings in the backbone of biological polyesters. The latter were structurally analogous to synthetic polyesters like PET and PEF, as well as PBST and PBAT. The obtained bio-polyesters underwent characterization for specific physicochemical properties, complemented by the prediction of uncharacterized physicochemical and mechanical properties. In a further advance, the transgenic strain was cultivated in a bio-electrochemical system under autotrophic conditions, enabling synthesis of aromatic bio-polyesters with *in situ* generated H_2_+O_2_, while assimilating CO_2_. Employing elementary flux-mode analysis, metabolic modeling confirmed the feasibility of producing an extended range of aliphatic, arylatic, and aromatic PHAs *de novo* from various feedstocks. This comprehensive study paves the way toward sustainable bio-production of advanced high-performance thermoplastics and thermosets.

**Significance Statement:** Biomaterials can help the chemical industry transition to a carbon-neutral and circular economy, reducing the accumulation of greenhouse gases and plastic waste by developing biological replacements for fossil carbon-based plastics and implementing end-of-life strategies.

Accomplished via the genetic engineering of a microbe that can assimilate carbon dioxide, this work demonstrates the first biocatalytic polymerization and incorporation of aromatic building blocks into the backbone of polyhydroxyalkanoates (PHAs). Employing a bio-electrochemical system for cultivation of the microbes, oxyhydrogen is formed and consumed *in situ*, thus avoiding explosive gas mixtures.

The obtained aromatic PHAs are structurally similar and functionally analogs to synthetic bulk- and high-performance polymers such as PET (polyethylene terephthalate) and PEF (polyethylene furanoate) as well as PBST (polybutylene succinate-co-terephthalate) and PBAT (polybutylene adipate-co-terephthalate).

**Graphical Abstract:** The biosynthesized novel aromatic polyhydroxyalkanoates described in this study are structurally similar to incumbent synthetic polyesters such as poly(ethylene terephthalate) (PET, orange box) and poly(ethylene furan-dicarboxylate) (PEF, yellow box). While PET and PEF may be bio-based to varying degrees, they are always synthesized chemically. In contrast, not only the monomers of poly(hydroxybutyrate-co-phloretate) and poly(hydroxybutyrate-co-hydroxymethylfuranoate) (green box) are fully bio-derived, but also the entire polymers are synthesized biologically. This was demonstrated by employing the autotrophic bacterium *Cupriavidus necator* H16.

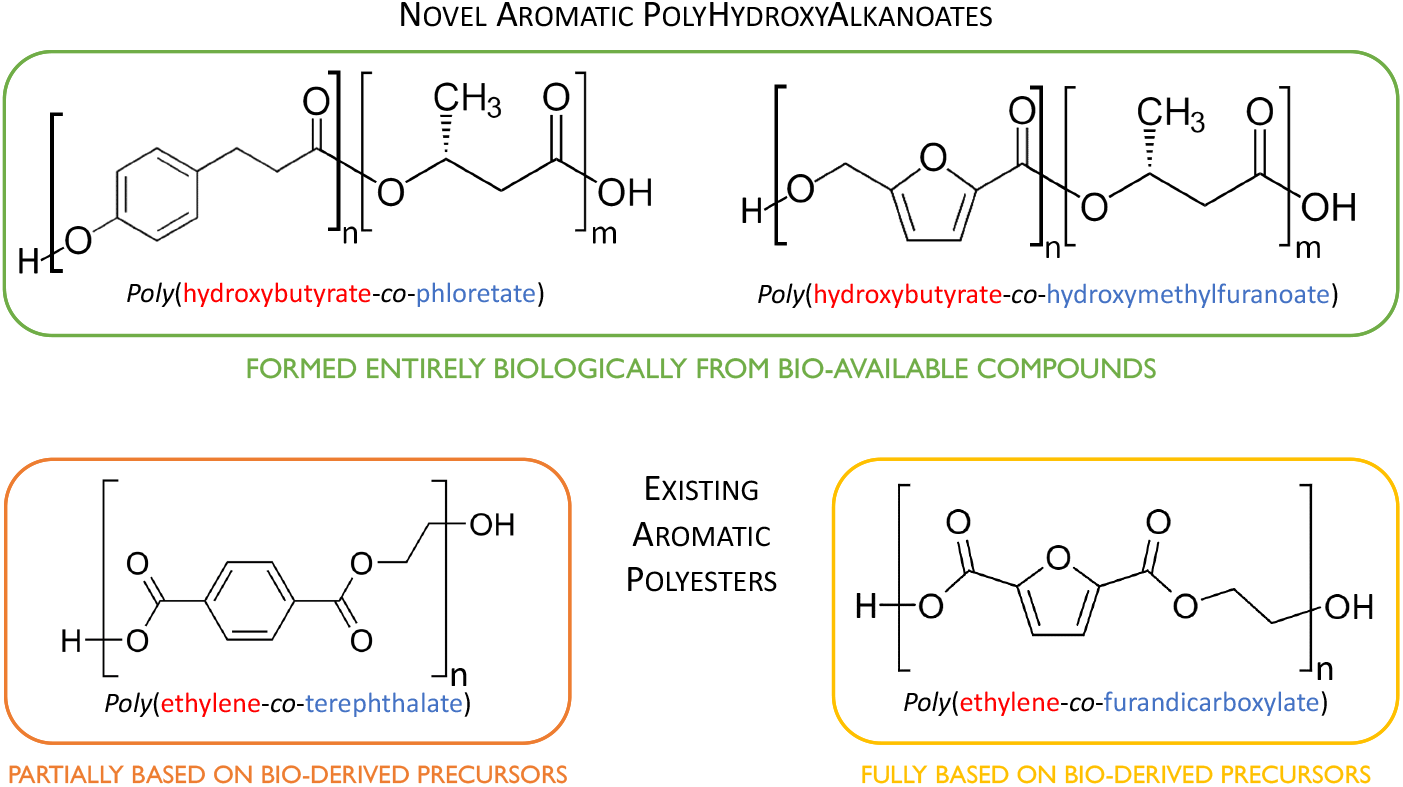

**Highlights:**

- First-ever biocatalytic formation of polyesters with aromatic rings in the backbone
- Microbial production of aromatic PHAs in a bio-electrochemical system on CO_2_ and H_2_+O_2_
- Expression-level of PHA synthase and molecular weight of polyesters are inversely correlated
- *In silico* design and pathway analysis of bio-polyesters from low-cost carbon-feedstocks
- Prediction of material properties for novel aromatic co-polyesters and structural analogs

## 1. Introduction

In 2023, the then-incumbent Administration of the United States of America defined the bold goal to, within 20 years, “demonstrate and deploy cost-effective and sustainable routes to convert bio-based feedstocks into recyclable-by-design polymers that can displace more than 90% of today’s plastics and other commercial polymers at scale”^1^. Retraction of this policy by the current Administration does not avert a global plastics crisis, but aggravates the remedy thereof.

### 1.1 Background

Modern society is based upon ever-increasing consumption of energy and material resources, accompanied by the emission of waste and pollutants^2^. Greenhouse gases are reaching critical levels, and pollution from the fabrication of recalcitrant synthetic materials is ubiquitous and undeniable^3^. A lifestyle and an economy dependent on plastic consumption risk serious harm to the biosphere^4^. To ensure a viable planet for generations to come, methods are needed to reduce waste accumulation while embracing readily available and renewable feedstocks^5^. Although plastic recycling is necessary for a circular economy, it will not be able to fully prevent the unintended release of synthetic materials into the environment^6^. This is of particular relevance in single-use applications where the long-term stability of plastics is as unnecessary as problematic^7^. While bio-degradable materials derived from renewable resources or from upcycling of existing materials are desirable^8^, production from cultivated feedstocks is not always cost-effective nor necessarily carbon-negative or -neutral^9^. An alternative is the utilization of ultra-low-cost (waste) carbon sources, such as C1-feedstocks, for the production of materials, which can directly mitigate greenhouse gas emissions while avoiding competition with the food-industry^10^. Such a strategy also improves commercial competitiveness by decreasing feedstock-cost^11^. Especially useful are compounds that can substitute for or directly replace industrial (petrochemistry-based) polymers. Such bio-replacements are attractive because they do not require extensive modifications to the existing infrastructure for processing and manufacturing^12^.

Polyhydroxybutyrate (PHB), which belongs to the class of polyhydroxyalkanoates (PHAs), is a thermoplastic polyester that can be biologically produced outgoing from non-edible, waste-derived carbon sources, such as carbon dioxide or methane, and could be a bio-degradable replacement for synthetic materials^13^. To date, however, insufficient material properties^14^ and high cost of production^11^ have hampered its widespread adoption by the chemical manufacturing industry. To explore possible remedies, a plethora of PHAs have been generated from alternative co-monomers, enabling the tuning of the material properties^14^: while linear short-chain PHAs, have high-strength and -flexibility^15–17^, aromatic groups in the side chain lead to more amorphous polymers with increased glass transition and degradation temperatures, conferring a wider processing range^18^. For example, polymers of mandelic and phenyllactic acid, poly(mandelite)^19^ and poly(phenyllactide)^20^, respectively, have been chemically synthesized and characterized. These PHAs exhibited properties mimicking those of polystyrene. However, compared to the properties of existing high-performance plastics, these improvements are rather incremental and most PHAs still fall short of even basic bulk materials.

All biocatalytically produced PHAs that contain aromatic rings only bear these on a side^18^. Incorporation of aromatic rings into the bio-polyester’s backbone could vastly improve material properties: such polymers would conceivably resemble synthetic aromatic polyesters like polyethylene terephthalate (PET), polyethylene furanoate (PEF), or even polyarylates. Analogous polymers based on bioderivable building blocks have been chemically synthesized and characterized, such as co-polyesters of glycolic and *para*-hydroxybenzoic acid^21^, as well as 3-(*para*-hydroxyphenyl)propionic (phloretic) or 3-(*para*-hydroxyphenyl)acrylic (coumaric) acid^22^. These “polyhydroxyarylates” exhibit liquid-crystal polymer properties; full biosynthesis of such polyesters could open the door to a new realm of renewable materials with superior properties^18^.

### 1.2 Concept

To explore the potential for production of advanced bio-polyesters from sustainable feedstocks, this study employs the ‘knallgas’ bacterium *Cupriavidus necator* H16 as a host. This well-characterized mixotroph utilizes a wide range of carbon- and energy sources, grows rapidly to high cell density^23^, and is an excellent producer of PHB^24^. As a chemolithoautotroph, *C. necator* H16 has previously been engineered for *de novo* production of non-natural bio-polyesters from carbon dioxide, including short and medium chain-length PHAs^25^. Polymerization of exogenously supplied aromatic precursors has also been explored^26^.

Here, a PHA synthase-deficient (Δ*phaC1*) knockout mutant of *C. necator* H16 was complemented with the genes coding for a hydroxyacyl-CoA transferase (*pct540* or *hadA* from *Clostridium propionicum* or *Clostridium difficile*, respectively) and a mutant PHA synthase (*phaC1437* from *Pseudomonas* sp. MBEL 6-19). The CoA transferases exhibit complementary substrate scopes, enabling activation of several non-canonical PHA precursors; notably, *hadA* has been shown to also activate hydroxybenzoic acids^27,28^. The endogenous *phaC1* of *C. necator* H16 has limited substrate specificity and was therefore replaced with *phaC1437*, which can efficiently polymerize several small and large hydroxy acids, including arylatic compounds, as demonstrated in *E. coli*^29^. This created a second entry into the PHA biosynthesis of *C. necator* (Fig. 1a), enabling the formation of several straight-chain aliphatic, and arylatic, as well as previously unknown aromatic PHAs.

**Figure 1.**
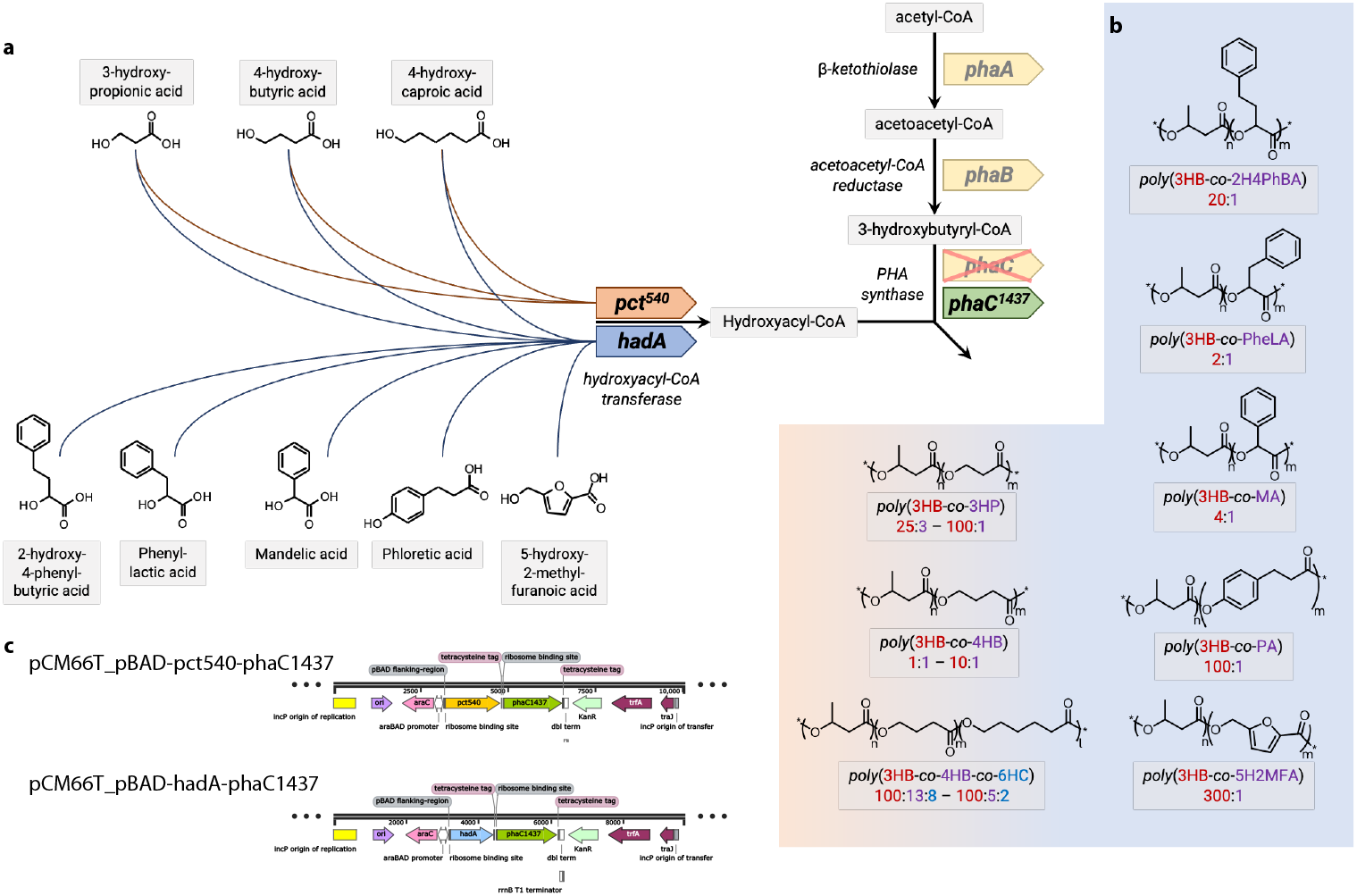
Partial breakdown of the pathway for biosynthesis of novel co-polyesters. **(a)**. The exogenous segment (horizontal) feeds non-canonical monomers to the endogenous pathway of poly(3-hydroxybutyrate) biosynthesis (vertical) with the genes responsible for the natural biosynthetic route represented in yellow. The genes coding for the alternative heterologous CoA transferases (*pct540* and *hadA*, represented in orange and blue, respectively) complement the route, activating the hydroxy-carbonic acids (by forming thiols thereof) with Coenzyme A. The mutant PHA synthase (*phaC1437*, represented in green) substitutes the natural PHA synthase (*phaC1*), enabling polymerization of the alternative polymer precursors. Together the two metabolic routes mediate the formation of novel polyhydroxyalkanoates **(b)** where “m”, “n”, and “l” indicate the uncorrelated statistic distribution of the co-monomers (3HB = 3-hydroxybutyrate, 4HB = 4-hydroxybutyrate, 6HC = 6-hydroxycaproate, PheLA = phenyllactate, PA = phloretate, 5HM2F = 5-hydroxymethyl-2-furanoate) throughout the chain. The composition of the obtained respective polymers is indicated as molar ratio under the structural designation. For the aliphatic PHAs, the first (left) ratio corresponds to the strain carrying the *pct540*, while the second (right) ratio corresponds to the strain carrying the *hadA*. The arylaliphatic and arylatic PHAs were only obtained with the *hadA*. The essential heterologous biochemical functions are conferred by two genetic vectors **(c)**, alternatively harbored by *Cupriavidus necator* H16 Δ*phaC1*.

In addition to the experimental demonstration of co-polymer synthesis from exogenously supplied precursors, we further explored the feasibility of their *de novo* production from various carbon sources using *in silico* metabolic modeling. This system-level analysis revealed the theoretical potential for high-yield production of a variety of aliphatic and aromatic PHAs from a range of feedstocks, including single-carbon compounds. Moreover, we demonstrated the formation of arylatic PHAs during autotrophic growth in a bio-electrochemical system, corroborating the relevance of our concept for the production of aromatic polyesters from carbon dioxide. Finally, we predicted key physicochemical and mechanical properties of the obtained and hypothetical polymers, revealing promising material characteristics that approach those of commercial synthetic polyesters.

## 2 Results and Discussion

Introduction of *phaC1437* into *C. necator* H16 Δ*phaC1* rescued the PHA-negative phenotype of the knockout mutant. The PHA-content [w_PHA_/w_CDW_] of the engineered strains was similar to the H16 wildtype (∼ 44% for Δ*phaC1 pct540*-*phaC1437* and ∼ 58% for Δ*phaC1 hadA*-*phaC1437*, see Tab. S3). Notably, bio-polyesters were formed independently from induction of expression of the heterologous genes under control of P_BAD_, however, composition was dependent on induction. While this represented a peculiar observation, the primary objective was to investigate the spectrum of PHAs that can be obtained from augmenting metabolism with the non-natural biochemical functions.

### 2.1 Toxicity of Polyester-Precursors

When commencing feeding experiments with various hydroxyacyls, some of the polymers’ precursors appeared to exhibit toxicity leading to impairment of growth. In particular, 2-hydroxy-4-phenylbutyric and phloretic acid could only be supplied at 10 mM and 5 mM, respectively. This dictated the maximum employable concentrations (see Extended Data 1 for comprehensive screening of all employed compounds). While not toxic, poor solubility of 6-hydroxycaproic acid (*<* 20 mM) limited the final concentration of the ω-fatty acid in the growth medium. Hence, aliphatic, aryl-aliphatic, and aromatic precursors were supplied at concentrations of 20 mM, 10 mM, and 5 mM, respectively.

### 2.2 Diversity of Obtained Non-Natural Bio-Polyesters

Several rounds of batch shakeflask cultivations on minimal medium containing different hydroxy-carbonic acids were conducted. The composition of the polyesters formed by the engineered strains and the wildtype of *C. necator* H16 as well as the respective fraction of the different repeat units in the co-polymer were determined by means of NMR spectroscopy (^1^H-, in certain cases complemented with ^13^C-, see Extended Data 2). The range of the obtained PHAs is shown in Fig. 1b, while the entire spectrum and composition of all formed co-polyesters in dependence of the strain and the concentration of exogenously provided precursors is consolidated in Fig. S5.

Aromatic hydroxy acids were polymerized only by the transgenic variant of *C. necator* H16 carrying *hadA* (and *phaC1437*) (Fig. 1c). Most significantly, the hydroxyphenylic compound ‘phloretic acid’ (PA) as well as hydroxyfuranoic compound 5-hydroxymethyl-2-furanoic acid (5HM2F), also known as ‘sumiki’s acid’, were successfully incorporated into co-polymers with 3-hydroxybutyrate (3HB). Although the obtained polymers only contained small fractions of the aromatic compounds no higher than 1%, this is the first time that incorporation of aromatic rings into the backbone of biological polyesters is demonstrated, paving the way to a new class of fully biologically formed materials: the obtained co-polymers, P(3HB-co-PA) and P(3HB-co-5HM2F), bear structural analogy to synthetic plastics like polyethylene terephthalate (PET) and polyethylene furanoate (PEF)^30^, respectively. Not only does this bring the truly sustainable production of such plastics into reach, but it also creates the tantalizing perspective for biocatalytic synthesis of high-performance poly(hydroxy)arylates^21,22^.

PHAs obtained from aryl-alkyl hydroxy acids had significant fractions of aromatics-containing repeat units, which varied depending on the precursor: with mandelate (MA) about 20% of the obtained polyester was composed of the non-canonical monomers; incorporation of phenyllactate (PheLA) yielded a PHA where approx. every third repeat unit contained an aromatic residue; from 2-hydroxy-4-phenylbutyrate (2H4PheB) a co-polymer with a roughly one order of magnitude lower fraction of the exogenous precursor was obtained.

With aliphatic hydroxy acids both engineered strains as well as the H16 wildtype formed non-natural PHAs. From 6-hydroxycaproic acid (6HC) the wildtype produced co-polymers containing 4-hydroxybutyrate (4HB), while PHA derived from the engineered strains contained both 4HB and 6HC. In all cases, the presence of the 4HB repeat units is attributed to the original C6 ω-fatty acid undergoing β-oxidation down to a C4 ω-fatty acid before being incorporated into the polymer. While the wildtype apparently degraded all of the 6HC, this was prevented in the transgenic strains, presumably by the heterologous enzymes: accelerated kinetics due to high abundance of the mutant PHA synthase, in concert with the over-expressed CoA transferase may have partially out-competed β-oxidation. Incorporation-level of 4HB by the engineered strain that carried *hadA* (and *phaC1437*) vs. by the H16 wildtype was similar (∼ 10%). This indicates that the natural PHA synthase (*phaC1*) is promiscuous enough to polymerize the non-canonical monomer while *phaC1437* offers no significant advantage for formation of poly(4HB). Consequentially, also a native CoA transferase must exist that accepts 4HB as a substrate at least as well as *hadA*; the respective function is likely provided by the acetate CoA transferase ‘YdiF’ (*pct*), as previously reported^31^. Contrarily, the strain carrying *pct540* (and *phaC1437*) formed a co-polymer with a high fraction of 4HB (approx. 66%, as per ^1^H-NMR). Further analytics (^13^C-NMR and GPC trace, see Extended Data 3 & 4) indicated that the obtained polyester was a block-co-polymer^32^.

While comparatively low, the incorporation ratio of 3-hydroxypropionate (3HP) was also much higher in the case of the strain that carried *pct540* with respect to the strain that carried *hadA*. Further, no co-polymers containing 2-hydroxypropionic (lactic) and 2-hydroxyacetic (glycolic) acid were obtained from any of the strains. It is well known that volatile fatty acids (VFAs) are good substrates for *C. necator*, which includes 3HP^33^ as well as lactic and glycolic acid^34–36^. Hence, it is likely that these were catabolized before they could be polymerized. This conclusion is supported by observations from the growth challenge assays (Extended Data 1) where biomass density on the gradient-agar plates was increased towards higher concentrations of the respective compounds.

Without exogenously provided precursors, PHAs formed by the engineered strains were mainly composed of 3HB repeat units with traces of other constituents, the occurrence of which depended on the employed heterologous CoA transferase gene (*pct540* or *hadA*) and level of induction thereof (see Fig. E2D and E2E).

### 2.3 Thermal Properties of Non-Natural Bio-Polyesters

Polymers that contained a significant fraction of non-canonical repeat units were subjected to Gel-Permeation Chromatography (GPC, see Extended Data 3 for traces) and Differential Scanning Calorimetry (DSC, see Extended Data 4 for profiles), to determine the impact of the co-monomers on the molecular weight distribution as well as thermal properties of the PHAs. Compared to the appropriate controls (i.e., PHAs obtained from wildtype and mutant strains not supplemented with non-canonical polymer precursors), a selection of the most significant data from GPC and DSC is compiled in Tab. 1; the molecular weight and associated polydispersity (PDI) of all polymers are provided in Tab. S1.

**Table 1.** Thermal properties and molecular weights of a subset of the PHAs obtained from engineered *C. necator* H16. ^*a*^ natural P3HB, ^*b*,*c*^ P3HB with minor fraction of indeterminate hydroxycarbonate of natural origin (see Fig. E2D & E2E). Abbreviations: P = *poly*-, 3HB = 3-hydroxybutyrate, 4HB = 4-hydroxybutyrate, 6HC = 6-hydroxycaproate, PheLA = phenyllactate, PA = phloretate, 5HM2F = 5-hydroxymethyl-2-furanoate; T_*g*_ = glass transition temperature, T_*m*_ = melting point, ΔH_*c*_ = enthalpy of crystallization, M_*n*_ = number average molecular weight, M_*w*_ = weight average molecular weight, PDI = polydispersity index (M_*w*_/M_*n*_), n.a. = not applicable, N/A = not available.

| Strain | Induced | Polymer | $T_g$<br>(°C) | $T_m$<br>(°C) | $\Delta H_c$<br>(J/g) | $M_n$<br>(kg/mol) | PDI<br>(mol/mol) |
| --- | --- | --- | --- | --- | --- | --- | --- |
| H16 (wildtype) | n.a. | P3HB<sup>a</sup> | 4 | 170 | 19 | 397.4 | 1.71 |
| H16 Δ*phaC1* *pct540* *phaC1437* | no | P3HB | 3 | 170 | 29 | 69.2 | 1.94 |
|  | yes | P3HB<sup>b</sup> | 1 | 163 | 43 | 45.3 | 2.72 |
| H16 Δ*phaC* *hadA* *phaC1437* | no | P3HB | 4 | 169 | 42 | 54.8 | 2.38 |
|  | yes | P3HB<sup>c</sup> | 1 | 164 | 34 | 10.9 | 2.54 |
| H16 Δ*phaC1* *pct540* *phaC1437* | yes | P(3HB-co-4HB)<br>~ 1:1 | -24 | 160 | 27 | 25.2 | 2.94 |
| H16 Δ*phaC* *hadA* *phaC1437* | yes | P(3HB-co-4HB-co-6HC)<br>~ 100:5:2 | -5 | 129 | 6 | 11.8 | 2.13 |
|  | yes | P(3HB-co-PheLA)<br>~ 2:1 | 5 | N/A | 0 | 13.2 | 1.65 |
|  | yes | P(3HB-co-PA)<br>~ 500:1 | 1 | 155 | 55 | 14.5 | 2.33 |
|  | yes | P(3HB-co-5HM2F)<br>~ 300:1 | -1 | 159 | 53 | 13.7 | 3.28 |

The molecular weights of the co-polymers formed with exogenously supplied precursors were on the same order of magnitude as PHA obtained from the non-supplemented cultures. The PDI of non-natural polyesters was generally higher than that of natural PHB (natural *<* 2 *<* non-natural) and peaked when expression of the heterologous genes was fully induced.

The glass transition- and melting temperatures (T_*m*_ and T_*g*_, respectively) of the P3HB generated from both the *pct540*- and the *hadA*-strain were comparable to those of natural P3HB (∼170^°^C and ∼4^°^C, respectively), whereas polymers generated with induction of the strains exhibited slightly lower T_*m*_ and T_*g*_ (∼163^°^C and ∼1^°^C, respectively). These slightly depressed melting points are attributed to small amounts of alternative co-monomers of natural origin incorporated upon high expression of the heterologous genes^37^, as evidenced by minor resonances in the ^1^H-NMR spectra (Fig. E2D & E2E).

The high melting point (160^°^C) and relatively high heat of fusion (ΔH_*c*_ ∼27 J/g, similar to P3HB), coupled with the low T_*g*_ (∼24^°^C, similar to P4HB) of the P(3HB-co-4HB) co-polymer derived from *C. necator* H16 Δ*phaC1 pct540*-*phaC1437* provide strong evidence for a block-co-polymer structure of the 3HB and 4HB repeat units^32^. Further, the monomodal GPC trace (Fig. E3F) observed for this sample is consistent with a block-co-polymer structure. Moreover, minor resonances associated with 4HB-3HB and 3HB-4HB sequences in the NMR spectra (Fig. E2G) are also consistent with a block-like organization of the co-polymer, as previously reported^32^.

The relatively low T_*m*_ and T_*g*_ obtained for the P(3HB-co-4HB-co-6HC) co-polymer from *C. necator* H16 Δ*phaC1 hadA*-*phaC1437* and the ^1^H-NMR spectrum (Fig. E2K) are consistent with a random organization of the repeat units. Similarly, the absence of a melting point for the P(3HB-co-PheLA) co-polymer formed by *C. necator* H16 Δ*phaC1 hadA*-*phaC1437* (Fig. E4H) and the NMR spectra (Fig. E2M) are consistent with a random co-polymer structure the fully amorphous polymer appeared ductile at room temperature. These observations are consistent with the thermal properties and the resulting impacts on physical/mechanical properties and implications for manufacturing applications, as previously discussed elsewhere^16,17,26,38^.

For the P(3HB-co-PA) and P(3HB-co-5HM2F) co-polymers, the modest depression of the melting point (155^°^C and 159^°^C vs. 170^°^C, respectively) is consistent with previous observations for the incorporation of small amounts of aromatic co-monomer units in aliphatic polyesters^39^.

### 2.4 Dynamic Dependence of Bio-Polyester Composition

When considering the obtained polymers’ molecular weights in context with the time between induction and harvesting of the cultures, a dynamic correlation was apparent (see Tab. S1). Subject to further investigation, it was also noted that induction of expression led to reduced growth of the strain carrying the *pct540* gene as compared to the strain bearing the *hadA* gene (Fig. 2a,b). Determining the abundance of the heterologous enzymes in dependence of inducer concentration and time revealed that growth stagnation coincided with the onset of gene expression (see Extended Data 5); fructose consumption also came to a halt at that point. These effects are indicative of severe protein toxicity, a known problem of heterologous propionyl-CoA transferases like *pct540*^28^. This could explain both, the high content of non-canonical monomer, as well as block-co-polymer formation: Apparently stalling all metabolic activity, conversion of acetyl-CoA to 3-hydroxybutyryl-CoA must have also come to a halt, while exogenously supplied precursor was likely still available in excess. Hence, in the early stage (pre-expression) mostly natural P3HB synthesis was active, while in the final stage (*pct540* expressed), pure catalysis would have taken place, solely adding non-canonical monomers to the still-growing polymer chain. As a measure for a polymer’s heterogeneity (varying length/mass of differently composed chains), the comparatively high PDI of the PHA obtained with 4HB (∼3, see Tab. S1) supports this interpretation^40^. The same tendency is apparent when comparing the 3HP-content of the respective strains’ co-polymers, where the fraction of non-canonical monomer is an order of magnitude higher in the case of the *pct540*-strain as compared to the *hadA*-strain, further supporting this hypothesis.

**Figure 2.**
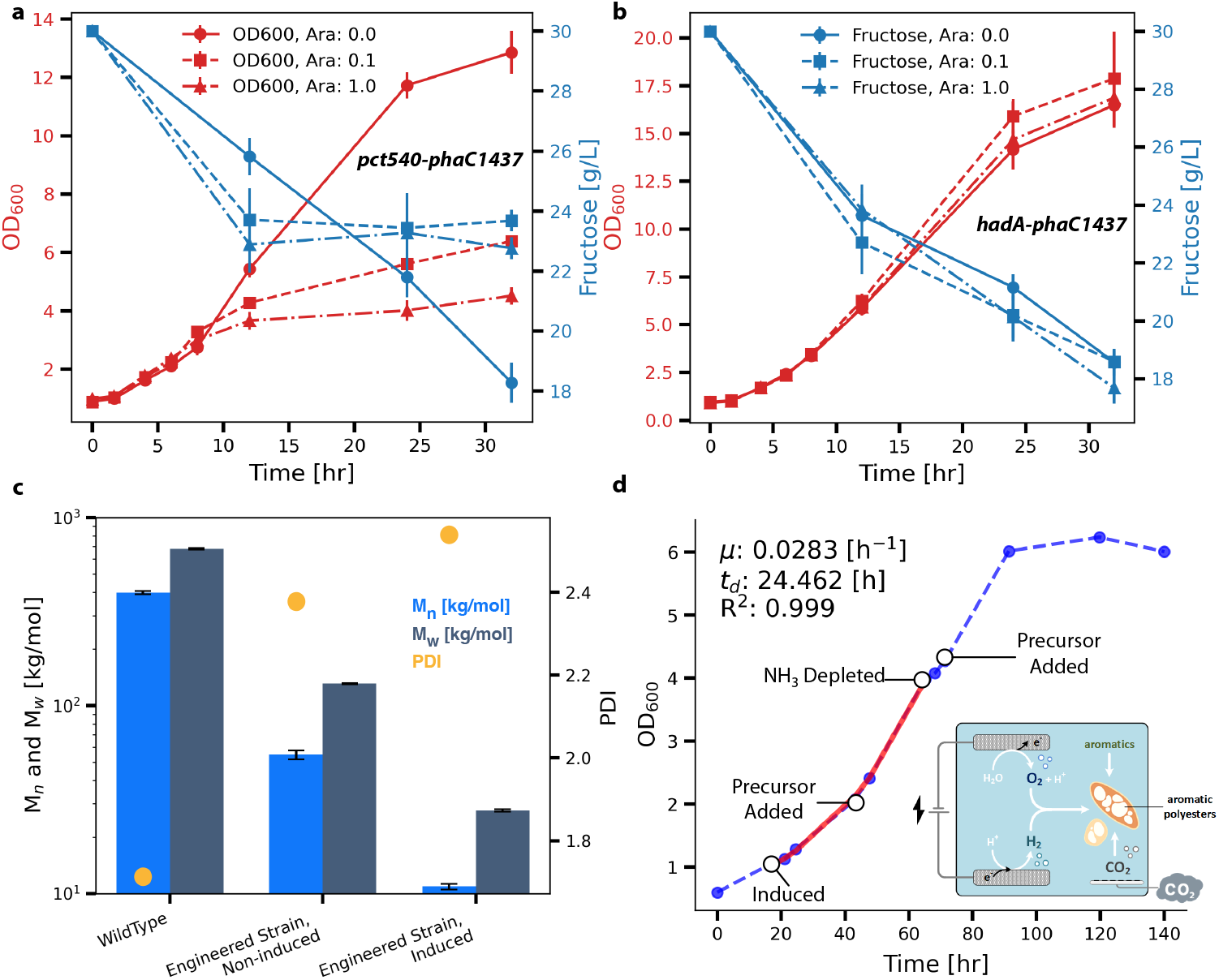
Typical cultivation profile of *C. necator* H16. **(a)** Δ*phaC1 pct540*-*phaC1437* and **(b)** Δ*phaC1 hadA*-*phaC1437* on minimal medium (MSM) with fructose as main carbon- and energy source. Growth is shown by means of inferred cell density (absorption of culture at 600 nm, red marker and lines) and fructose consumption (concentration in supernatant by HPLC, blue marker and lines) over time at different levels of induction (arabinose; circle = 0 g/L, square = 0.1 g/L, triangle = 1 g/L).**(c)**Comparison of molecular weight distribution (number average ‘M_*n*_’ and weight average ‘M_*w*_’) and polydispersity index (PDI) for PHA obtained from wildtype and engineered (Δ*phaC1 hadA*-*phaC1437*) *C. necator* H16, respectively. Molecular weights M_*n*_ and M_*w*_ are represented by blue bars, respectively, scaled on the left axis, while the PDI (M_*w*_/M_*n*_) is represented by an orange dot-marker scaled on the right axis.**(d)**Cultivation profile (growth measured as relative cell density) of a representative experiment with transgenic *C. necator* H16

The effects of enzyme expression levels on the polymers’ M_*w*_ and M_*n*_ were further investigated: compared to the bio-polyester produced by the wildtype of *C. necator* H16, the molecular weights of the PHAs formed by the non-induced transgenic strains were approx. five-fold lower (Fig. 2c). Induction of the engineered strain and over-expression of the heterologous genes (*hadA* and *phaC1437*) resulted in another five-fold reduction in molecular weights while the PDI also increased. This trend was prevalent among all analyzed polymers (without and with supplemented precursors, see Tab. S1), underscoring that enzyme expression level(s) and polymer chain length are apparently inversely correlated.

To determine the cause of this correlation, the expression levels of heterologous genes during induced and non-induced conditions were qualitatively compared. In the latter case, the proteins were not detectable (see Extended Data 5), albeit catalytic activity of the PHA synthase evidently existed. The hypothesis posits that PhaC1437 exhibits high activity, requiring minimal enzyme levels for PHA formation. This activity, when coupled with abundant catalyst levels, may deplete substrate locally, leading to premature termination of chain elongation, resulting in a reduced polymer molecular weight during induced conditions. These observations are consistent with a congruent phenomenon, observed when the gene order of the natural *phaCAB* operon was shuffled^41^.

### 2.5 Formation of Aromatic Bio-Polyesters under Autotrophic Conditions

As a lithoautotroph, *C. necator* H16 can assimilate carbon dioxide, deriving the reducing power for oxidative phosphorylation from molecular hydrogen. Aerobic gas fermentation, however, is costly and comprises serious safety concerns^42,43^. Therefore, innovative solutions have been proposed that allow the *in situ* formation of H_2_ and O_2_ via water splitting in a bio-electrochemical system (BES)^44–46^ where the electron donor is immediately consumed. To demonstrate the sustainable production of novel aromatic PHAs via microbial electrosynthesis (MES), *C. necator* H16 Δ*phaC1 hadA*-*phaC1437* was cultivated in a single-chamber BES with CO_2_ and electrical current as the only constant inputs, as shown schematically in Fig. 2d. In the membrane-less system, a constant current was applied, which led to water being split into oxygen at the anode and hydrogen at the cathode. CO_2_ was continuously purged into the reactor, providing a gas-mix for chemolithotrophic metabolism. The BES was inoculated from a pre-culture grown on a mixture of H_2_/O_2_/CO_2_ to facilitate adaptation to the electroautotrophic conditions. This resulted in immediate microbial growth in the BES with no lag-phase, as discernible from Fig. 2d. Preliminary experiments (see Fig. S6 & Tab. S4) suggested that the initial amount of nitrogen source in the BES was critical to achieving nutrient-limiting conditions and triggered PHA accumulation. Consequently, the concentration of ammonium in the medium was reduced and closely monitored in combination with growth to detect depletion of the nitrogen source. Polymer precursor (PheLA) was added in two steps to control toxicity, which appeared to be more severe during autotrophic metabolism. Further, the growth rate was lower under conditions of MES than on straight gas-mixture (BES: t_*D*_ ∼24 h, serum Δ*phaC1 hadA*-*phaC1437* in a BES. Induction of expression of the heterologous genes, addition of aromatic polymer precursor, and depletion of nitrogen source were performed/determined as indicated. The section of the growth curve used to estimate the rate of biomass formation and doubling time of the cells is shown in red. The BES is represented schematically in the insert on the bottom right: H_2_ and O_2_ evolve at the cathode and anode, respectively, serving as electron-carrier and acceptor, for autotrophic growth and metabolism from CO_2_ and electricity, enabling the formation of aromatic PHAs.

bottles: t_*D*_ ∼12 h; see Fig. 2d and Fig. S6, respectively), which can be explained with mass-transfer limitations in the BES (ambient pressure and limited agitation). Nevertheless, biomass accumulation exceeded the batch experiments due to continuous substrate replenishment: a maximum OD_600_ of 6.2, corresponding to a biomass concentration of 1.8 ∼g/L, was reached after 90 h. The culture was harvested after 140 h and extraction of polyesters yielded a polymeric resin, which was identified as P(3HB-co-PheLA) through ^1^H-NMR spectroscopy (see Extended Data 2). The fraction of non-canonical repeat units in the PHA obtained from the BES was lower than in the shakeflask experiments (3HB:PheLA ratio of 50:3 under autotrophic vs. 2:1 under heterotrophic condition), which could be explained by the limited energy available to metabolism under autotrophic conditions^47^, as well as the reduced concentrations of precursor (1/2 i.e., 5 mM PheLA) and inducer (1/10 i.e., 0.1 g/L arabinose). Nevertheless, this experiment demonstrates the feasibility of forming arylatic PHAs autotrophically and thereby the potential for the biosynthesis of aromatic polyesters from carbon dioxide.

### 2.6 Modelling the Biosynthesis of Novel Bio-Polyesters

To support the prospective *de novo* production of advanced PHAs, the feasibility of their biosynthesis from basic microbial feedstocks was assessed in an *in silico* analysis. Since for many promising precursors so far only co-polymers of various compositions have been reported, the scope of the analysis was limited to homo-polymers, even though this meant some structures were hypothetical. A reconstructed metabolic model of *C. necator* H16 was amended with biochemical pathways for formation of several aliphatic or aromatic hydroxy acids and the polymerization thereof. The entire set of metabolic networks is provided as SI 3; a schematic representation of the biosynthetic pathways for the aromatic compounds is provided in Fig. S8. Using elementary flux-mode analysis (EMA), the production capacity for the different (non-natural) polyesters in terms of maximum theoretical carbon yields was compared, considering fructose, glycerol, acetate, formate, and carbon dioxide + hydrogen as feedstocks. The results are visualized in Fig. 3, the numerical carbon yields can also be found in Tab. S5.

**Figure 3.**
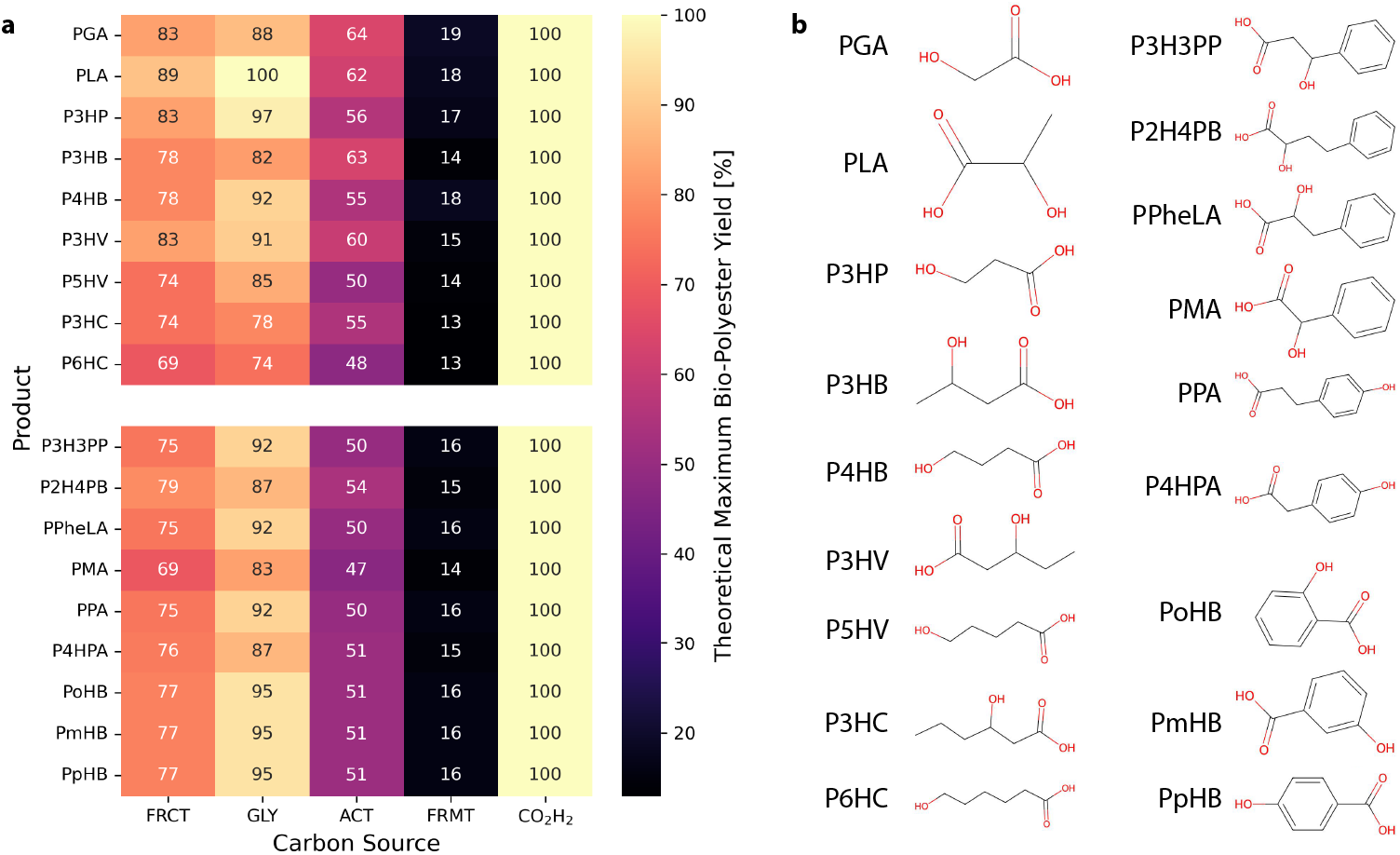
Theoretical maximum carbon-yields. **(a)** for a range of aliphatic, arylatic, and aromatic PHAs that could potentially be obtained from metabolism-derived precursors **(b)**, employing different carbon- and energy sources. Yields were calculated using Elementary Flux-Mode Analysis with pathways implemented in a metabolic model of *Cupriavidus necator* H16. Substrates: FRMT = formate, ACT = acetate, FRC = fructose, GLY = glycerol, CO_2_ H_2_ = carbon dioxide and hydrogen gas. Products: PGA = poly(glycolate) [poly(2-hydroxyacetate)], PLA = poly(lactate) [poly(2-hydroxypropionate)], P3HP = poly(3-hydroxypropionate), P3HB = poly(3-hydroxybutyrate), P4HB = poly(4-hydroxybutyrate), P3HV = poly(3-hydroxyvalerate), P5HV = poly(5-hydroxyvalerate), P3HC = poly(3-hydroxycaproate), P6HC = poly(6-hydroxycaproate), P3H2PP = poly(3-hydroxy-3-phenylpropanoate), P2H4PB = poly(2-hydroxy-4-phenylbutanoate), PPheLA = poly(phenyllactate) [poly(2-hydroxy-3-phenylpropanoate)], PMA = poly(mandelate) [poly(2-hydroxy-3-phenylacetate)], PPA = poly(phloretate) [poly(3-(4-hydroxyphenyl)propanoate)], P4HPA = poly(4-hydroxyphenylacetate), PoHB = poly(*ortho*-hydroxybenzoate), PmHB = poly(*meta*-hydroxybenzoate), PpHB = poly(*para*-hydroxybenzoate).

All pathways for biosynthesis of the considered aliphatic and aromatic PHAs appeared feasible within the context of the employed metabolic network. Hence, the respective non-natural bio-polyesters can theoretically be produced *de novo*, under the condition that (A) metabolic engineering efforts to enable the formation of the respective monomers are successful and (B) these are polymerizable via a PHA biosynthesis-analogous mechanism.

Certain feedstocks appeared superior over others in terms of theoretical maximum yield: carbon yields on formate were lowest, followed by acetate. Fructose and glycerol, as more reduced carbon sources, could deliver higher yields, only surpassed by carboxy-hydrogen. In the latter case, the near-100% carbon efficiency can be explained by the separation of carbon- and energy source into two chemical species (CO_2_ and H_2_). Their independent supply allows for a flexible ratio of carbon source and donor of reducing equivalents, such that steady-state conditions can result in feasible flux distributions. Effectively, this means that if sufficient energy is available, carbon dioxide can be recycled infinitely. The real-world consequence of this finding is imperative for process control, to optimize the gas-mix and/or implement effective recycling of unused substrate. The low yields on formate are owed to the low energy that is chemically bound by this combined carbon- and energy carrier. Much of the formate is simply converted to carbon dioxide to obtain reducing equivalents used to drive metabolism while little of the carbon is assimilated. An analogous outcome could be expected from a carboxy-hydrogen feed with fixed stoichiometry (and low hydrogen to carbon dioxide ratio). Nevertheless, formate can be electrochemically regenerated from carbon dioxide^49^. Therefore, if an economically viable and technically practical solution can be developed, a closed-loop recycling system could effectively mimic a scenario where overall carbon yields from formate should approach those on carboxy-hydrogen, without the drawbacks of gas-fermentation^42^.

### 2.7 Predicted Physicochemical and Mechanical Properties of Novel Bio-Polyesters

To allow an appraisal of the potential applications the novel aromatic bio-polyesters discovered in this study could have, their assessment of biocatalytic production was complemented with predictions of their material properties. For this purpose, the machine-learning tool “PolyID” was leveraged, which can estimate thermal, gas transport, and mechanical properties of polymers, outgoing from the structures of the monomers^48^. To this end, the density, glass transition and melting temperature, elastic modulus, as well as barrier to gases (N_2_, O_2_, CO_2_, and H_2_O) of phloretic, 4-hydroxyphenylacetic, *para*-hydroxybenzoic, and 5-hydroxymethyl-2-furanoic acid co-polymers with common aliphatic C2-C4 hydroxy acids were compared among each other as well as to other relevant polyesters (see SI 4). The results of the predictions are visualized in Fig. 4, where panel (a) comprises the basic aliphatic homo-polymers and the aromatic co-polymers that are derived from them, panel (b) provides an overview of certain aromatic homo-polymers for comparison, and panel (c) comprises aromatic polyesters that are currently produced at scale for commercial applications.

**Figure 4.**
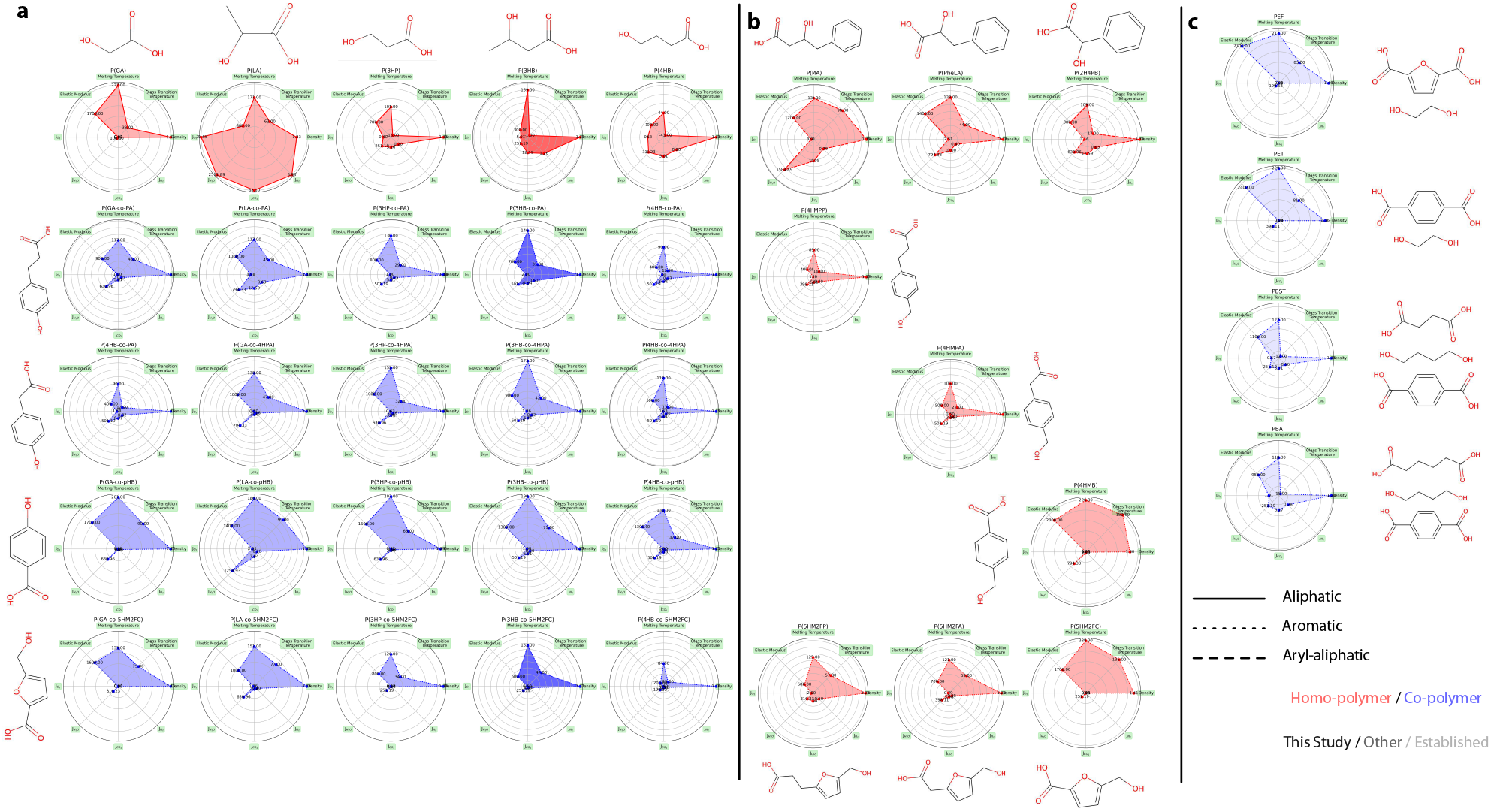
Comparison of selected aliphatic polyesters and co-polymers thereof with arylatic and aromatic hydroxy carboxylates in terms of relevant chemical, thermal, and mechanical properties. (as predicted by PolyID^48^), which are (in clockwise order): Melting Temperature [^°^C], Glass Transition Temperature [^°^C], Density [g/cm^3^], Permeability of N_2_ [Barrer], Permeability of CO_2_ [Barrer], Permeability of H_2_O [Barrer], Permeability of O_2_ [Barrer], Elastic Modulus [MPa]. Plots of homo-polymers are shown in red, co-polymers (equal fractions of repeat units) in blue. Plots in solid color pertain to polymers that have been bio-synthesized in this study (albeit with lower aromatics fraction). Molecular structures next to the plots are the monomers of the repeat units of the respective polymers. For the co-polymers in panel **(a)** the respective building-blocks are organized tabular on top as well as on the side of the plots. Panel **(b)** comprises several polymers that serve as proxies for the purpose of comparison to their structural analogs in panel **(a)**, specifically, these are: P4HMPP for PPA, P4HMPA for P4HPA, P4HMB for PpHB. Panel **(c)** comprises common commercial aromatic polyesters that are produced chemically. All source data can be found in SI 4. Abbreviations: PGA = poly(glycolate) [poly(2-hydroxyacetate)], PLA = poly(lactate) [poly(2-hydroxypropionate)], P3HP = poly(3-hydroxypropionate), P3HB = poly(3-hydroxybutyrate), P4HB = poly(4-hydroxybutyrate), P6HC = poly(6-hydroxycaproate), P2H4PB = poly(2-hydroxy-4-phenylbutanoate), PPheLA = poly(phenyllactate) [poly(2-hydroxy-3-phenylpropanoate)], PMA = poly(mandelate) [poly(2-hydroxy-3-phenylacetate)], PPA = poly(phloretate) [poly(3-(4-hydroxyphenyl)propanoate)], P4HPA = poly(4-hydroxyphenylacetate), PpHB = poly(*para*-hydroxybenzoate), P5H2MC = poly[5-(hydroxymethyl)furan-2-carboxylic], P5H2MA = poly[5-(hydroxymethyl)furan-2-acetic], P5H2MP= poly[5-(hydroxymethyl)furan-2-propionic], P4HMB= poly[4-(hydroxymethyl)benzoic], P4HMPA= poly[4-(hydroxymethyl)phenylacetic], P4HMPP= poly[4-(hydroxymethyl)phenylpropionic], PPA = poly(phloretate) [poly(3-(4-hydroxyphenyl)propanoate)], P4HPA = poly(4-hydroxyphenylacetate), PpHB = poly(*para*-hydroxybenzoate), PEF = poly(ethylene furan-2,5-dicarboxylate), PET = poly(ethylene terephthalate), PBST = poly(butylene succinate-co-terephthalate), PBAT = poly(butylene adipate-co-terephthalate).

When comparing the predicted properties of aliphatic-co-aromatic PHAs to those of the homo-polymers of each monomer, it appeared that the co-polymers typically exhibited weighted averages of the respective building blocks derived properties. Overall, a wide range of properties could be covered, supporting the concept of tailoring bio-polyesters specific to a desired application through tuning of the species and proportional content of the respective repeat units. Considering specifically the here obtained P(3HB-co-PA) and P(3HB-co-5HM2F) they may, depending on the fraction of co-monomers, exhibit unique material properties that are not currently known among existing PHAs. Most notably, these novel aromatic PHAs could approach or exceed the properties of commercial aromatic co-polyesters like PBST and PBAT. Other aromatic PHAs that emerged as potentially very similar to commercial aromatic co-polyesters such as PEF and PET, and resembled their properties, were the hypothetical co-polymers of 5HM2F and PA with glycolic acid. Further, given how closely the eight considered material properties of the novel aromatic PHAs and existing commercial aromatic polyesters align, it is plausible that this extends to other properties (e.g., tensile strength, elongation at break) not currently included in this analysis.

## 3 Conclusion

Through the co-polymerization of bio-derived aliphatic, arylatic, and aromatic hydroxycarbonates with 3-hydroxybutyrate we demonstrated the biosynthesis of novel polyhydroxyalkanoates. Most significantly, we report the first-ever incorporation of aromatic rings in the backbone of biological polyesters. The obtained polymers bear structural resemblance to synthetic polyesters such as PBST and PBAT as well as PET and PEF. Prediction of their physicochemical and mechanical properties corroborated the analogies among the novel aromatic PHAs and existing polyesters, substantiating the potential to replace incumbent plastics with sustainable alternatives. Eventually, even high-performance “polyhydroxyarylates” with liquid-crystal polymer characteristics that resemble polyarylates^21,50^ may become bio-available, shaping a new class of bio-polyesters. Additionally, we obtained arylatic co-polymers bearing phenyl groups on a side chain, which significantly altered the materials’ properties. These polyesters could become biodegradable alternatives to indispensable but non-recyclable bulk plastics such as polystyrene^19,20^. Moreover, our system enabled co-polymerization of aliphatic short-chain *ω*-hydroxy acids. Depending on the fraction, this can balance the brittleness of natural P3HB, while maintaining adequate processability^15,16^. Valued for their biocompatibility and resorbability, viscoelastic, homo-polymers thereof, such as polycaprolactone (PCL), are already used in medical applications^51^.

Characterization of the obtained polymers revealed an inverse correlation between the expression level of the PHA synthase and the obtained polymers’ molecular weights. This finding underscores the importance of balancing enzyme catalytic rates with substrate availability to reach a critical molecular weight that leads to chain entanglement (which occurs approx. at two-fold of the entanglement molecular weight^52^, for PHB about 14 kDa^53^) and this a useful material. Optimizing the biocatalytic formation of advanced polyesters, especially from aromatic hydroxycarbonates, will likely also require the discovery of novel PHA synthases. This offers the opportunity to expand and tailor the enzyme’s functionality to the precise composition required for a polymer with the desired material properties.

While PHAs are inherently biodegradable, the greatest advantage of polyester is their deconstructability: recovery of polymer building blocks from the breakdown of spent plastics into their monomers can avoid excessive and resource-intense downcycling, enabling closed-loop re- and upcycling^54,55^. In addition, the precursors for the here-obtained PHAs are also available from other waste streams (e.g., VFAs from anaerobic digestion) as well as non-food carbon sources. In particular the aromatics can be directly derived from wood-waste: phloretic acid from lignin^56,57^ and 5-hydroxymethyl-2-furanoic acid via 5-(hydroxymethyl)furfural (HMF) from lignocellulosic^58^ as well as lignin-free biomass^59^. Through metabolic modeling, we showed that the here-considered hydroxycarbonic acids could also be *de novo* biosynthesized from single-carbon compounds such as carbon dioxide via metabolic engineering. For a similar purpose, we demonstrated autotrophic cultivation of the transgenic *C. necator* strain in a BES and obtained arylatic PHAs. The electro-fermentation approach could defossilize a large part of the polymer industry and integrate this sector into a circular economy^4^; if extended to all bulk plastics currently produced, nearly 5% of global GHG emissions could potentially be mitigated^6^. While extensive metabolic and process engineering will be required to make the one-step bioproduction of advanced bioplastics at scale a reality, recent developments in synthetic biology and advancement of genetic tools for *C. necator* have brought this strategy for valorization of CO_2_ within reach^47^.

## 4 Materials and Methods

### 4.1 Strains and Media

Rich Broth (RB) was composed of Nutrient Broth (16 g/L), Yeast Extract (10 g/L) and ammonium sulfate (5 g/L). When applicable, kanamycin (kan, 300 mg/L for *C. necator* and 100 mg/L for *E. coli*) and/or diaminopimelate (DAP, 300 µM / 100 mg/L) were added.

Modified Mineral Salts Medium (MSM) was composed as described before^60^; in brief, it contained 6.4 g/L Na_2_HPO_4_, 4.8 g/L KH_2_PO_4_, 1 g/L NH_4_Cl, MgSO_4_ 0.36 g/L, CaCl_2_ ×2H_2_O 0.088 g/L and 1 mL/L of 1000 ×trace elements (0.2 g/L CoCl_2_ ×6H_2_O, 0.01 g/L CuSO_4_ ×5H_2_O, 0.15 g/L FeSO_4_ ×7H_2_O, 0.06 g/L NH_4_Fe(III) citrate, 0.3 g/L H_3_BO_4_, 0.035 g/L MnCl_2_ ×4H_2_O, 0.035 g/L (NH_4_)6Mo_7_O_24_× 4H_2_O, 0.031 g/L NiSO_4_ ×7H_2_O, 0.1 g/L ZnSO_4_ ×7H_2_O). The medium was supplemented with 20 g/L fructose as carbon- and energy source and 300 mg/L kanamycin (to ensure plasmid maintenance) when applicable.

Analogously to previously described toxicity tests^61^, gradient-agar was produced by sequentially pouring two layers of MSM agar of varying thickness with 2 ppm Nile red, with or without polymer precursor added, on top of each other in square culture/Petri dishes. This resulted in a horizontal gradient through the linear difference in vertical thickness of each individual layer. The plates were used immediately to avoid lateral equilibration of the concentration gradient, spreading 200 µL of *C. necator* H16 culture with an effective OD_600_ of 0.2, followed by incubation at 30^°^C for 72 h.

For (electro)autotrophic cultivation, a modified minimal medium that contained no chlorine salts (to avoid electrocatalytic formation of Cl_2_) was adapted from literature^62^. Specifically, the “electro-medium” contained 4.52 g/L Na_2_HPO_4_, 5.17 g/L NaH_2_PO_4_ ×2H_2_O, 0.17 g/L K_2_SO_4_, 0.097 g/L CaSO_4_ ×2H_2_O, 0.8 g/L MgSO_4_ ×7H_2_O, 0.94 g/L (NH_4_)_2_SO_4_ and 350 µL/L of 2857 ×trace elements (10 g/L FeSO_4_ ×7H_2_O, 2.4 g/L MnSO_4_ ×H_2_O, 2.4 g/L ZnSO_4_× 7H_2_O, 0.48 g/L CuSO_4_ ×5H_2_O, 1.8 g/L Na_2_MoO_4_× 2H_2_O, 1.5 g/L NiSO_4_ ×6H_2_O, 0.04 g/L Co(NO_3_)_2_ H_2_O in 0.1 M HCl). The medium was supplemented with 100 mg/L kanamycin to ensure plasmid maintenance. For production of PHA under autotrophic conditions, the initial concentration of (NH_4_)_2_SO_4_ was reduced to 5 mM to achieve nitrogen-limiting conditions.

*E. coli* WM3064, a B2155 derivative and DAP-auxotrophic RP4-mobilizing λ *pir* cell-line 60, was a gift from the Spormann Laboratory (Stanford University). WM3064 was routinely cultivated on LB (solid or liquid, at 30^°^C) supplemented with DAP (300 µM) and kanamycin (50 µg/mL) as applicable. *C. necator* H16 and *C. necator* H16 Δ*phaC1* were gifts from the Silver Laboratory (Harvard Medical School)^63^ and routinely cultivated on RB or MSM (solid or liquid, at 30^°^C). A comprehensive list of the strains used and constructed in this study can be found in Tab. 2.

**Table 2.**
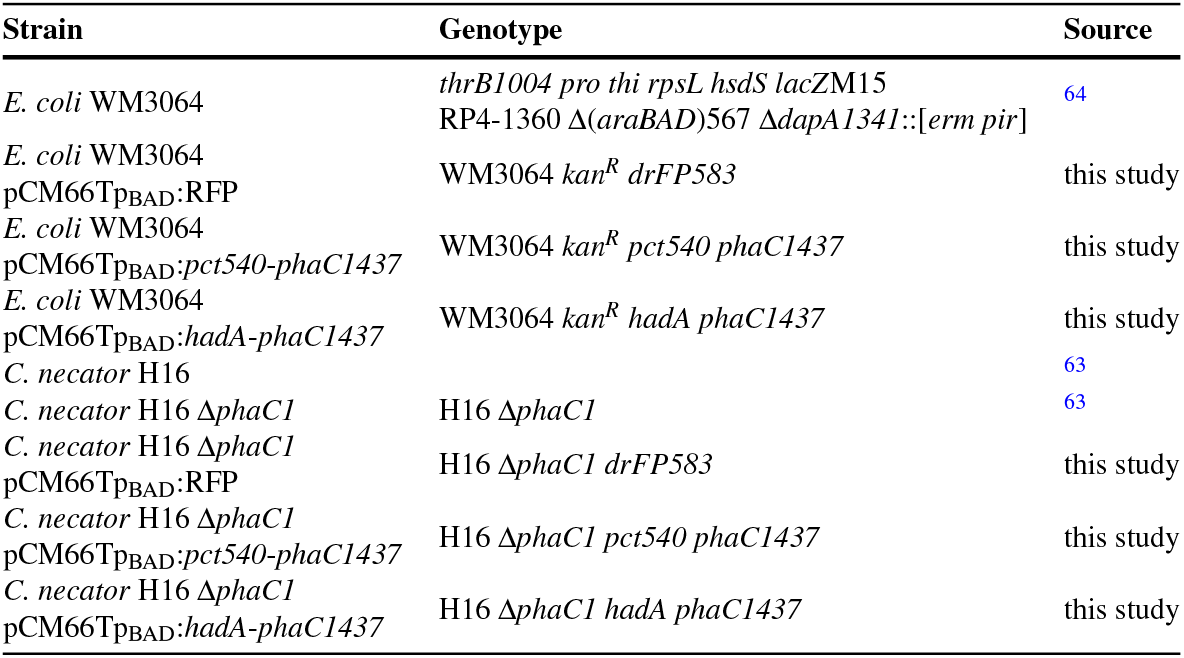
Microbes used in this study and identity of genetically engineered mutant strains.

### 4.2 Vector Construction

The RK2/RP4 oriV (IncP) plasmid pCM66T was obtained from AddGene. pBBR1MCSp_BAD_:RFP, a derivate of pBBR1MCS with a red fluorescent protein (*drFP583*) under control of the araBAD promoter, was a gift from the Silver Laboratory (Harvard Medical School). pCM66Tp_BAD_:RFP was constructed by cloning the P_BAD_-RFP cassette into pCM66T using NEBuilder, replacing the polylinker and its regulatory elements. The protein sequences of Q9L3F7_CLOPR_ (*pct*), Q188I3_PEPD6_ (*hadA*) and B9W0T0_9PSED_ (*phaC*) were derived from UniProt, implementing the previously described mutations (V193A for *pct540* and E130D, S325T, S477G, Q481K for *phaC1437*) as applicable. The sequences were fused with a C-terminal tripleglycine spacer and tetracysteine-(Lumio)-tag and codon optimized for expression in *C. necator* with GeneArt (Invitrogen). The genes were arranged in a single operon under control of the *araBAD* promoter in combination with the strong T7 ribosomal binding site and a T7Te – rrnB T1 double-terminator. The plasmids pCM66Tp_BAD_:*pct540*-*phaC1437* and pCM66Tp_BAD_:*hadA*-*phaC1437* were constructed by GenScript Biotech, cloning the synthetic operons containing *pct540 & phaC1437* and *hadA & phaC1437*, respectively, into pCM66Tp_BAD_:RFP, replacing the RFP (see Fig 1c and SI 2 for annotated sequences of plasmids).

### 4.3 Conjugation

Plasmid vectors were introduced into *C. necator* by bacterial conjugation, using the *E. coli* donor-strain WM3064, which had been transformed with the plasmid vectors pCM66Tp_BAD_:*pct540*-*phaC1437* and pCM66Tp_BAD_:*hadA*-*phaC1437* using the Mix & Go! *E. coli* Transformation Kit (Zymo).

Conjugation was performed as follows: the recipient strain (*C. necator* H16 Δ*phaC1*) was incubated at 30^°^C on RB plates overnight; simultaneously the donor strains (WM3064) carrying the plasmid vectors were incubated at 30^°^C on LB plates with kan and DAP. From the agar plates, liquid cultures of the donor strains were inoculated (LB with kan and DAP) and incubated overnight with shaking at 30^°^C. Likewise, RB was inoculated with the recipient strain and incubated overnight with shaking at 30^°^C. On the next day, 3 mL of LB with DAP (but no antibiotics) per donor-strain were each inoculated with 6 µL of overnight culture and incubated with shaking at 37^°^C. Similarly, a matching number of recipient strain cultures were inoculated combining 20 µL of overnight culture and 10 mL of RB each, which were incubated with shaking at 30^°^C. After 4 h the cultures were combined using 3 mL of the *E. coli* and 10 mL of the *C. necator* culture. Cells were collected by centrifugation (4816 × g for 10 min) and the supernatant was discarded. The cell pellet was re-suspended in 200 µL of RB with DAP and a “blob” of each cell mixture was pipetted on an RB plate with DAP. The agar plates were incubated at 30^°^C overnight with the place facing up. On the next day, a single scoop of biomass was collected with an inoculation loop and suspended into 500 µL of 25% glycerol. The suspension was diluted 1:10 and 1:100 and all three concentrations were plated on separate RB plates with kan. The plates were incubated at 30^°^C; colonies that appeared after about two days were picked and isolated on separate RB plates with kan for propagation and screening.

### 4.4 Cultivation of Microbes

#### 4.4.1 Cultivation of C. necator in Shake- and Serum-Flasks

Liquid cultures under heterotrophic conditions were maintained in 500 mL vented and baffled shakeflasks (WHEATON ® Erlenmeyer Flasks with DuoCap®, DWK Life Sciences), incubated at 30^°^C with shaking at 180 rpm on an innova 2300 platform shaker (New Brunswick Scientific). For feeding experiments, precultures of the engineered *C. necator* strains were inoculated in liquid RB + kan (50 mL) from solid RB + kan and grown overnight. In the morning the medium was exchanged for MSM + kan, washing and diluting the cultures 1:2 (in 100 mL). In the afternoon the seed culture was washed again with MSM and again diluted 1:2 (in 200 mL) with MSM + kan + polymer precursor. In the evening the culture was induced with 1 g/L arabinose unless indicated otherwise. Growth was monitored by means of OD (absorbance at 600 nm), accompanied by collection of supernatant samples, as applicable. The cultures were harvested after approx. 48 h or when no further increase in biomass density was observed.

Precultures for the bio-electrochemical system were two-staged: 125 mL serum bottles (25 mL liquid medium, 100 mL gas headspace) were filled with 25 mL “electro-medium”, sealed with butyl rubber stoppers, and secured with crimp caps. The cultures were initiated on 1 g/L fructose, inoculating the medium with the engineered *C. necator* from solid RB + kan and incubated at 30^°^C with shaking at 200 rpm overnight. Subsequently, the cultures were adapted to autotrophic conditions with H_2_/CO_2_/O_2_: the initial gas-phase was H_2_/CO_2_ (80%/20%) at 17 psi, which was further pressurized with O_2_ (100%) to 22 psi, resulting in a final gas composition of 64:16:20 (H_2_/CO_2_/O_2_) [v/v/v]. After three days of incubation, autotrophic growth was observed; the culture reached a maximum OD_600_ of 3.5± 0.1 (Fig. S6). For inoculation of the BES, 35 mL of exponentially growing culture was harvested at an OD_600_ of ∼0.7 (by centrifugation at 4000 ×g for 6 min) and re-suspended in 25 mL of fresh medium, to be transferred to the bioreactor.

#### 4.4.2 Cultivation of C. necator in the Bio-Electrochemical System

The bio-electrochemical system was a custom (500 mL) glass vessel (Adams & Chittenden Scientific). The reactor was operated membraneless as a three-electrode setup, magnetically stirred at 300 rpm. The cathode was a Nickel-Molybdenum alloy on graphite support (total surface area 50 cm^2^), which has previously been characterized and was found to evolve H_2_ at 100% selectivity under biologically relevant conditions^65,66^. The anode was platinized titanium mesh (PLANODE1X4, TWL) and an Ag/AgCl reference electrode (NaCl saturated; RE-5B, BASi®), both of which were inserted via a rubber stopper side port. The electrochemical reactor was controlled by applying a constant current of 100 mA using a multichannel potentiostat (VMP3; BioLogic Science Instruments, EC-Lab 11.21), providing a constant electron flow and therefore constant rate of H_2_ and O_2_ (GC analysis of the reactor’s headspace in abiotic pre-tests confirmed that H_2_ and O_2_ were the sole gaseous products). A constant flow rate of CO_2_ was ensured at 1 mL/min via a mass flow controller (EL-Flow F 100D, Bronkhorst®).

Before inoculation, the reactor was filled with 300 mL of medium, and the BES was operated for at least one hour under abiotic conditions to saturate the medium with the gases. Using 25 mL of concentrated cell suspension from exponentially growing, autotrophic cultures as inoculum, a starting OD_600_ between 0.5 and 0.6 (and a final liquid volume of 325 mL) was achieved. Preliminary tests showed that under autotrophic growth conditions, the accumulation of PHA required tight and precise limitation of the nitrogen source. Therefore, the initial concentration of ammonium salt in the BES was reduced to 5 mM and consumption was monitored qualitatively (using “EasyStrips” Ammonia Test Strips, Tetra® GmbH with a sensitivity *<* 0.5 mg/L) throughout the experiment in combination with growth and pH by drawing samples manually. After 24 h of stable growth, the culture was induced with arabinose (0.1 g/L final conc.). After another ∼ 24 h from induction, the first dose of polymer precursor was added (2.5 mM final conc.), the second dose (additional 2.5 mM to reach a total of 5 mM) when the detection limit of ammonium was undershot, i.e. when the nitrogen source was depleted. The experiment was terminated when no further growth was observed. Cells were harvested via centrifugation followed by polymer extraction. Pictures of the setup (for illustration purposes) are provided in SI 1 (Fig. S7).

### 4.5 Extraction of PHAs

Liquid cultures of *C. necator* were harvested by centrifugation and the cell pellet was freeze-dried. After determining the dry cell weight (CDW), PHAs were extracted by lysis of the cell pellet (wet or dry) with 10% sodium hypochlorite solution (Honeywell Fluka™) using approx. 0.2 L/g_CDW_. The pellet was completely suspended and incubated at room temperature for 20 min with intermittent mixing. The suspension was diluted with water 1:2 and centrifuged at 4816 ×g for 20 min. As the only remaining solids, the PHAs were washed twice with water and once with methanol, repeating the centrifugation step. The dried polymer was weighed to determine the product yield, dissolved in chloroform, filtered through a 0.2 µm PTFE “Titan 3™” (Thermo Scientific™) syringe filter, and dried for analysis.

### 4.6 Protein Extraction and Detection

Culture derived from distinct time-points of batch cultivation (exponential-phase / stationary-phase) was collected (sample volume [mL] ∼ 10 / OD_600_) and cells were pelleted by centrifugation (4816×g at 4^°^C for 10 min). The pellet was washed with purified water and stored as cell paste at -20^°^C for later processing. For extraction of proteins, CelLytic™B (Sigma) was used as per manufacturer’s directions (approx. 1 mL per cells from 10 mL culture at an OD of 1), in combination with Protease Inhibitor Cocktail (Sigma-Aldrich). The mixture was vortexed for 2 min to lyse the cells and extract the soluble protein. Centrifugation (4816×g for 10 min) pelleted the cell debris; the supernatant, which contained the soluble proteins, was separated. Total protein concentration was determined using the BCA Protein Assay Kit (Pierce™). Using the Lumio™ Green Detection Kit (ThermoFisher) as per manufacturer’s directions, 10 µg crude protein extract of each sample were prepared for gel electrophoresis. Size-separation was performed on a Bolt™ 4 - 12% Bis-Tris Plus Gel (ThermoFisher), run at 150 V for approx. 40 min with Bolt™ MES SDS Running Buffer (ThermoFisher). The marker was BenchMark™ Fluorescent Protein Standard (ThermoFisher). A Gel-Doc (BioRad) was used to visualize fluorescent-conjugated proteins. For visualization of all proteins, the gels were re-stained with One-Step Lumitein™ Protein Gel Stain (Biotium) as per the manufacturer’s directions and imaged again as before.

### 4.7 Analytics

#### 4.7.1 Determination of OD and Cell Dry-Weight Correlation

Microbial growth was characterized by measuring the optical density at 600 nm (OD_600_) with a DR2800™ Spectrophotometer (HACH) for shakeflask cultures and an Ultrospec™ 2100 pro (Amersham BioSciences) in case of MES using the BES.

A correlation between OD_600_ and biomass (BM) concentration was determined gravimetrically (see Tab. S2) from batch shakeflask cultivations with the wildtype (five samples) and engineered strains (15 samples) of *C. necator* H16. Shakeflask cultures of 50 mL and 100 mL volume with different cell densities were harvested via centrifugation and vacuum dried. The quotient of OD_600_/BM (cell dry-weight in mg) averaged from all samples was 0.31±0.05, such that the correlation is: OD_600_×0.3 ∼ BM [g/L].

#### 4.7.2 High-performance liquid chromatography (HPLC)

Quantification of fructose in the culture-broth was based on a previously published HPLC method for the detection of organic acids^67^. In short, the procedure was as follows: Samples (1 mL) were filtered (PVDF or PES syringe filters, 0.2 µm pore-size) and diluted 1:100 into HPLC sampling vials. Analysis of 50 µL sample volume was performed on a 1260 Infinity HPLC system (Agilent), using an Aminex HPX87H column (BioRad) with 5 mM H2SO4 as the eluent, at a flow rate of 0.7 mL/min. Fructose was identified and quantified by comparison to standards (3 g/L, 1.5 g/L, 0.6 g/L, 0.3 g/L, 0.15 g/L, 0.03 g/L), according to retention time (8.8 min) using a refractive index detector (35^°^C).

#### 4.7.3 Nuclear Magnetic Resonance (NMR) Spectroscopy

NMR samples were prepared as previously reported^68^. In short, a few mg of polymer were dissolved in deuterated chloroform and ^1^H-NMR as well as ^13^C-NMR spectra were recorded at 25^°^C on a Unity INOVA™ 500 NMR Spectrometer (Varian Medical Systems) with chemical shifts referenced in ppm relative to tetramethylsilane.

#### 4.7.4 Gel Permeation Chromatography (GPC)

Gel Permeation Chromatography (GPC) TSKgel SuperHZM-H column (Tosoh) connected in series with a DAWN multiangle laser-light scattering (MALLS) detector and an Optilab T-rEX differential refractometer (Wyatt Technology). Polystyrene calibrated (from M_*p*_ = 500 - 275,000 g/mol) molecular weights were determined using a GPCmax autosampler at 25^°^C at a flow rate of 1 mL/min.

#### 4.7.5 Differential Scanning Calorimetry (DSC)

Differential Scanning Calorimetry (DSC) was conducted on a Q2500 (TA Instruments) using a heating and cooling rate of 10^°^C/min and a nitrogen flow rate of 50 mL/min. Heat flow (mW) was recorded relative to an empty reference pan. Glass transition temperatures and melting points were determined on the second heating scan at a heating rate of 10^°^C/min. The crystallization enthalpy was determined from the area of the crystallization peak of the cooling curve.

### 4.8 Metabolic Modelling

Based on previously established metabolic networks of *C. necator* for elementary flux-mode analysis^69,70^, a refined and fully compartmentalized model was fundamentally re-constructed. Expansions were made to describe C1- and energy-metabolism more precisely^34,71^. The model was amended with additional pathways for carbon assimilation and product formation, deducted from metabolic databases such as KEGG^72,73^. Reaction thermodynamics of the heterologous pathways were verified using the eQuilibrator tool for estimation of Δ_*r*_G’^*m*^, the Gibbs free energy of a reaction at relevant metabolite concentrations^74^.

Elementary flux modes were calculated in MATLAB® (MathWorks®), using ‘FluxModeCalculator’^75^, and evaluated as described previously^76^. Balances were established around boundary reactions, allowing carbon yields [C-mol/C-mol] to be determined for all considered products.

#### 4.8.1 Prediction of Polymer Properties

The density, elastic modulus, melting and glass transition temperature, as well as permeability for gases (N_2_, CO_2_, H_2_O, O_2_) of different polyesters was predicted using the web-interface of PolyID^48^. Specifically, the SMILES codes of the polymeric compounds were generated from the structures of their monomers, allowing their processing and individual estimation of the material properties.

## Supporting information

Inventory of Supplementary Information

## Abbreviations

3HP: 3-hydroxypropionic acid
4HB: 4-hydroxybutyric acid
5HM2F: 5-hydroxymethyl-2-furanoic acid
6HC: 6-hydroxycaproic acid
BES: bio-electrochemical system
BM: biomass
CDW: cell dry-weight
DAP: diaminopimelic acid
DSC: differential scanning calorimetry
EMA: elementary flux-mode analysis
GPC: gel-permeation chromatography
HA: hydroxy acid
MA: mandelic acid
MES: microbial electrosynthesis
Mn: number-average molecular weight
MSM: mineral salts medium
MW: weight-average molecular weight
NMR: nuclear magnetic resonance
OD: optical density
PA: phloretic acid
PBAT: polybutylene adipate-co-terephthalate
PBST: polybutylene succinate-co-terephthalate
PDI: polydispersity index
PEF: polyethylene furanoate
PET: polyethylene terephthalate
PHA: polyhydroxyalkanoate
PHB: polyhydroxybutyrate
PheLA: phenyllactic acid
PLA: polylactic acid
RB: rich broth
rpm: rounds per minute
VFA: volatile fatty acid2

## 5 Data Availability

The authors confirm that the data supporting the findings of this study are available within the article and its supplementary information. The extent of the data provided is sufficient for replication of the study. Any other data that may be pertinent are available on request from the corresponding author. All code used for aggregation and evaluation of the data as well as generation of the figures is available at GitHub.

## 6 Extended Data

Extended Data are provided as separate figures accompanying the manuscript and contain additional experimental results that support key findings presented in the main text. A full description of each piece of Extended Data is provided in the accompanying “Inventory of Supporting Information”.

## 7 Acknowledgements

This study was supported by a Stanford Natural Gas Initiative (NGI) program grant (SPO #139138), as well as NASA grant/cooperative agreement award number NNX17AJ31G. Any opinions, findings, and conclusions or recommendations expressed in this material are those of the author and do not necessarily reflect the views of the National Aeronautics and Space Administration (NASA). Part of this work was performed at the Stanford Nano Shared Facilities (SNSF), supported by the National Science Foundation (NSF) under award number ECCS-2026822.

We would like to dedicate this work to Vince, who was instrumental in the discovery of the described aromatic polyesters and without whose perseverance and resilience throughout the years we could not have proven the core hypothesis of this study.

## 8 Author Contributions

NJHA conceived the study, designed, and conducted the experiments, and composed the manuscript. VEP and FK performed analytics (NMR, GPC, DSC, and HPLC, respectively) with help from MZ and SNN who also aided in strain construction. Further, FK designed and conducted experiments in the bio-electrochemical system. For the revision of the manuscript SC performed growth-challenge assays and AJB helped to improve the presentation of the data. RMW and CSC supported and advised the project. All authors have read and approved the final manuscript.

