## Supplementary material for "Novel Aromatic Polyhydroxyalkanoates from Engineered *Cupriavidus necator* H16: Expanding Bio-Polyester Synthesis from Renewable Feedstocks": Inventory of Supplementary Information

**Inventory of Extended Data & Supporting Information** for "Novel Aromatic Polyhydroxyalkanoates from Engineered *Cupriavidus necator* H16: Expanding Bio-Polyester Synthesis from Renewable Feedstocks"

1. Extended Data:

**Fig. E1 A-V:** Growth challenge assays of *C. necator* H16 on gradient-agar plates containing polymer precursors and qualitative visualization of PHA content by means of fluorescence staining.

**Fig. E2 A-S:** Nuclear Magnetic Resonance (NMR) spectra of certain aromatic polymer precursors and bio-polyesters formed by wild-type and engineered *C. necator* H16 – $^{1}$H-NMRs for all compounds and $^{13}$C-NMRs in certain instances.

**Fig. E3 A-N:** Gel Permeation Chromatography (GPC) traces of bio-polyesters formed by wild-type and engineered *C. necator* H16 co-fed with hydroxy carbonic acids.

**Fig. E4 A-N:** Dynamic Scanning Calorimetry (DSC) plots of bio-polyesters formed by wild-type and engineered *C. necator* H16 co-fed with hydroxy carbonic acids.

**Fig. E5A & B:** Protein gels visualizing the expression levels of the heterologous enzymes by the engineered *C. necator* H16 at different time-points post induction and at different inducer concentrations.

2. Supporting Information 1:
    a. Extended discussion correlating the growth-experiments and expression of the heterologous enzymes under different conditions (pertaining to replication of figures S1 & S2 and S3 & S4, which are replications of figures 3A&B and E5A&B, respectively).
    b. Evaluation of NMR spectra and identification of co-polymers by means of resonant frequency of nuclei in the magnetic field relative to the standard (chemical shifts).
    c. Figure S5 comprises the composition of the full spectrum of biologically generated polyhydroxyalkanoates as per NMR analysis in terms of fraction of repeat-units.
    d. Tables S1 contains date on the composition and physical properties of the biologically generated polyhydroxyalkanoates as per GPC and DSC analyses.
    e. Figure S6 shows preliminary growth-trials of *C. necator* H16 on $CO_2/O_2/H_2$ for elucidation of optimum-limiting concentration of nitrogen-source.
    f. Table S2 contains key-data for autotrophic growth of *C. necator* H16 in serum-bottles.
    g. Figure S7 shows the setup of the bio-electrochemical system (BES) for autotrophic cultivation of *C. necator* H16 (left: abiotic system before inoculation, right: operational bioreactor with fully developed culture).
    h. Figure S8 is an illustration of biochemical pathways for production of aromatic PHAs (natural as well as synthetic routes of hypothetical nature).
    i. Table S3 comprises numerical values of maximum theoretical carbon-yields of polyesters as per metabolic network modeling by Elementary Flux-Mode Analysis.

3. Supporting Information 2: annotated vector maps of genetic constructs (pCM66T-pBAD:pct540-phaC1437, pCM66T-pBAD:hadA-phaC1437) used to establish the biochemical functions that enabled polymerization of the non-natural hydroxy acids.

4. Supporting Information 3: metabolic models comprising central metabolism of *Cupriavidus necator* H16 and synthetic biochemical pathways for formation of non-natural bio-polyesters as well as a compilation of the aggregated results (maximum theoretical carbon-yields). This data supports Figure 3.

5. Supporting Information 4: dataset of predicted material properties for novel aromatic bio-polyesters and related reference materials. This data supports Figure 4 and includes both experimentally obtained and hypothetical co-polymers, enabling comparison to commercial synthetic polyesters.
