## Extended Data 1 for "Novel Aromatic Polyhydroxyalkanoates from Engineered *Cupriavidus necator* H16: Expanding Bio-Polyester Synthesis from Renewable Feedstocks"

- Screening of *C. necator* H16 for precursor-toxicity on gradient-agar plates (minimal medium)
- Visualization of biomass formation under ambient light and UV transillumination
- Detection of bio-polyesters by fluorescence staining (2 ppm of Nile red)

Compounds evaluated over a range of 0 – 30 mM:

**5-hydroxymethyl-2-furanoic acid**
**3-(4-hydroxyphenyl)propionic (phloretic) acid**
**2-hydroxy-2-phenylacetic (mandelic) acid**
**2-hydroxy-3-phenylpropionic (phenyllactic) acid**
**2-hydroxy-4-phenylbutyric acid**
**6-hydroxycaproic acid**
**4-hydroxybutyric acid**
**3-hydroxypropionic acid**
**2-hydroxypropionic (lactic) acid**
**2-hydroxyacetic (glycolic) acid**

**5-hydroxymethyl-2-furanoic acid**

A

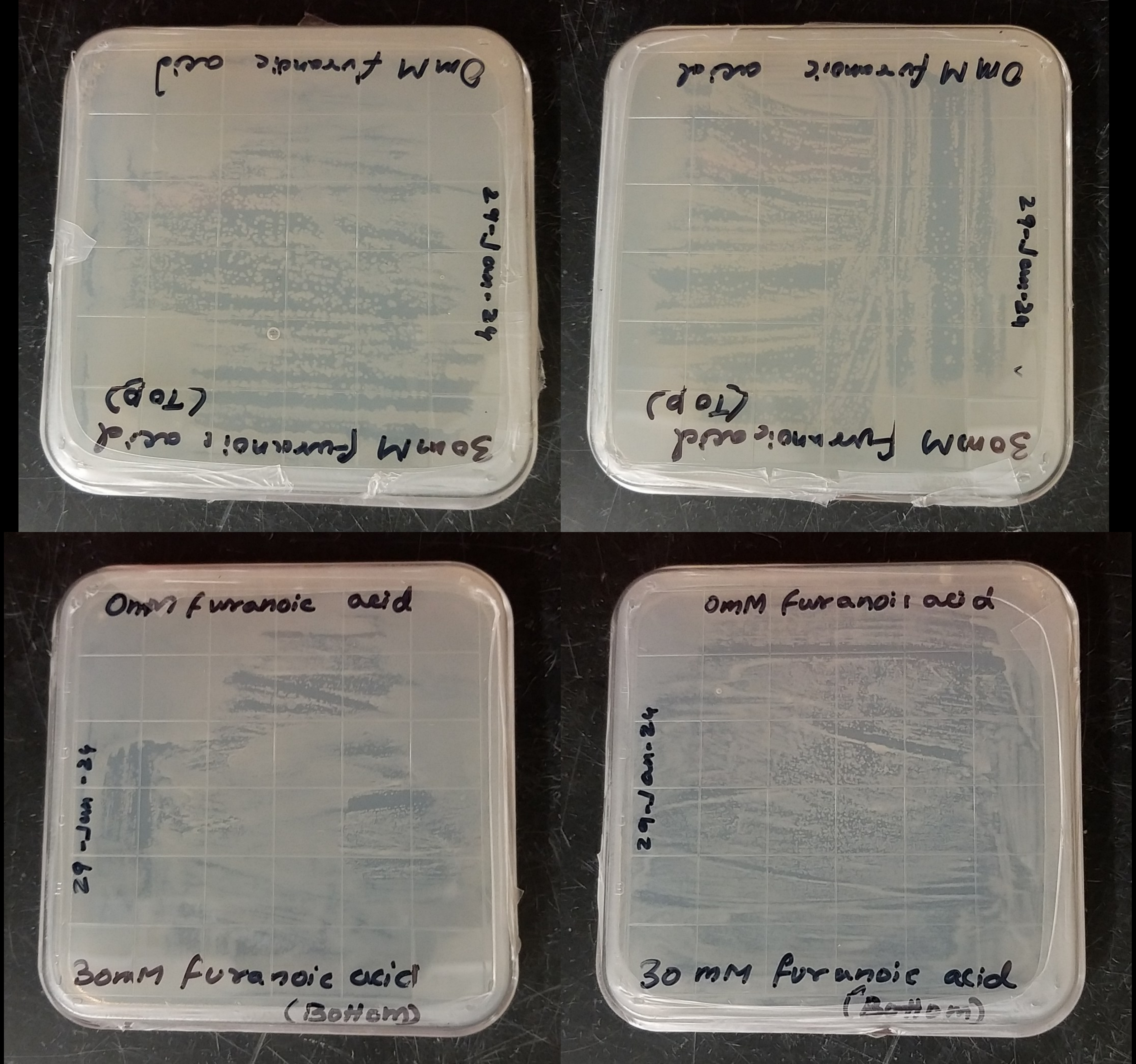
0mM furanoic acid
30mM furanoic acid
(Top)
0mM furanoic acid
30mM furanoic acid
(Top)
0mM furanoic acid
29-Jan-24
30mM furanoic acid
(Bottom)
0mM furanoic acid
29-Jan-24
30 mM furanoic acid
(Bottom)

B

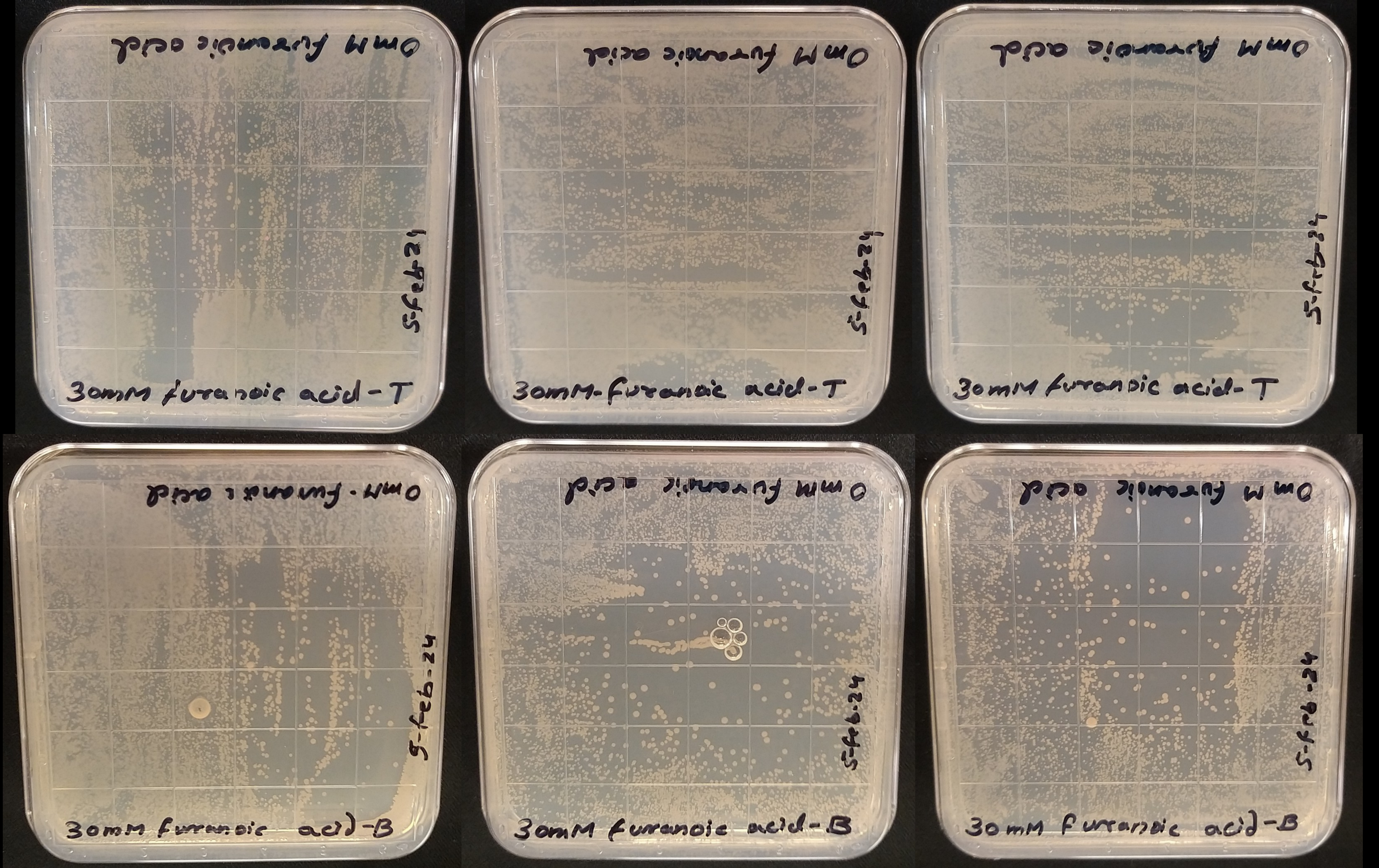
30mM furanoic acid-T
30mM-furanoic acid-T
30mM furanoic acid-T
5-feb-24
30mM furanoic acid-B
30mM furanoic acid-B
30mM furanoic acid-B

C

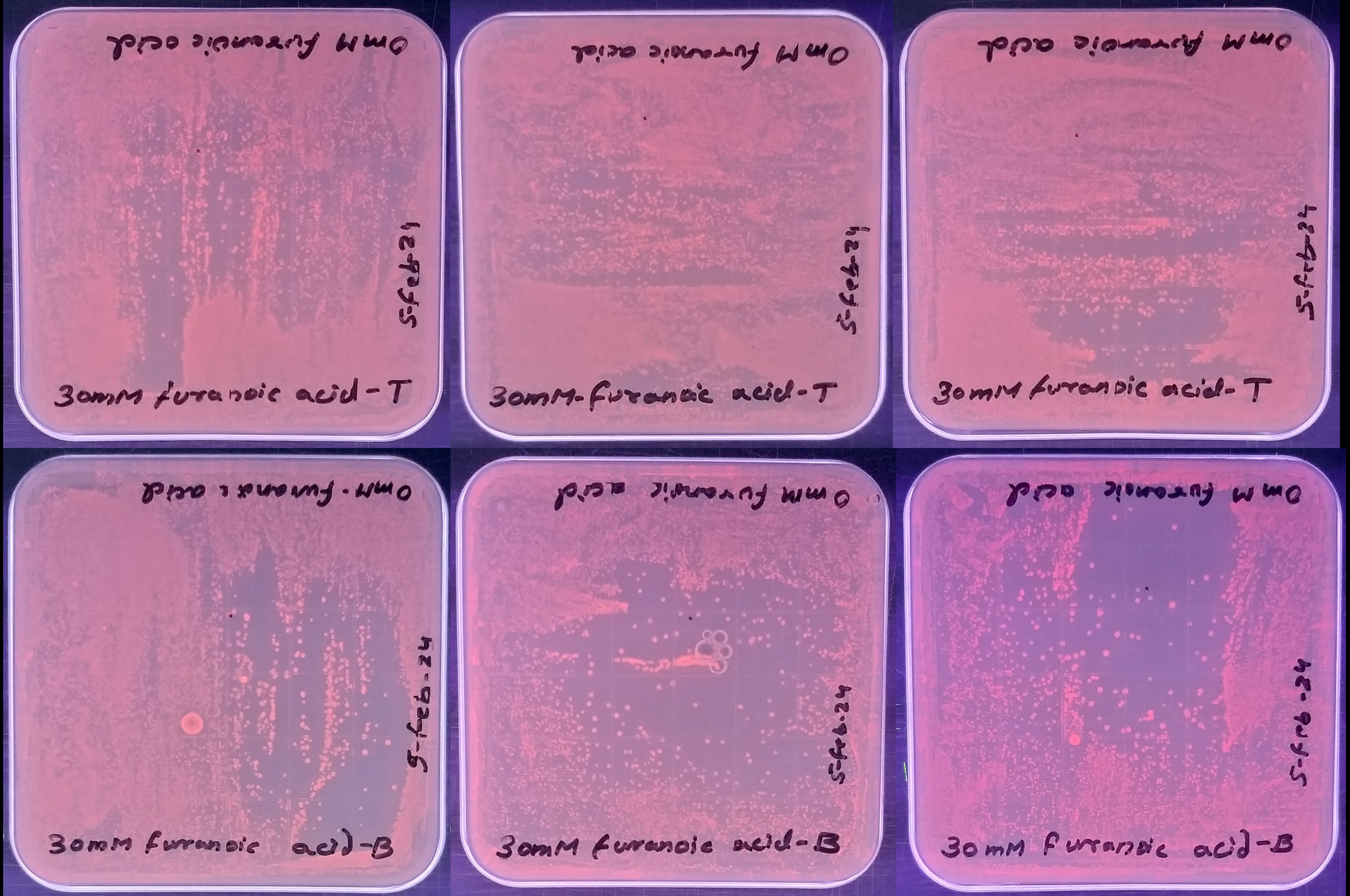
0mM furanoic acid
5-feb-24
30mM furanoic acid-T
30mM furanoic acid-B

### Phloretic acid

D

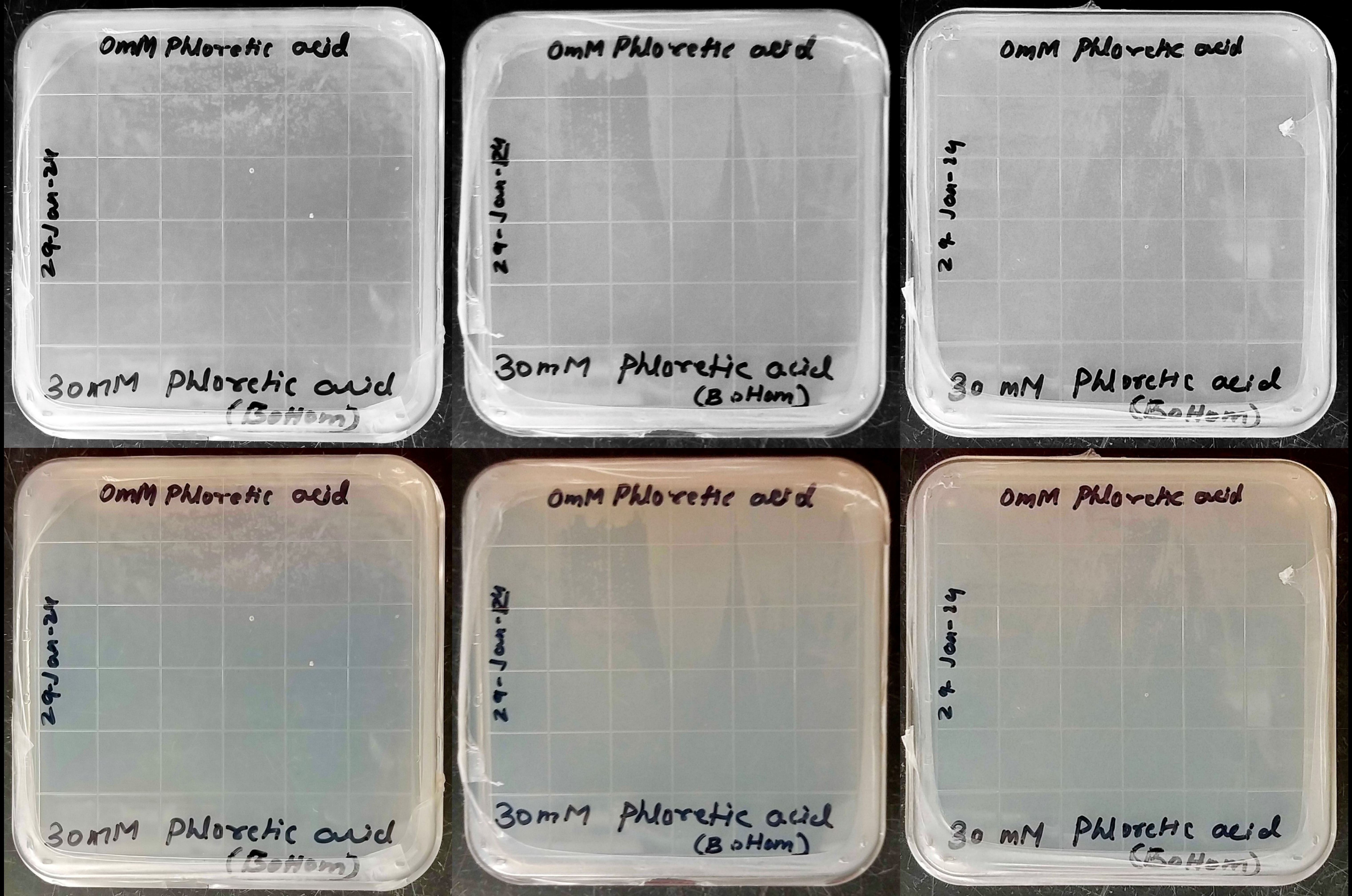
0mM Phloretic acid
29-Jan-24
30mM Phloretic acid
(Bottom)
0mM Phloretic acid
29-Jan-24
30mM Phloretic acid
(Bottom)
0mM Phloretic acid
29 Jan-24
30 mM Phloretic acid
(Bottom)
0mM Phloretic acid
29-Jan-24
30mM Phloretic acid
(Bottom)
0mM Phloretic acid
29-Jan-24
30mM Phloretic acid
(Bottom)
0mM Phloretic acid
29 Jan-24
30 mM Phloretic acid
(Bottom)

E

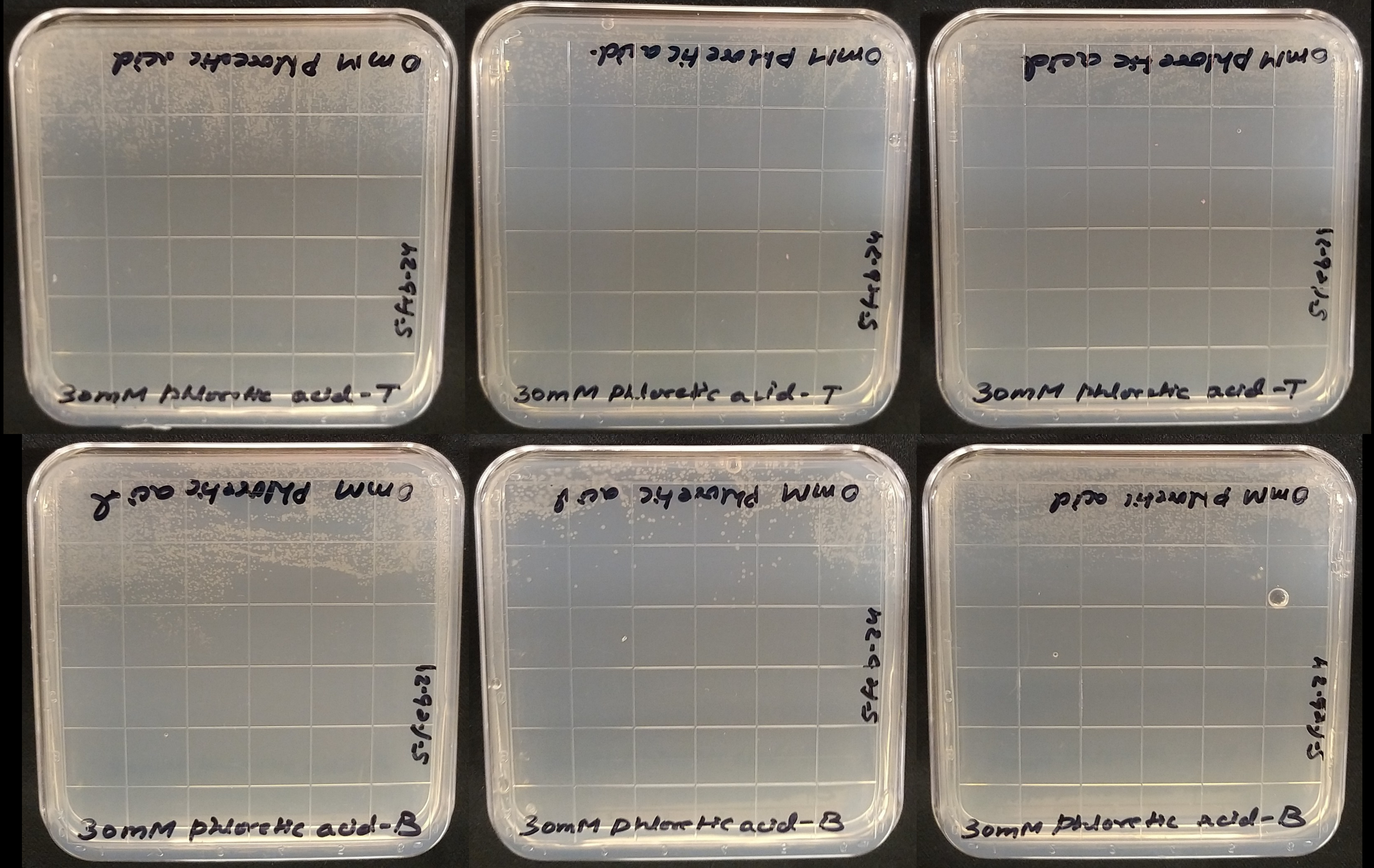
30mM Phloretic acid-T
30mM Phloretic acid-T
30mM Phloretic acid-T
30mM Phloretic acid-B
30mM Phloretic acid-B
30mM Phloretic acid-B

F

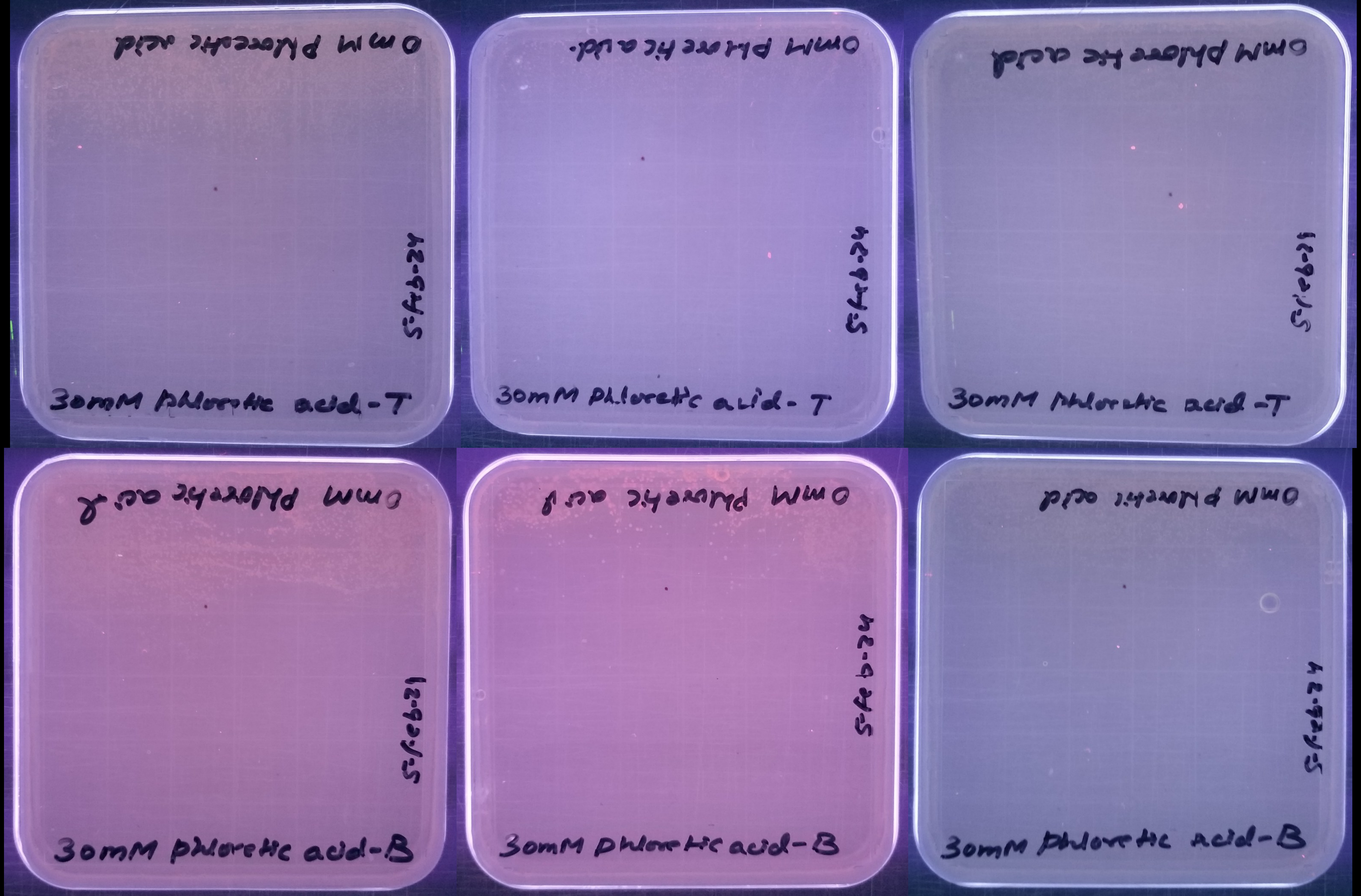
30mM Phloretic acid-T
30mM Phloretic acid-T
30mM Phloretic acid-T
30mM Phloretic acid-B
30mM Phloretic acid-B
30mM Phloretic acid-B

### Mandelic acid

G

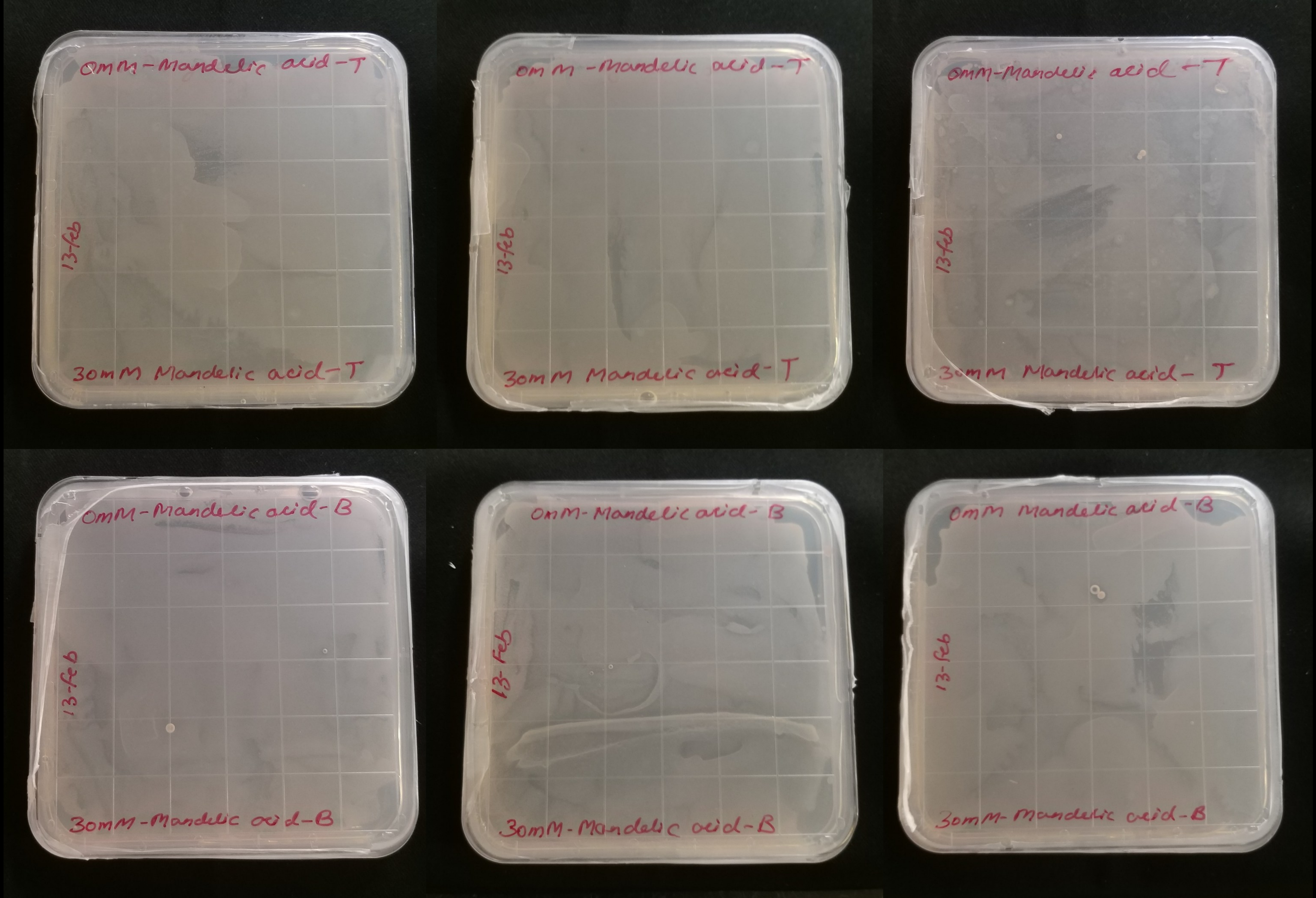

0mM-Mandelic acid-T
13-feb
30mM Mandelic acid-T
0mM-Mandelic acid-T
13-feb
30mM Mandelic acid-T
0mM-Mandelic acid-T
13-feb
30mM Mandelic acid-T
0mM-Mandelic acid-B
13-feb
30mM-Mandelic acid-B
0mM-Mandelic acid-B
13-feb
30mM-Mandelic acid-B
0mM Mandelic acid-B
13-feb
30mM-Mandelic acid-B

H

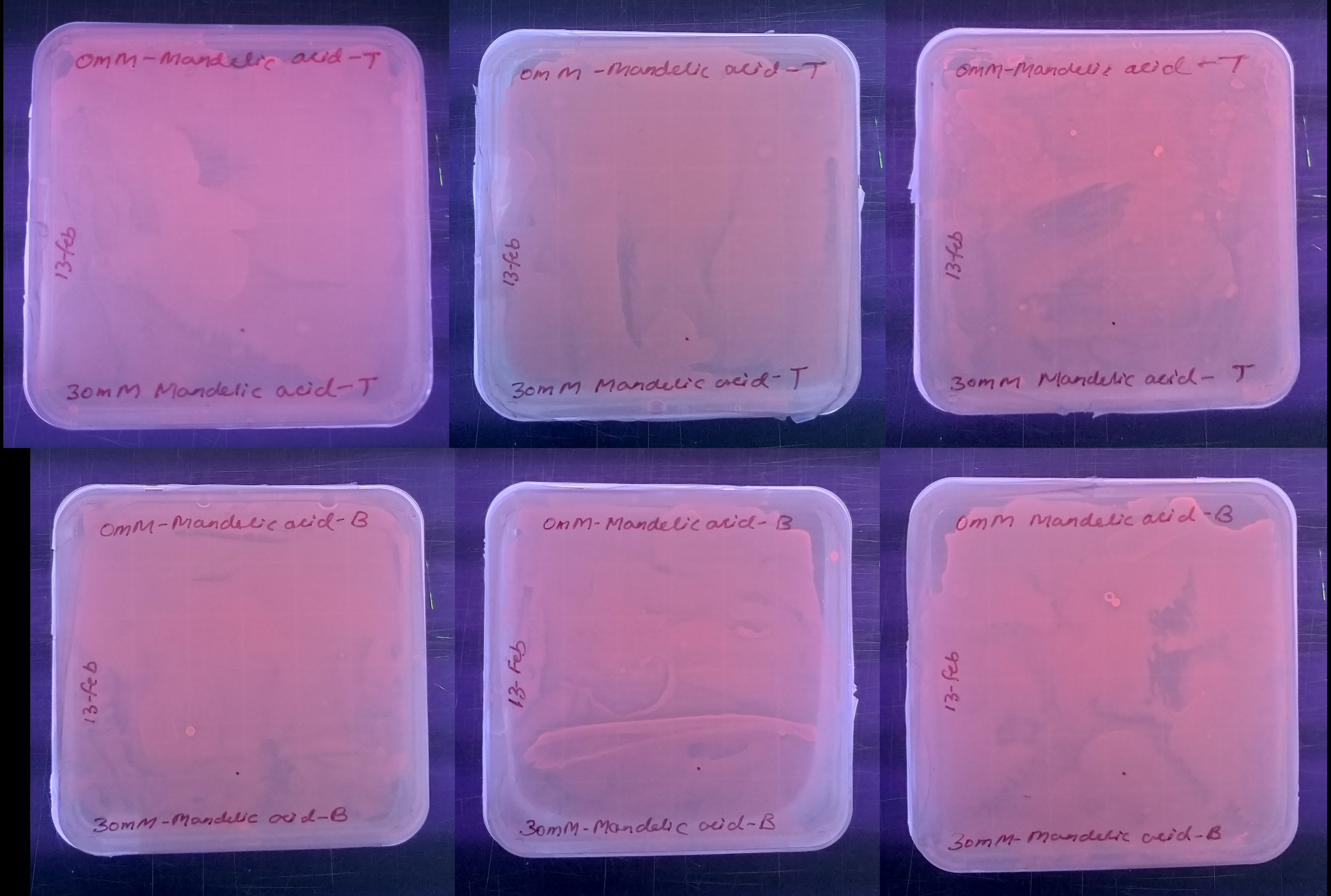
0mM-Mandelic acid-T
13-feb
30mM Mandelic acid-T
0mM-Mandelic acid-T
13-feb
30mM Mandelic acid-T
0mM-Mandelic acid-T
13-feb
30mM Mandelic acid-T
0mM-Mandelic acid-B
13-feb
30mM-Mandelic acid-B
0mM-Mandelic acid-B
13-feb
30mM-Mandelic acid-B
0mM Mandelic acid-B
13-feb
30mM-Mandelic acid-B

### Phenyllactic acid

I

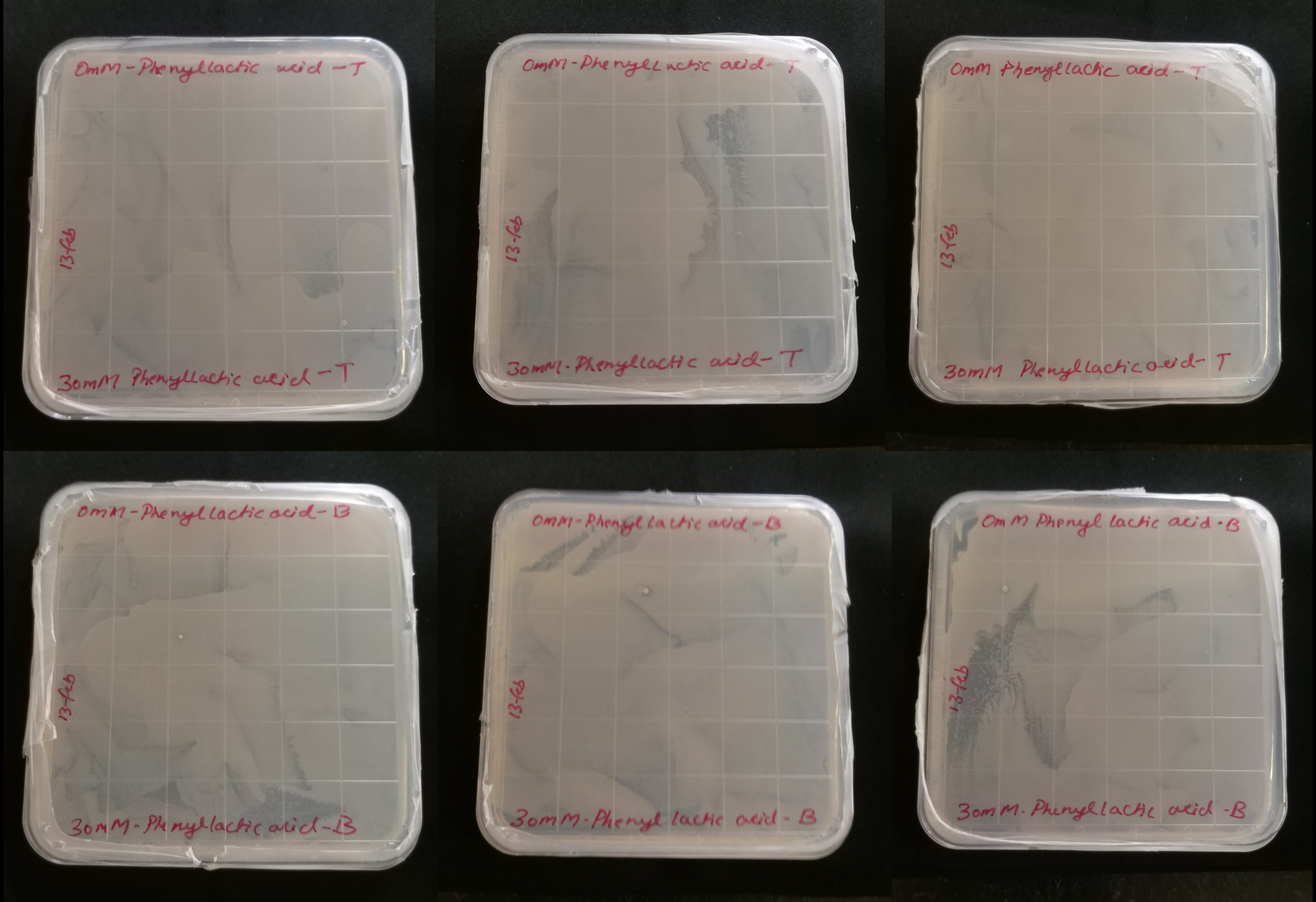
0mM-Phenyllactic acid -T
13-feb
30mM Phenyllactic acid -T
0mM-Phenyllactic acid-T
13-feb
30mM-Phenyllactic acid-T
0mM Phenyllactic acid -T
13-feb
30mM Phenyllactic acid- T
0mM-Phenyllactic acid-B
13-feb
30mM-Phenyllactic acid-B
0mM-Phenyl lactic acid -B
13-feb
30mM-Phenyl lactic acid - B
0m M Phenyl lactic acid-B
13-feb
30mM-Phenyllactic acid -B

J

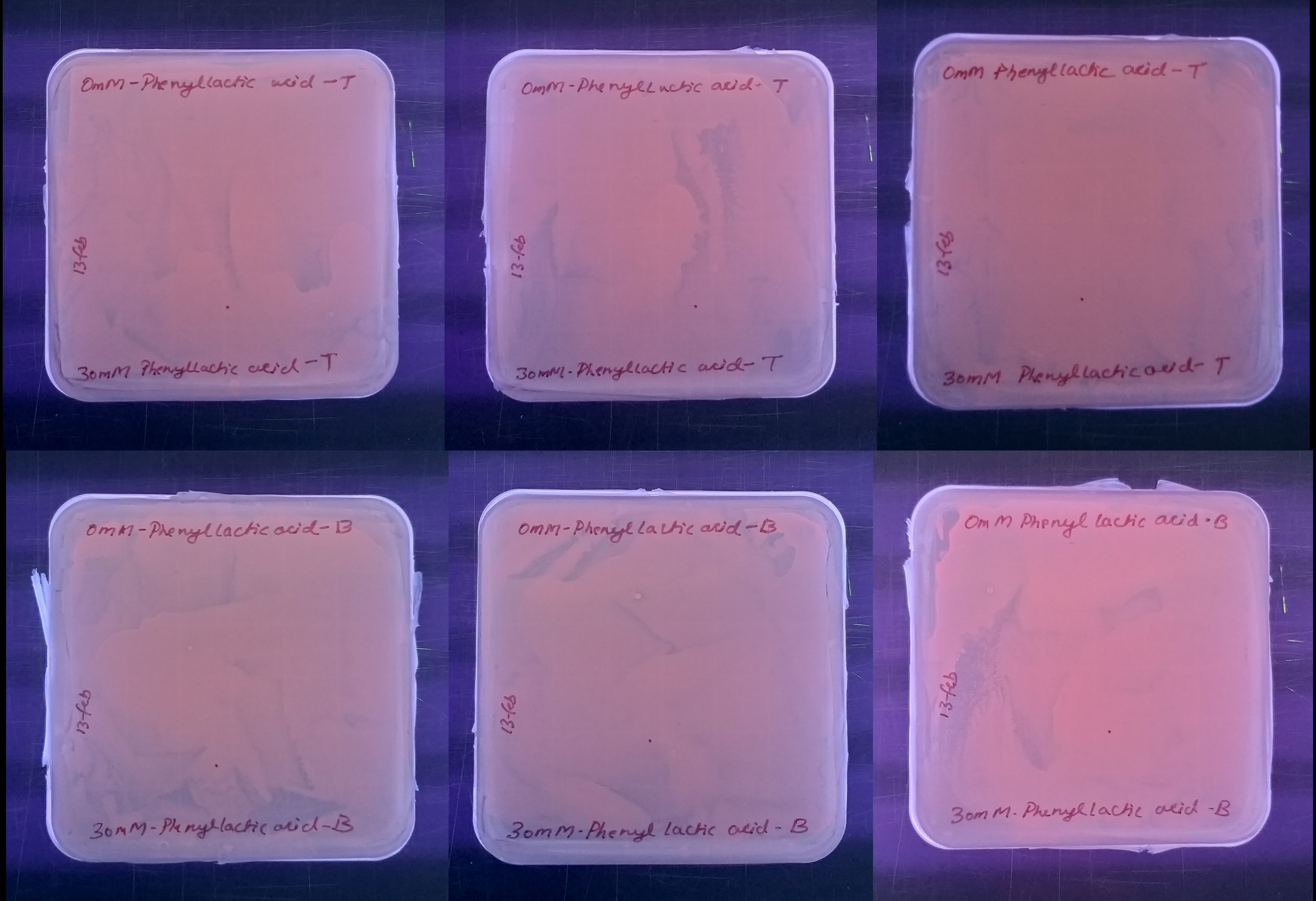

OmM-Phenyllactic acid -T
13-feb
30mM Phenyllactic acid -T
OmM-Phenyllactic acid- T
13-feb
30mM-Phenyllactic acid- T
OmM Phenyllactic acid - T
13-feb
30mM Phenyllactic acid- T
OmM-Phenyllactic acid- B
13-feb
30mM-Phenyllactic acid-B
OmM-Phenyl lactic acid-B
13-feb
30mM-Phenyl lactic acid - B
OmM Phenyl lactic acid - B
13-feb
30mM-Phenyllactic acid -B

### 2-hydroxy-4-phenylbutyric acid

K

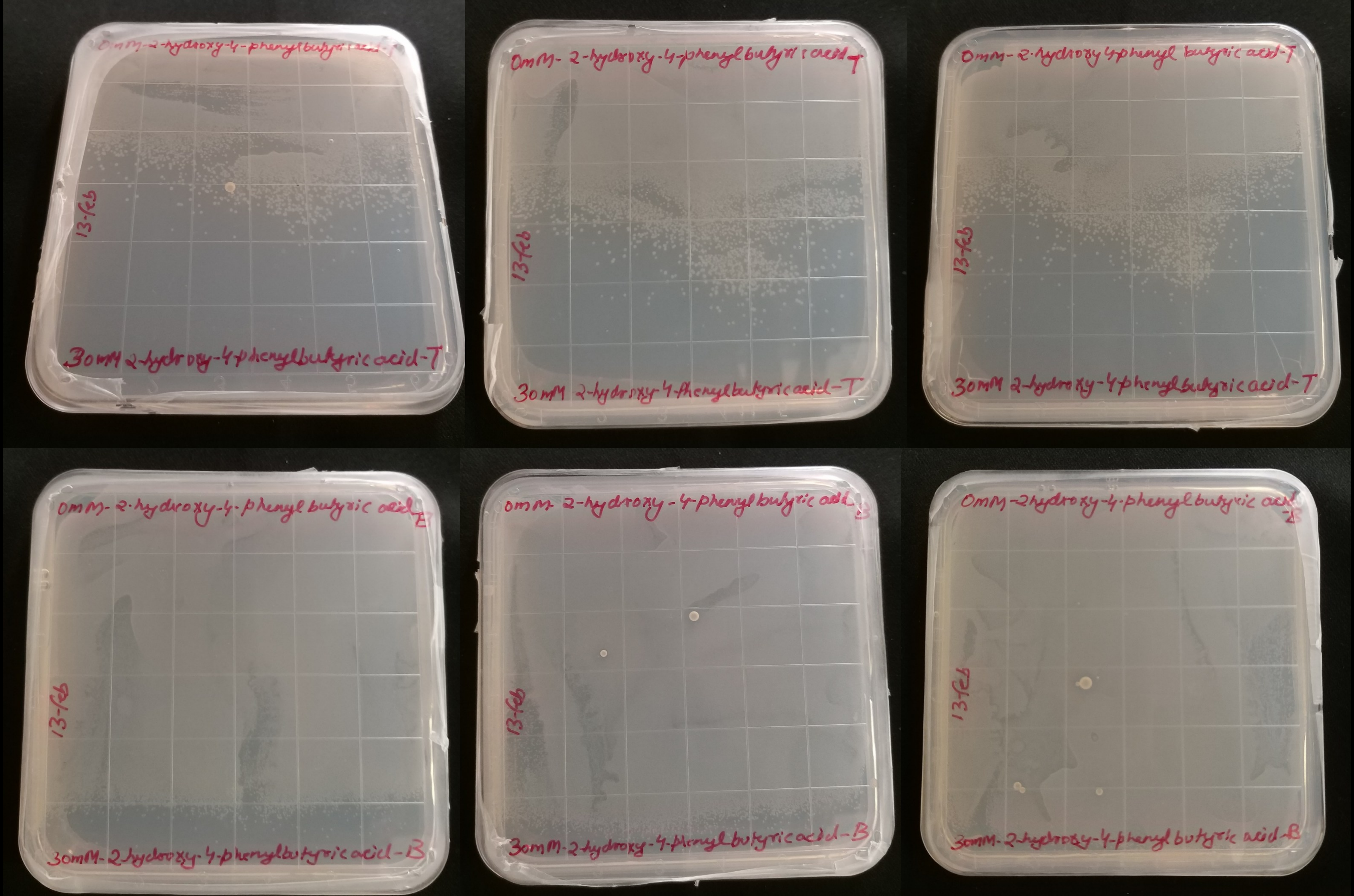
0mM-2-hydroxy-4-phenylbutyric acid-T
13-feb
30mM 2-hydroxy-4-phenylbutyric acid-T
0mM-2-hydroxy-4-phenylbutyric acid-T
13-feb
30mM 2-hydroxy-4-phenylbutyric acid-T
0mM-2-hydroxy-4-phenyl butyric acid-T
13-feb
30mM 2-hydroxy-4-phenylbutyric acid-T
0mM-2-hydroxy-4-phenyl butyric acid-B
13-feb
30mM-2-hydroxy-4-phenylbutyric acid-B
0mM-2-hydroxy-4-phenylbutyric acid-B
13-feb
30mM-2-hydroxy-4-phenyl butyric acid-B
0mM-2-hydroxy-4-phenyl butyric acid-B
13-feb
30mM-2-hydroxy-4-phenyl butyric acid-B

L

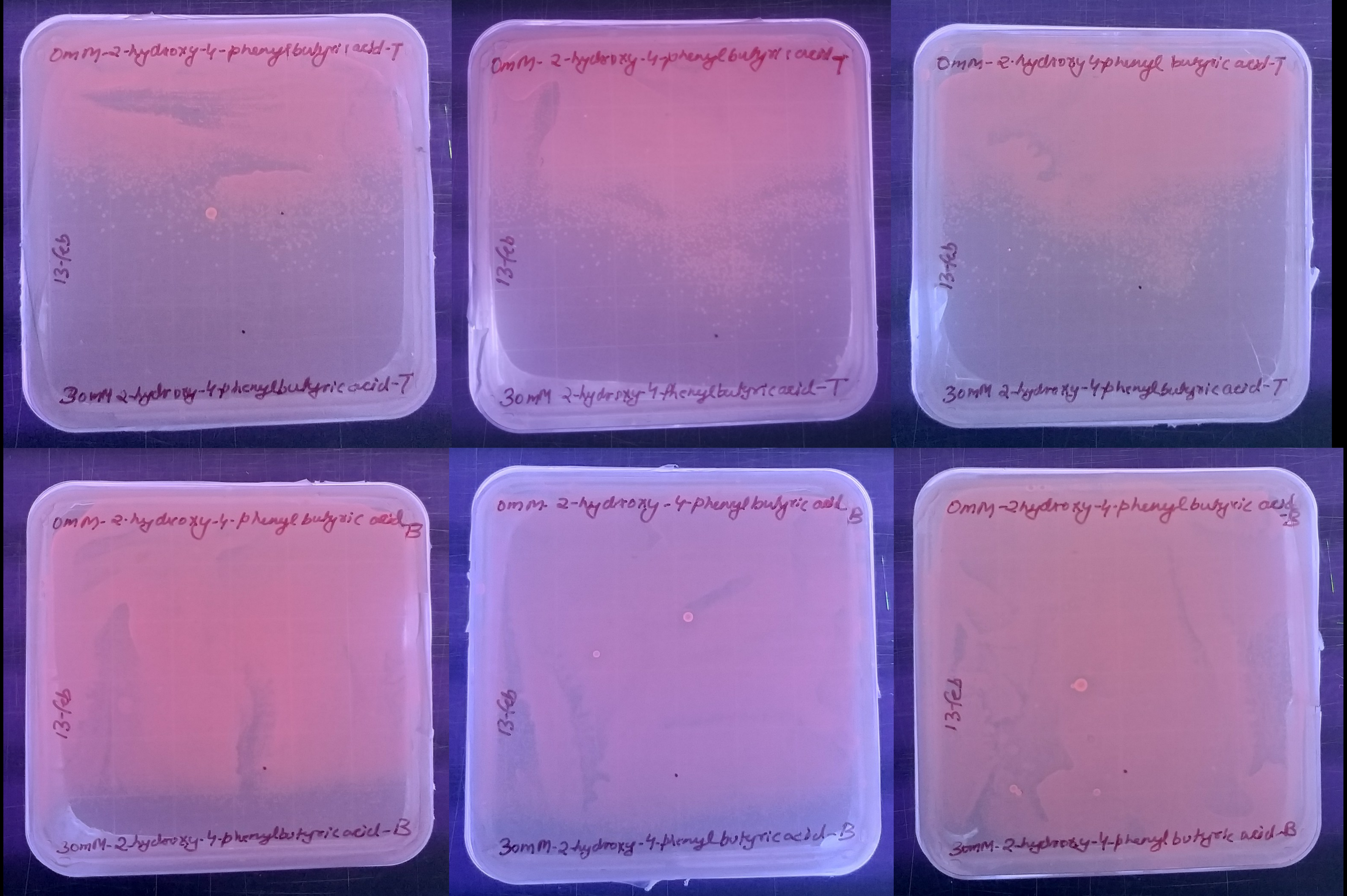
0mM-2-hydroxy-4-phenylbutyric acid-T
13-feb
30mM 2-hydroxy-4-phenylbutyric acid-T
0mM-2-hydroxy-4-phenylbutyric acid-T
13-feb
30mM 2-hydroxy-4-phenylbutyric acid-T
0mM-2-hydroxy-4-phenyl butyric acid-T
13-feb
30mM 2-hydroxy-4-phenylbutyric acid-T
0mM-2-hydroxy-4-phenyl butyric acid-B
13-feb
30mM-2-hydroxy-4-phenylbutyric acid-B
0mM-2-hydroxy-4-phenyl butyric acid-B
13-feb
30mM-2-hydroxy-4-phenyl butyric acid-B
0mM-2hydroxy-4-phenyl butyric acid-B
13-feb
30mM-2-hydroxy-4-phenyl butyric acid-B

### 6-hydroxycaproic acid

M

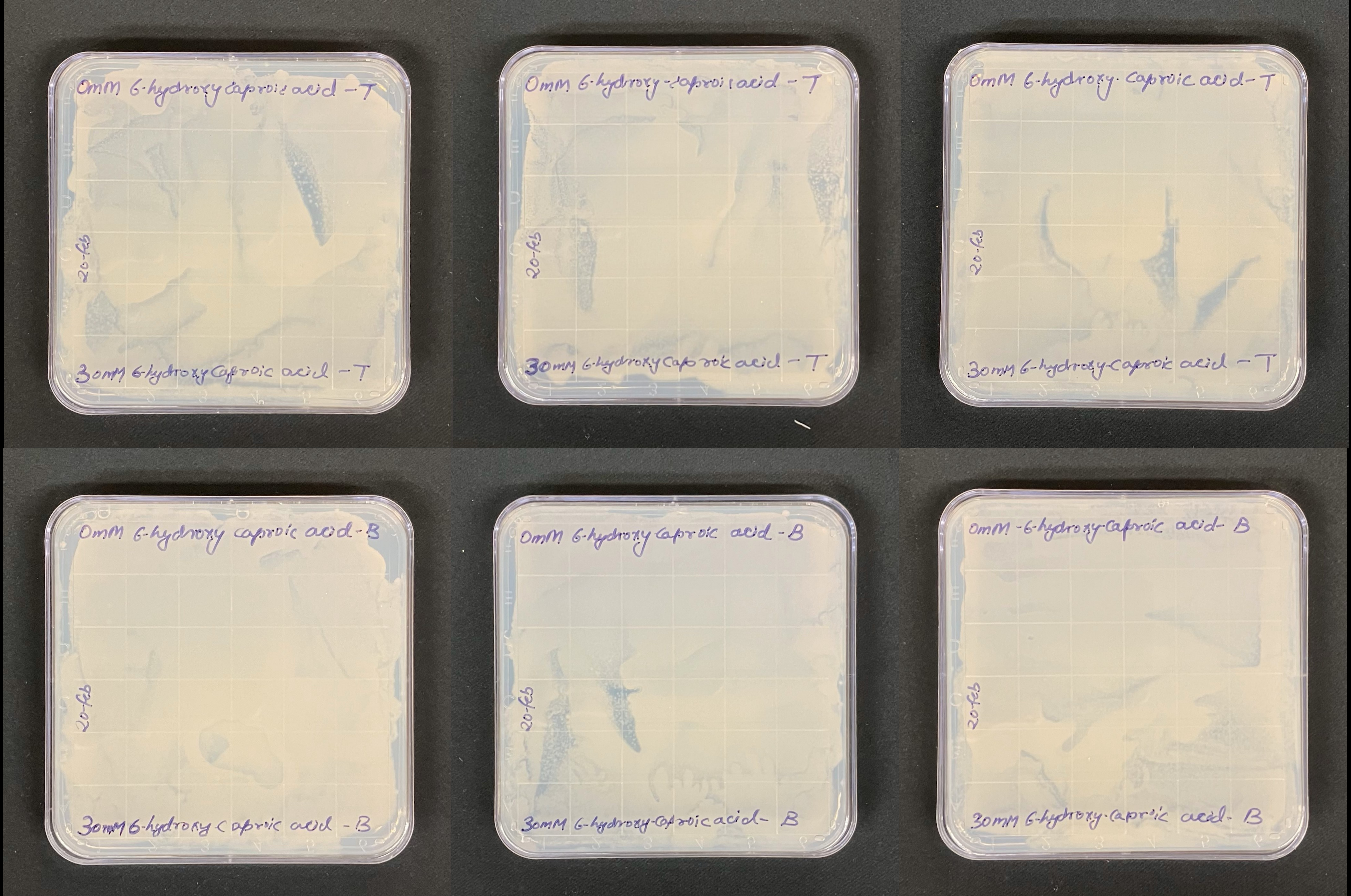

0mM 6-hydroxy caproic acid - T
20-feb
30mM 6-hydroxy caproic acid - T
0mM 6-hydroxy-caproic acid - T
20-feb
30mM 6-hydroxy caproic acid - T
0mM 6-hydroxy caproic acid - T
20-feb
30mM 6-hydroxy-caproic acid - T
0mM 6-hydroxy caproic acid - B
20-feb
30mM 6-hydroxy caproic acid - B
0mM 6-hydroxy caproic acid - B
20-feb
30mM 6-hydroxy-caproic acid - B
0mM -6-hydroxy-caproic acid - B
20-feb
30mM 6-hydroxy-caproic acid - B

N

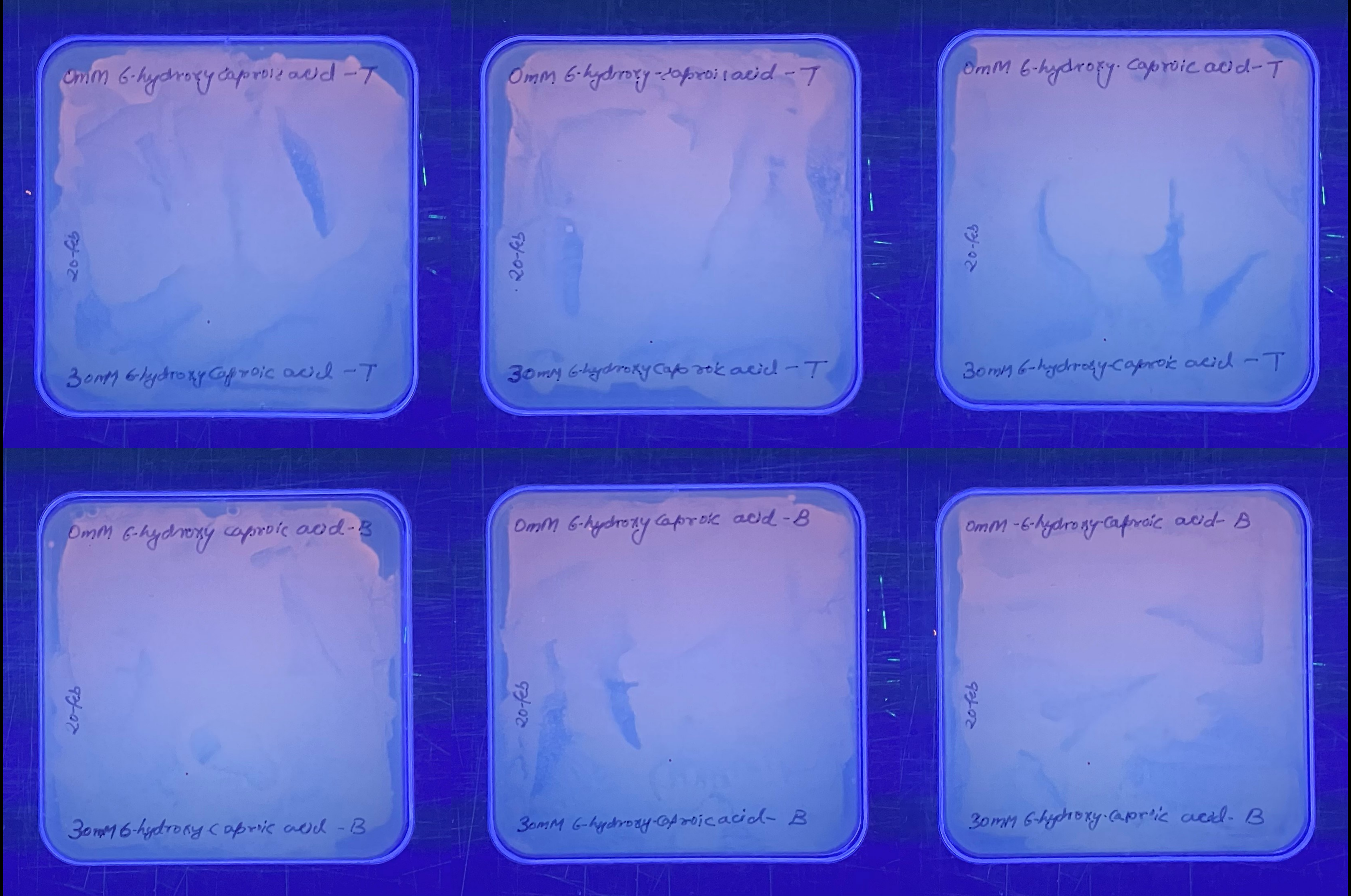
0mM 6-hydroxy caproic acid -T
20-feb
30mM 6-hydroxy caproic acid -T
0mM 6-hydroxy-caproic acid -T
20-feb
30mM 6-hydroxy caproic acid -T
0mM 6-hydroxy. caproic acid -T
20-feb
30mM 6-hydroxy-caproic acid -T
0mM 6-hydroxy caproic acid -B
20-feb
30mM 6-hydroxy capric acid -B
0mM 6-hydroxy caproic acid -B
20-feb
30mM 6-hydroxy-caproic acid- B
0mM -6-hydroxy-caproic acid- B
20-feb
30mM 6-hydroxy-capric acid- B

### 4-hydroxybutyric acid

O

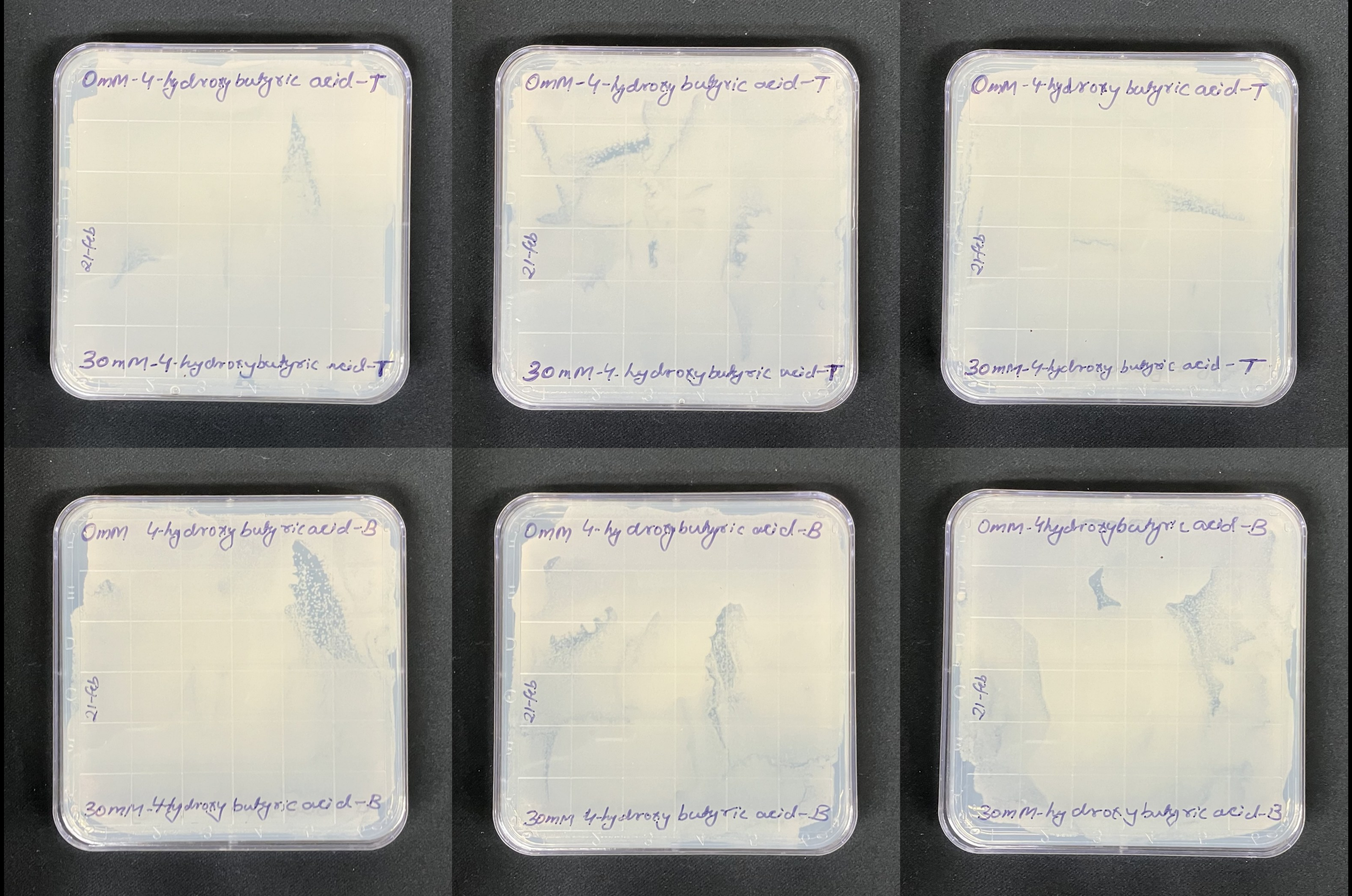
0mM-4-hydroxy butyric acid-T
21-feb
30mM-4-hydroxybutyric acid-T
0mM-4-hydroxy butyric acid-T
21-feb
30mM-4-hydroxybutyric acid-T
0mM-4-hydroxy butyric acid-T
21-feb
30mM-4-hydroxy butyric acid-T
0mM 4-hydroxy butyric acid-B
21-feb
30mM-4-hydroxy butyric acid-B
0mM 4-hydroxy butyric acid-B
21-feb
30mM 4-hydroxy butyric acid-B
0mM-4hydroxybutyric acid-B
21-feb
30mM-hydroxy butyric acid-B

P

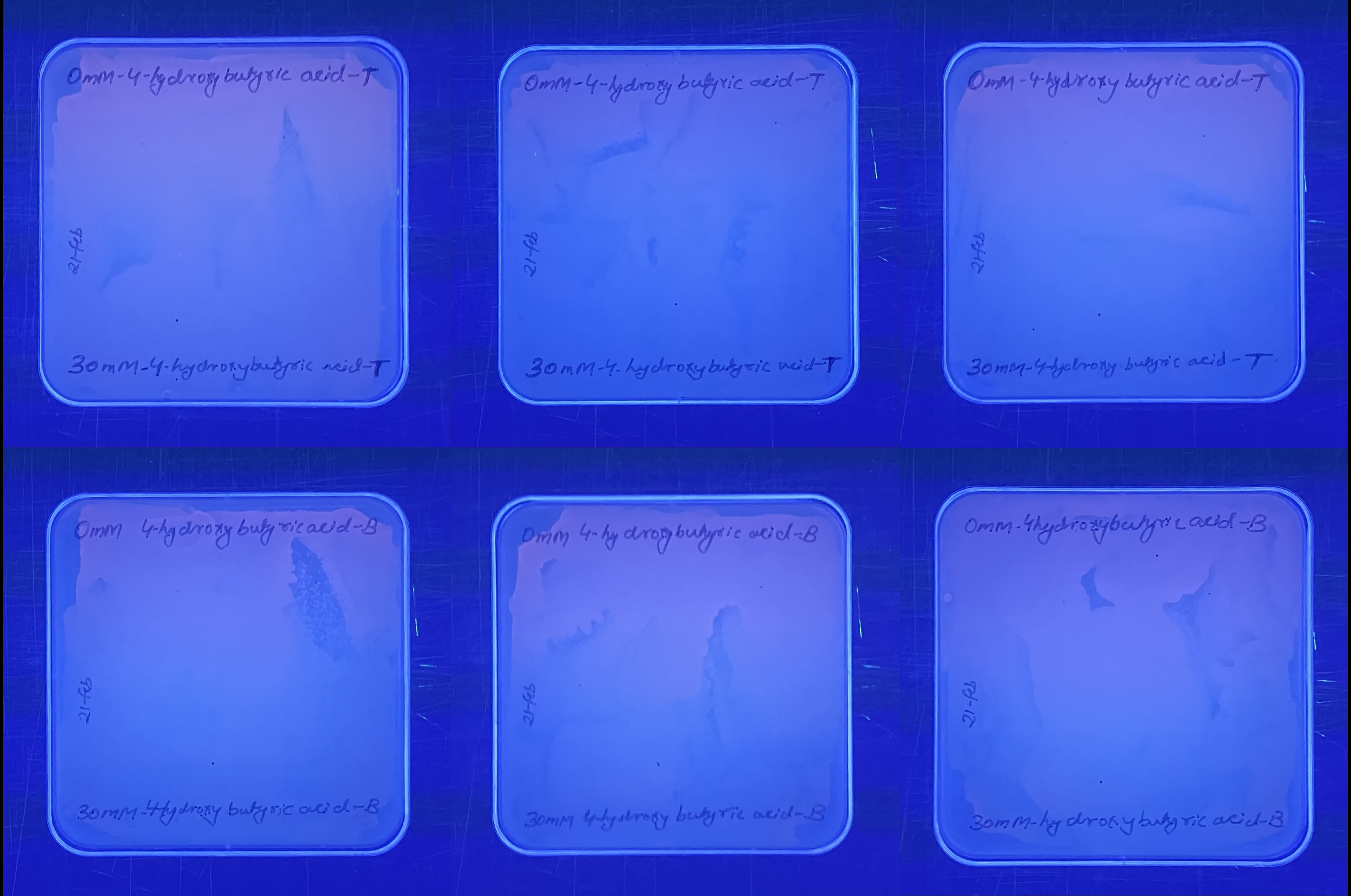
0mM-4-hydroxy butyric acid-T
21-feb
30mM-4-hydroxybutyric acid-T
0mM-4-hydroxy butyric acid-T
21-feb
30mM-4-hydroxy butyric acid-T
0mM-4-hydroxy butyric acid-T
21-feb
30mM-4-hydroxy butyric acid-T
0mM 4-hydroxy butyric acid-B
21-feb
30mM-4hydroxy butyric acid-B
0mM 4-hydroxy butyric acid-B
21-feb
30mM 4hydroxy butyric acid-B
0mM-4hydroxybutyric acid-B
21-feb
30mM-hydroxy butyric acid-B

### 3-hydroxypropionic acid

Q

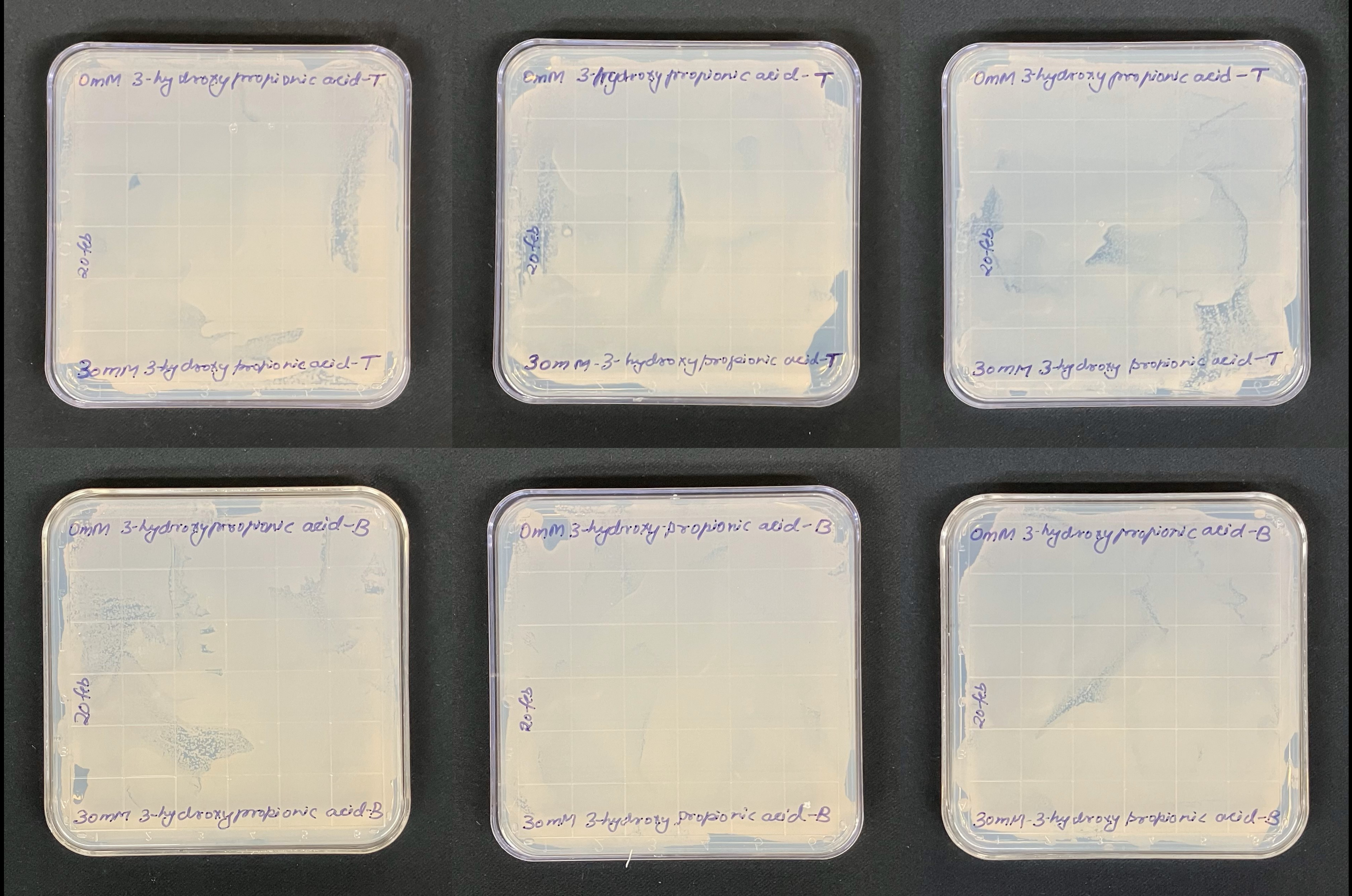
0mM 3-hydroxy propionic acid-T
20 feb
30mM 3-hydroxy propionic acid-T
0mM 3-hydroxy propionic acid-T
20 feb
30mM-3-hydroxy propionic acid-T
0mM 3-hydroxy propionic acid-T
20 feb
30mM 3-hydroxy propionic acid-T
0mM 3-hydroxy propionic acid-B
20 feb
30mM 3-hydroxy propionic acid-B
0mM 3-hydroxy-propionic acid-B
20 feb
30mM 3-hydroxy propionic acid-B
0mM 3-hydroxy propionic acid-B
20 feb
30mM-3-hydroxy propionic acid-B

R

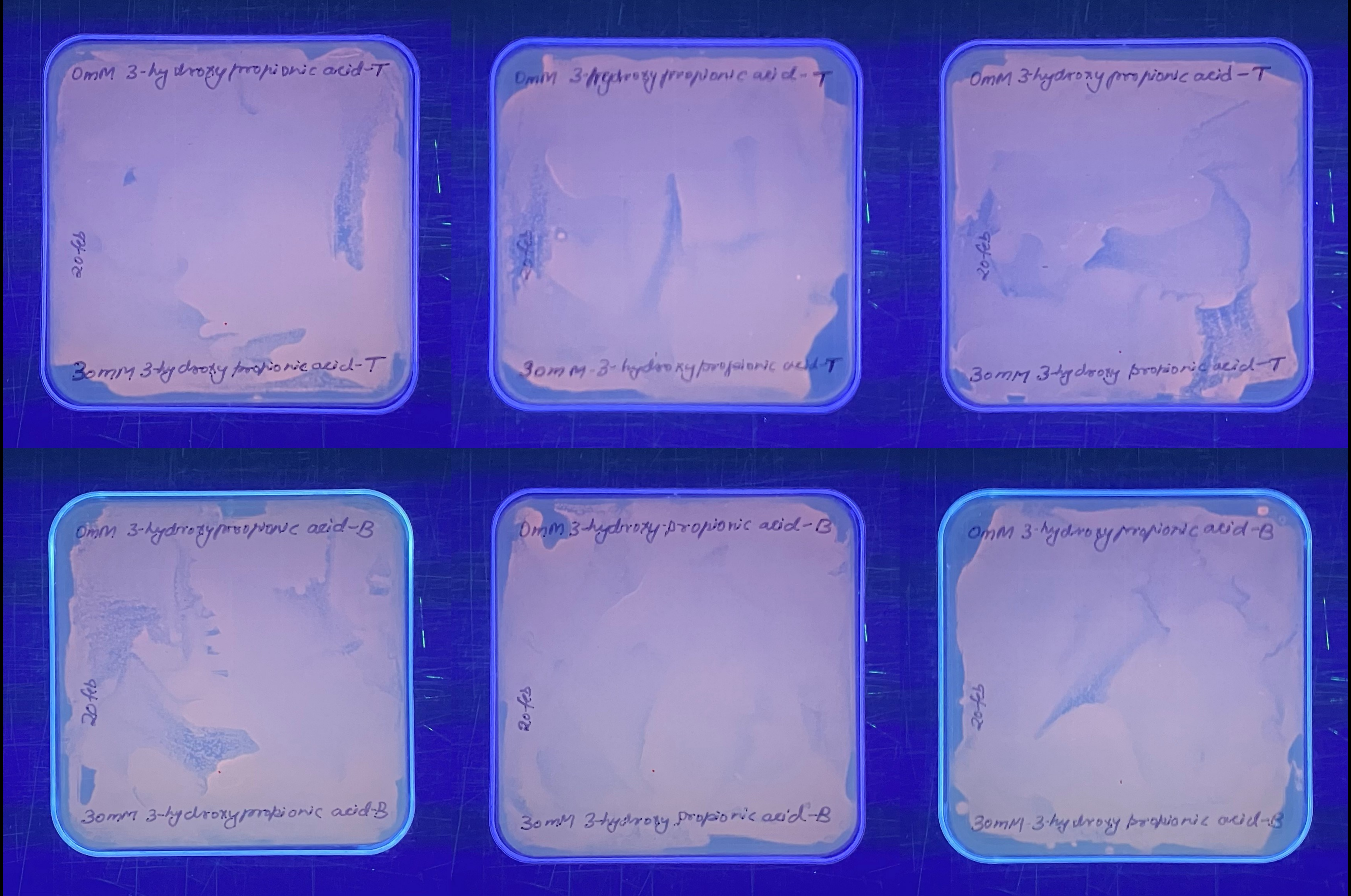
0mM 3-hydroxy propionic acid-T
20 feb
30mM 3-hydroxy propionic acid-T
0mM 3-hydroxy propionic acid-T
20 feb
30mM-3-hydroxy propionic acid-T
0mM 3-hydroxy propionic acid-T
20 feb
30mM 3-hydroxy propionic acid-T
0mM 3-hydroxy propionic acid-B
20 feb
30mM 3-hydroxy propionic acid-B
0mM 3-hydroxy propionic acid-B
20 feb
30mM 3-hydroxy propionic acid-B
0mM 3-hydroxy propionic acid-B
20 feb
30mM 3-hydroxy propionic acid-B

### Lactic acid

S

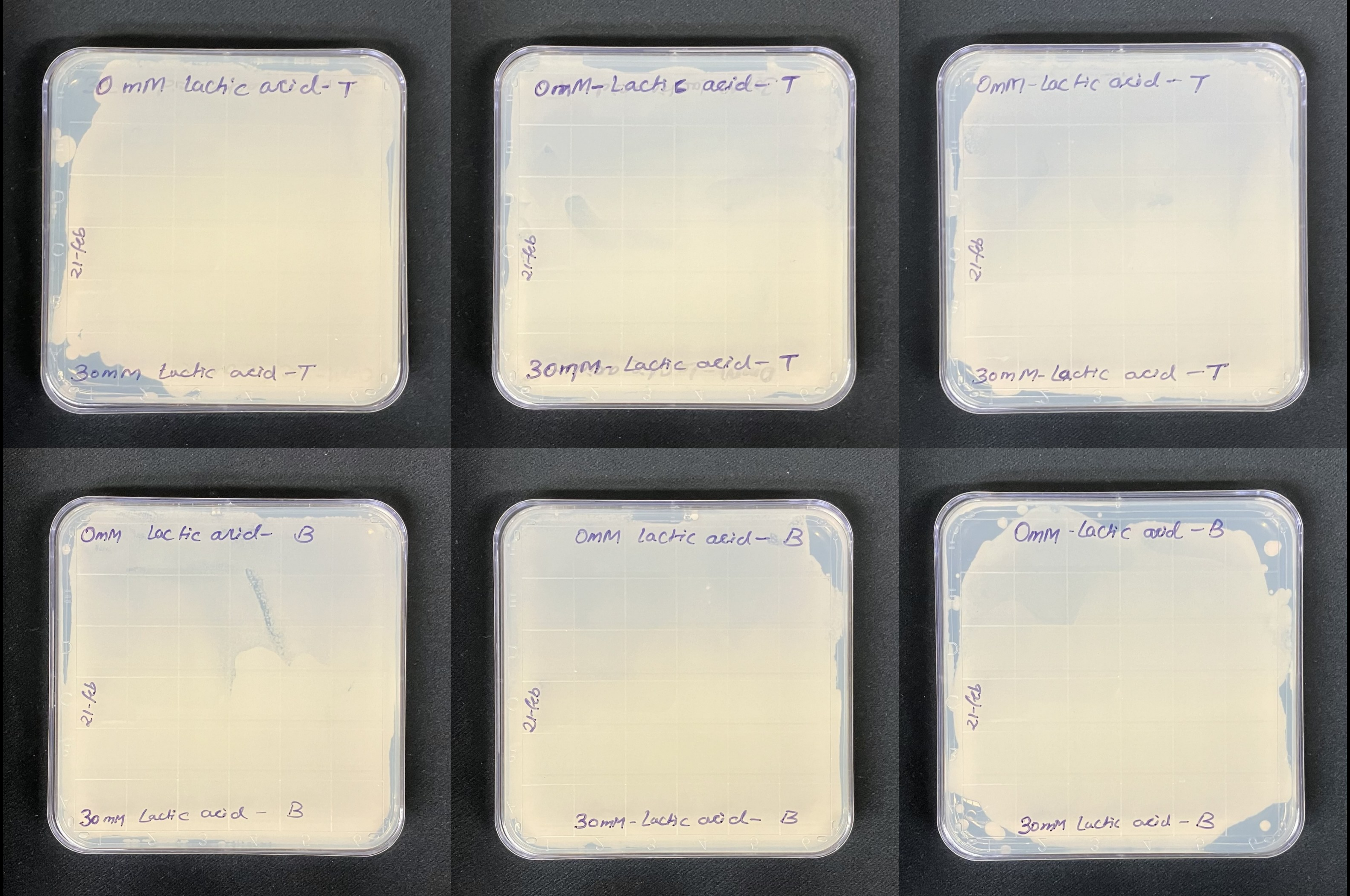
0 mM lactic acid- T
21-feb
30mM Lactic acid -T
0mM-Lactic acid- T
21-feb
30mM-Lactic acid- T
0mM-Lactic acid - T
21-feb
30mM-Lactic acid -T
0mM Lactic acid- B
21-feb
30mM Lactic acid - B
0mM lactic acid- B
21-feb
30mM-Lactic acid- B
0mM -Lactic acid - B
21-feb
30mM Lactic acid - B

T

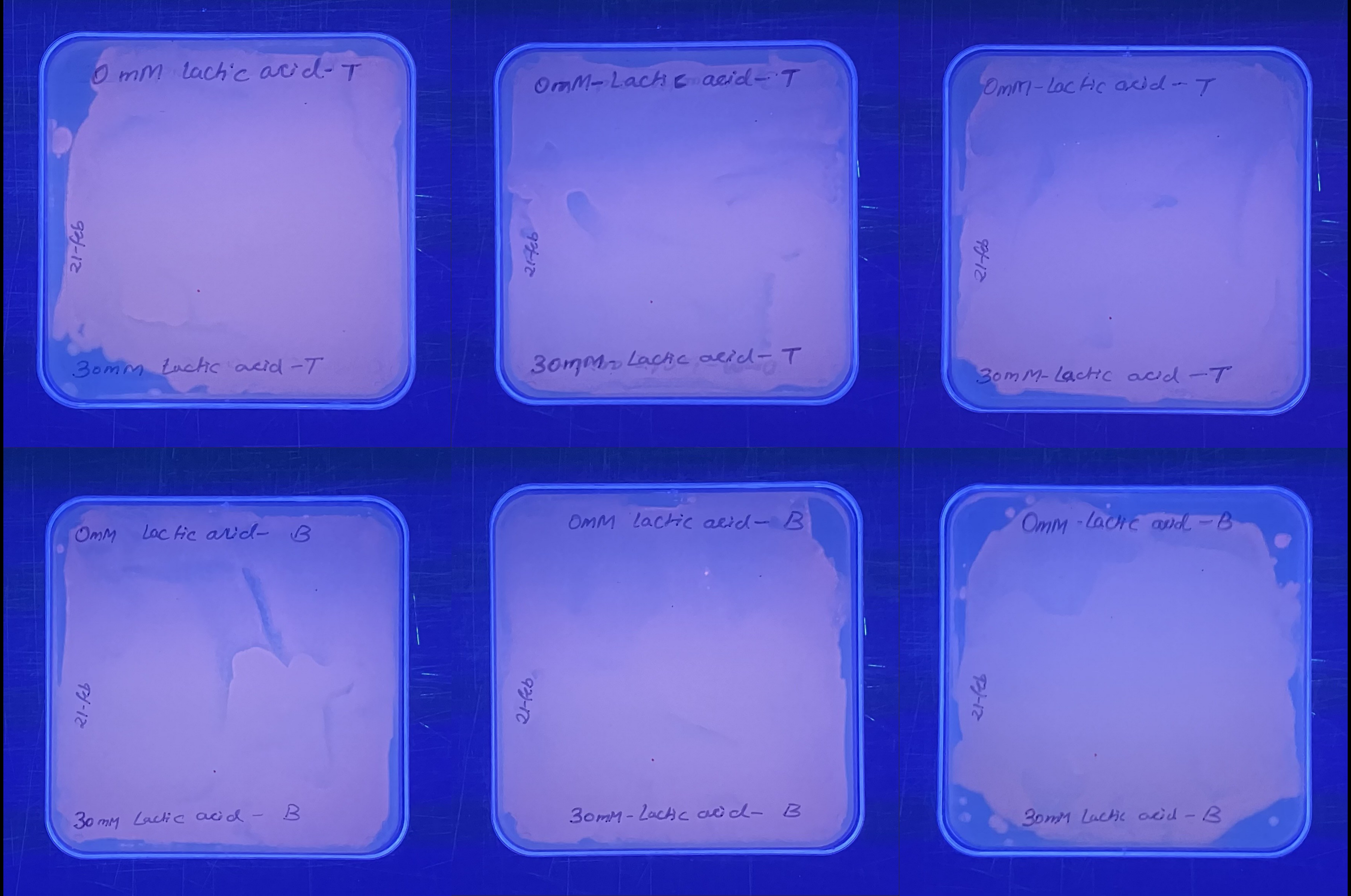
0 mM lactic acid - T
21-feb
30mM Lactic acid - T
0mM - Lactic acid - T
21-feb
30mM - Lactic acid - T
0mM - Lactic acid - T
21-feb
30mM - Lactic acid - T
0mM lactic acid - B
21-feb
30mM Lactic acid - B
0mM lactic acid - B
21-feb
30mM - Lactic acid - B
0mM - Lactic acid - B
21-feb
30mM Lactic acid - B

### Glycolic acid

U

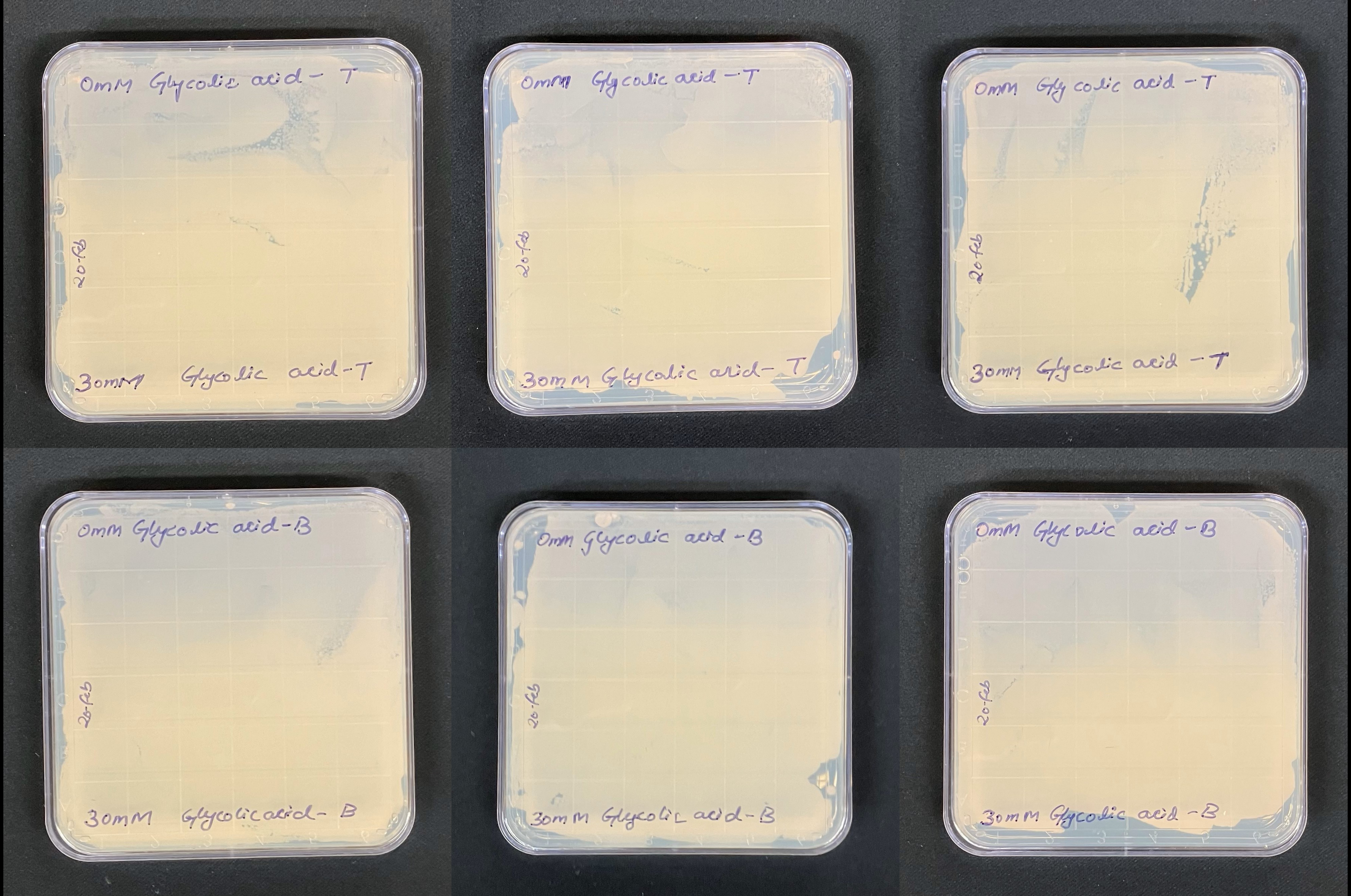
0mM Glycolic acid - T
20-feb
30mM Glycolic acid - T
0mM Glycolic acid - T
20-feb
30mM Glycolic acid - T
0mM Glycolic acid - T
20-feb
30mM Glycolic acid - T
0mM Glycolic acid - B
20-feb
30mM Glycolicacid - B
0mM Glycolic acid - B
20-feb
30mM Glycolic acid - B
0mM Glycolic acid - B
20-feb
30mM Glycolic acid - B

V

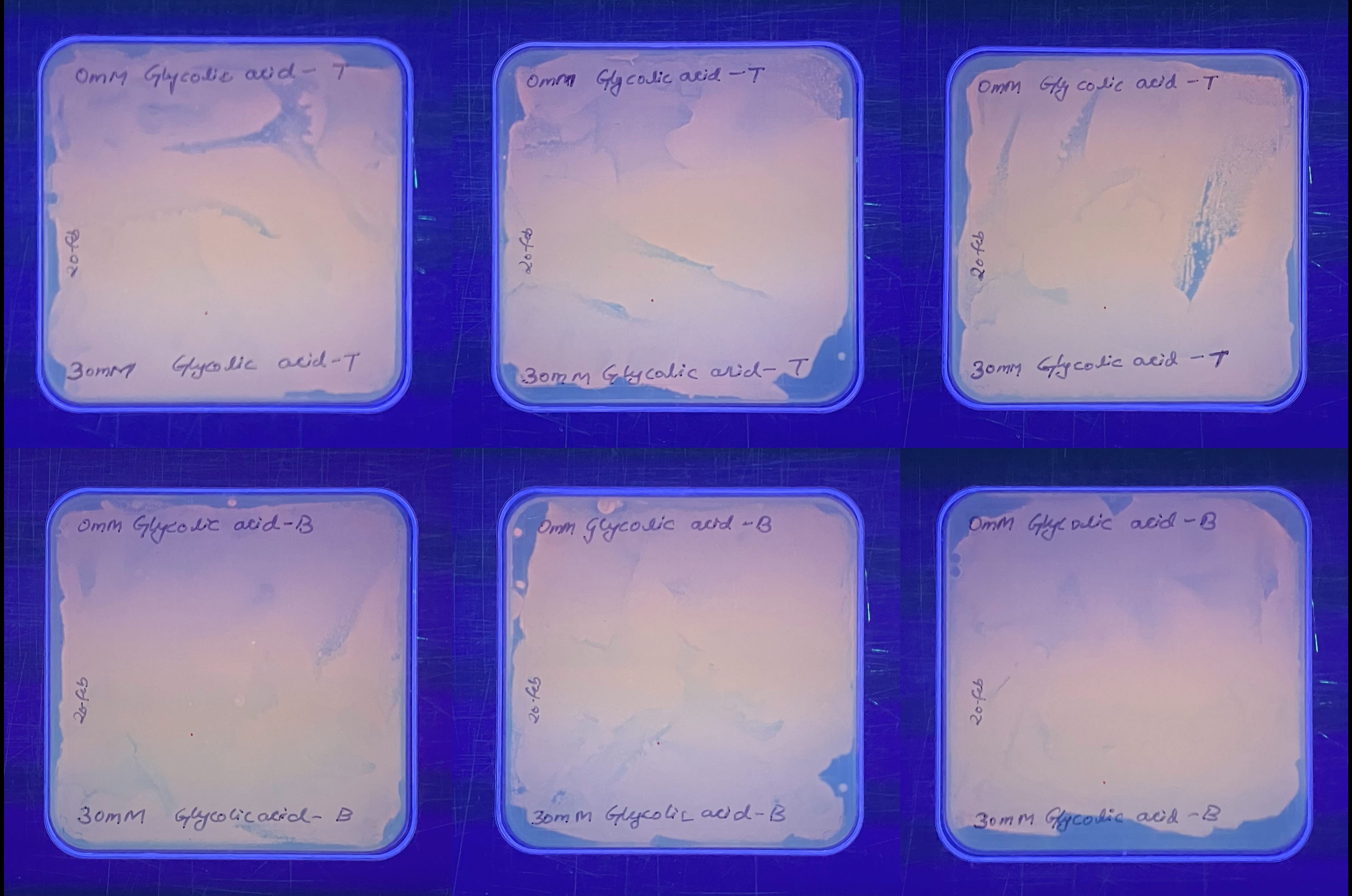
0mM Glycolic acid - T
20 feb
30mM Glycolic acid - T
0mM Glycolic acid - T
20 feb
30mM Glycolic acid - T
0mM Glycolic acid - T
20 feb
30mM Glycolic acid - T
0mM Glycolic acid - B
20 feb
30mM Glycolicacid - B
0mM Glycolic acid - B
20 feb
30mM Glycolic acid - B
0mM Glycolic acid - B
20 feb
30mM Glycolic acid - B
