## Extended Data 2 for "Novel Aromatic Polyhydroxyalkanoates from Engineered *Cupriavidus necator* H16: Expanding Bio-Polyester Synthesis from Renewable Feedstocks"

NMR spectra of bio-polyesters formed by wild-type and engineered *C. necator* H16

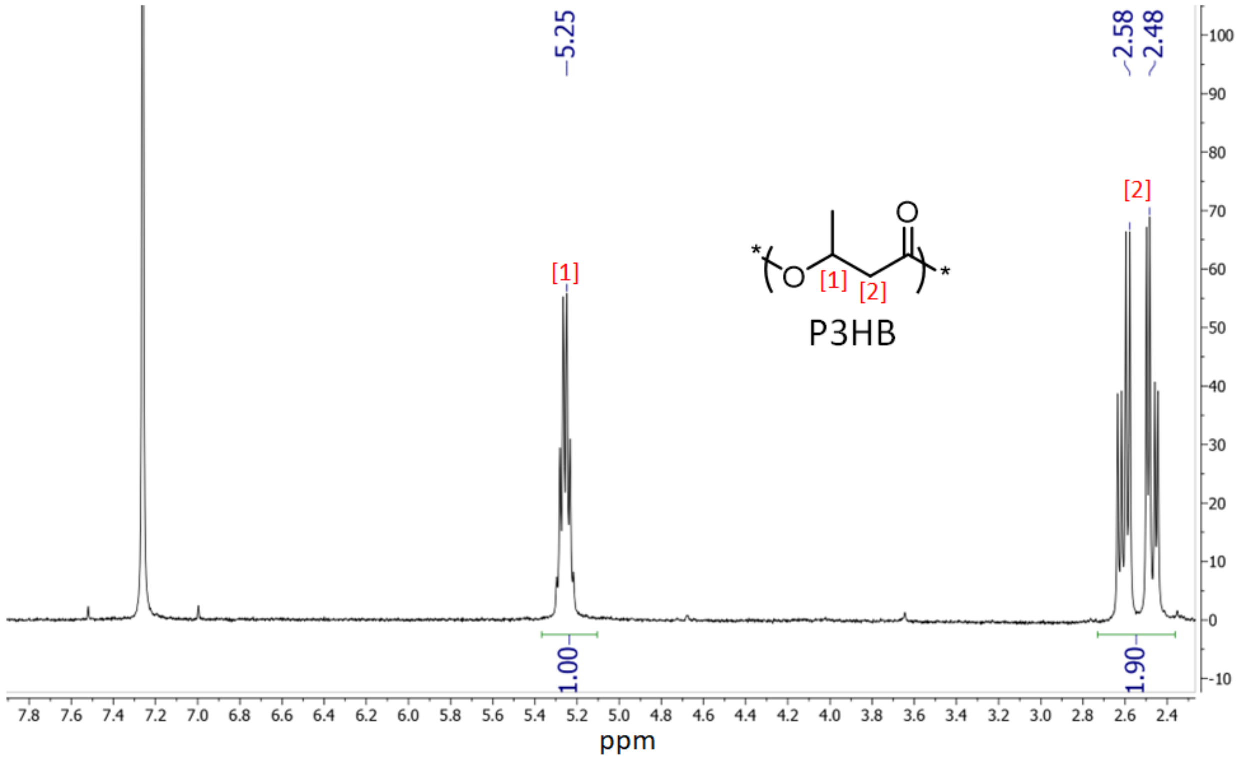


**Figure E2A: $^1$H-NMR in $CDCl_3$ of natural PHB produced by *C. necator* H16 (wild-type).**

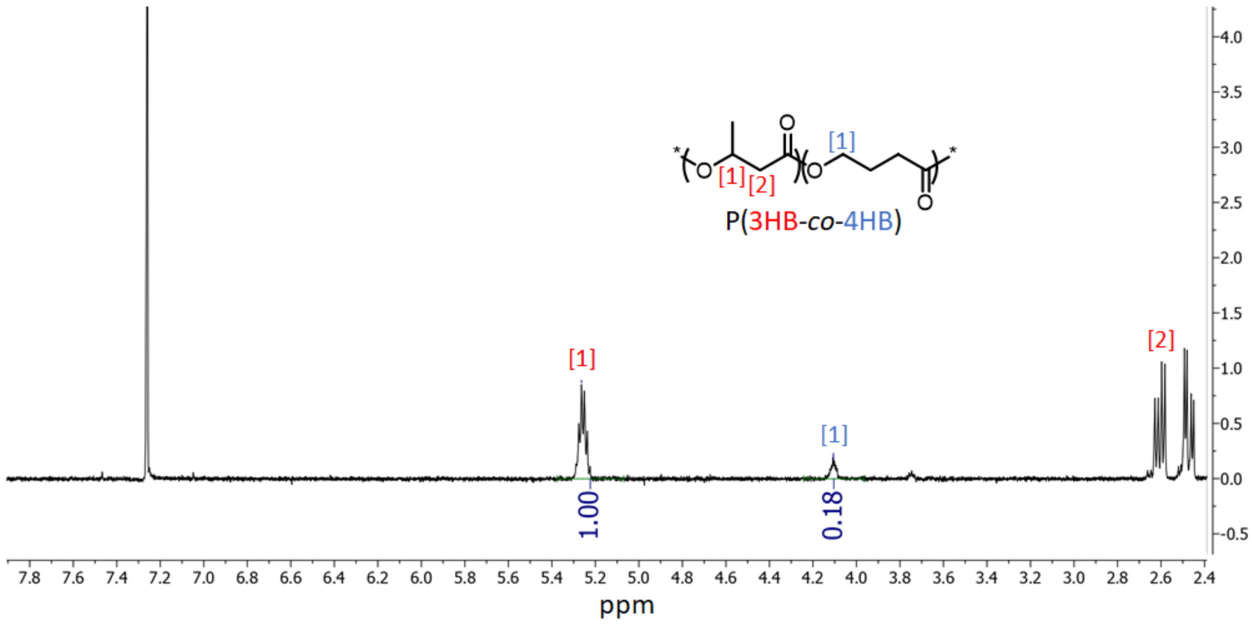


**Figure E2B: $^1$H-NMR in $CDCl_3$ of PHA produced by *C. necator* H16 (wild-type) co-fed with 4-hydroxybutyrate.**

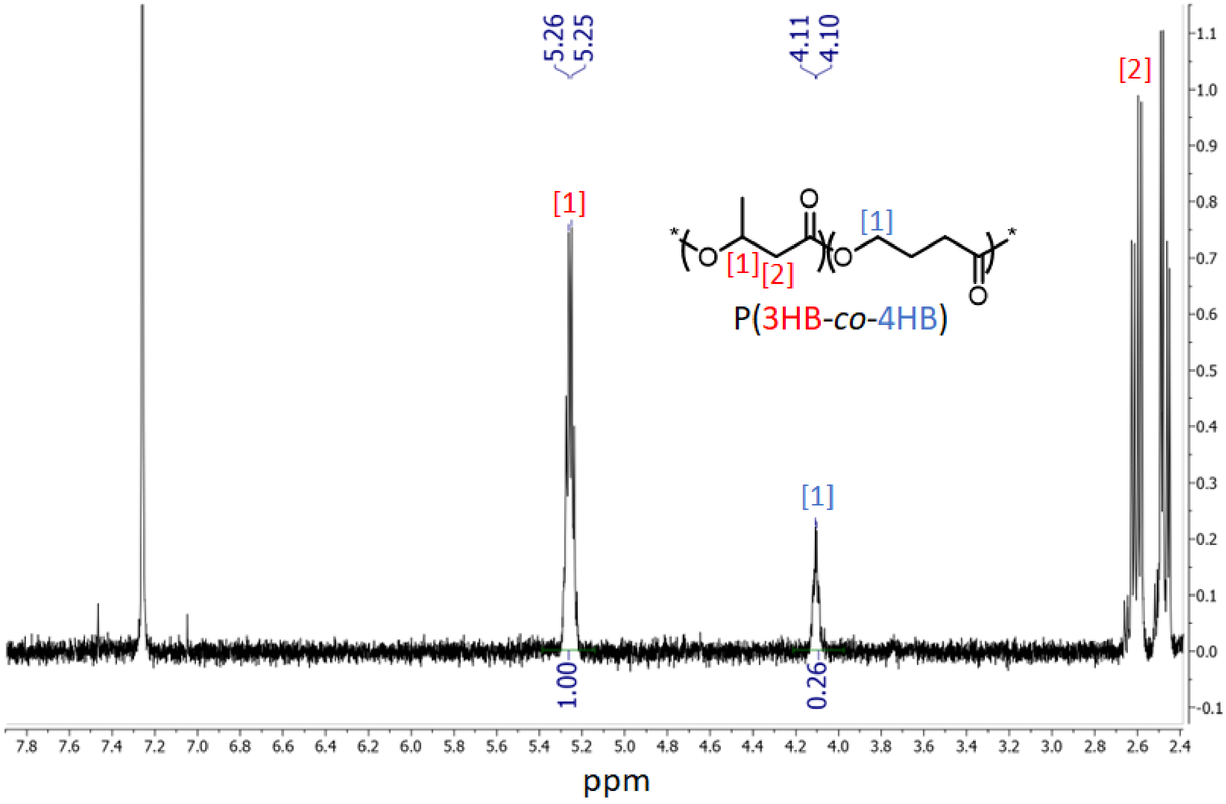


**Figure E2C: $^1$H-NMR in $CDCl_3$ of PHA produced by *C. necator* H16 (wild-type) co-fed with 6-hydroxycaproate.**

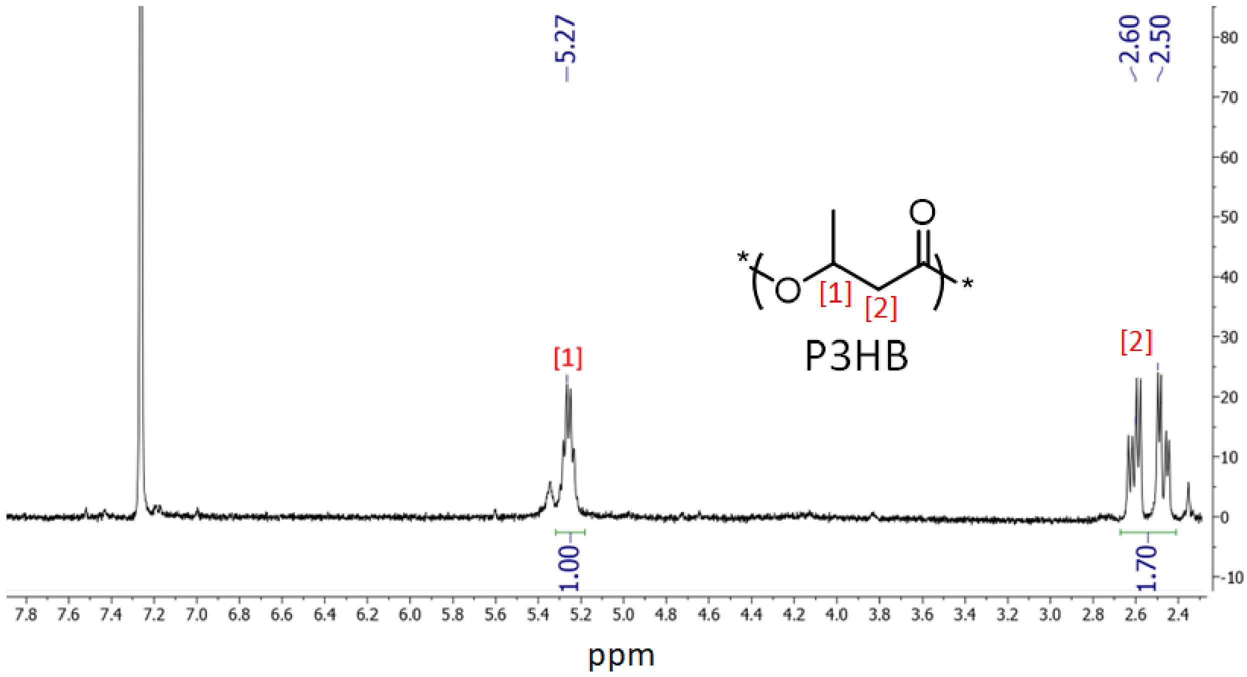


**Figure E2D: [1]H-NMR in $CDCl_3$ of PHA produced by *C. necator* H16 Δ*phaC1* pCM66Tp$_{BAD}$:*pct540-phaC1437*.**

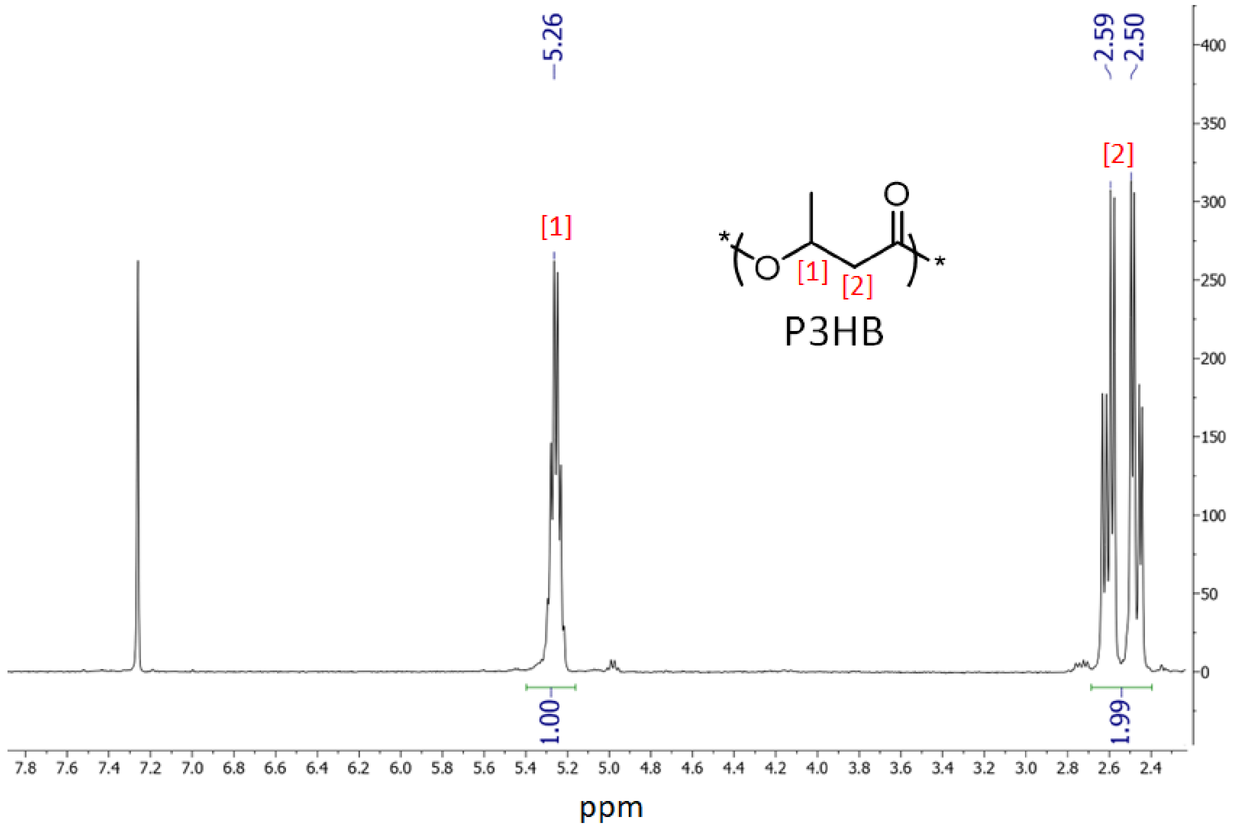


**Figure E2E: [1]H-NMR in $CDCl_3$ of PHA produced by *C. necator* H16 Δ*phaC1* pCM66Tp$_{BAD}$:*hadA-phaC1437*.**

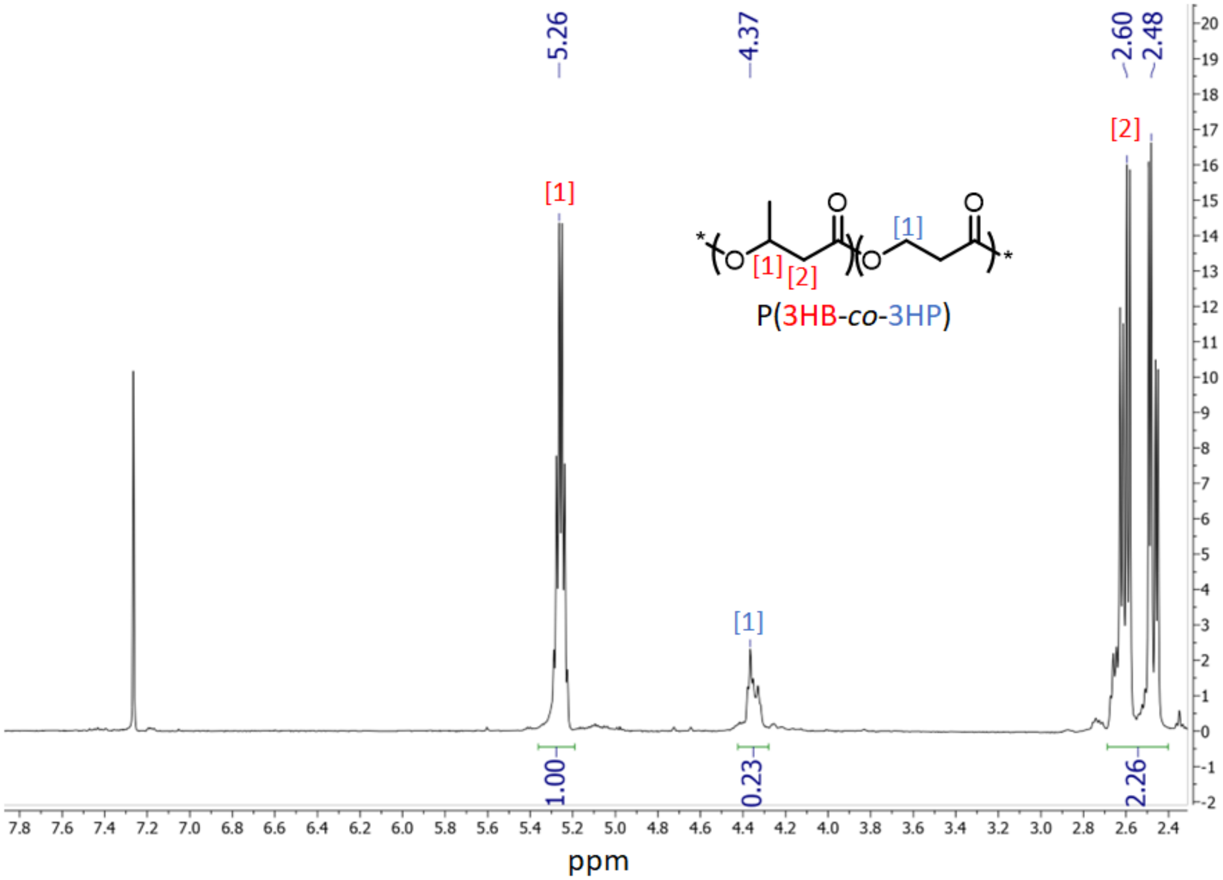


**Figure E2F: $^{1}$H-NMR in CDCl$_3$ of PHA produced by *C. necator* H16 Δ*phaC1* pCM66Tp$_{BAD}$:*pct540-phaC1437* co-fed with 3-hydroxypropionate.**

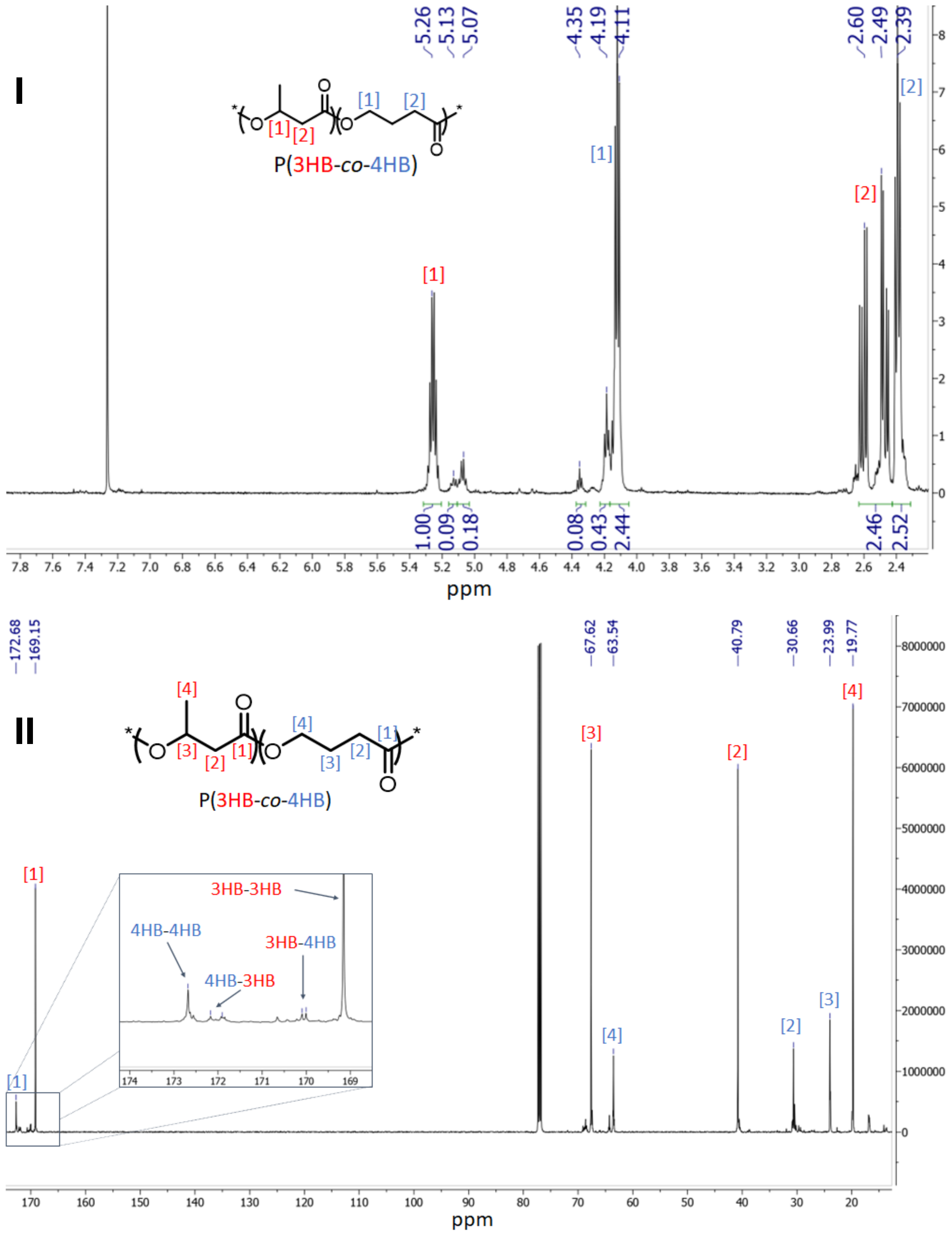


**Figure E2G: $^{1}$H-NMR (I) and $^{13}$C-NMR (II) in $CDCl_3$ of PHA produced by *C. necator* H16 Δ*phaC1* pCM66Tp$_{BAD}$:*pct540-phaC1437* co-fed with 4-hydroxybutyrate.**

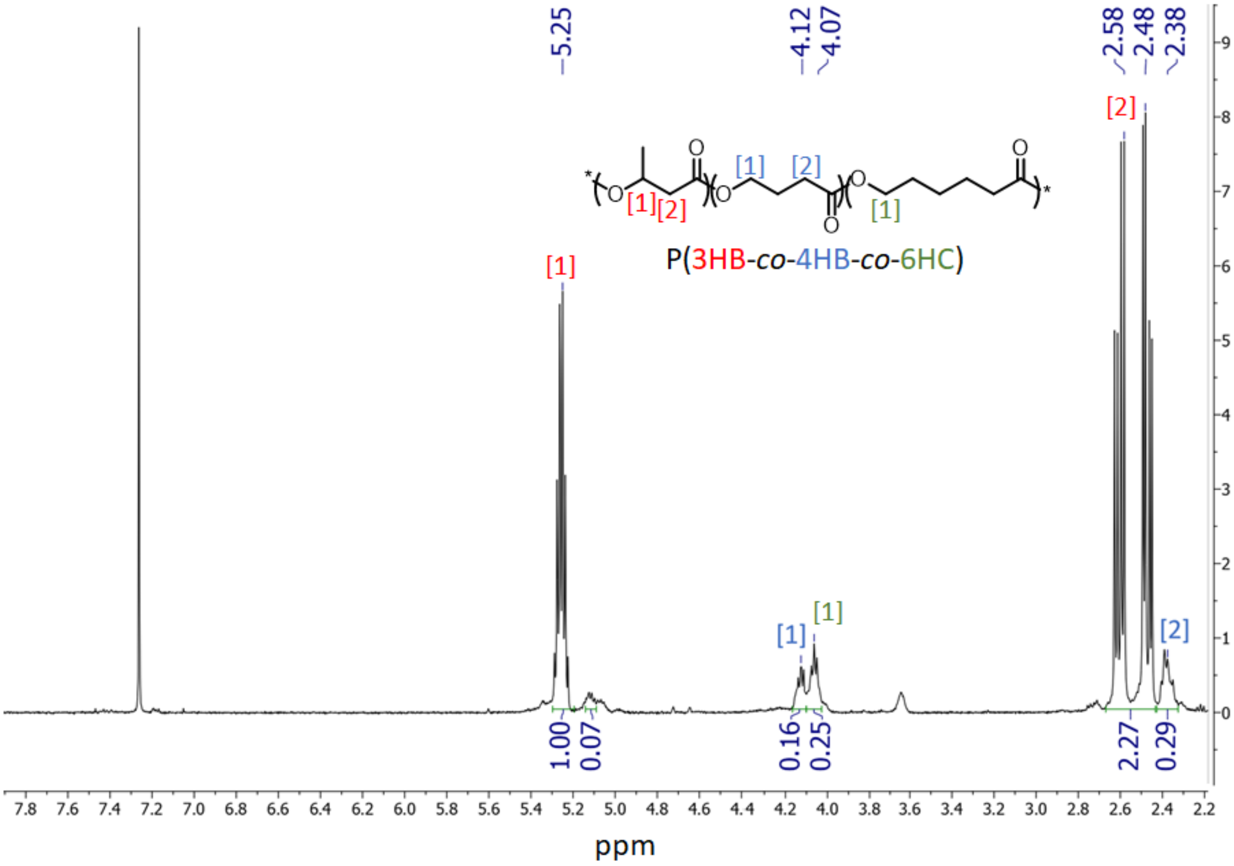


**Figure E2H: $^1$H-NMR in $CDCl_3$ of PHA produced by *C. necator* H16 Δ*phaC1* pCM66Tp$_{BAD}$:*pct540-phaC1437* co-fed with 6-hydroxycaproate.**

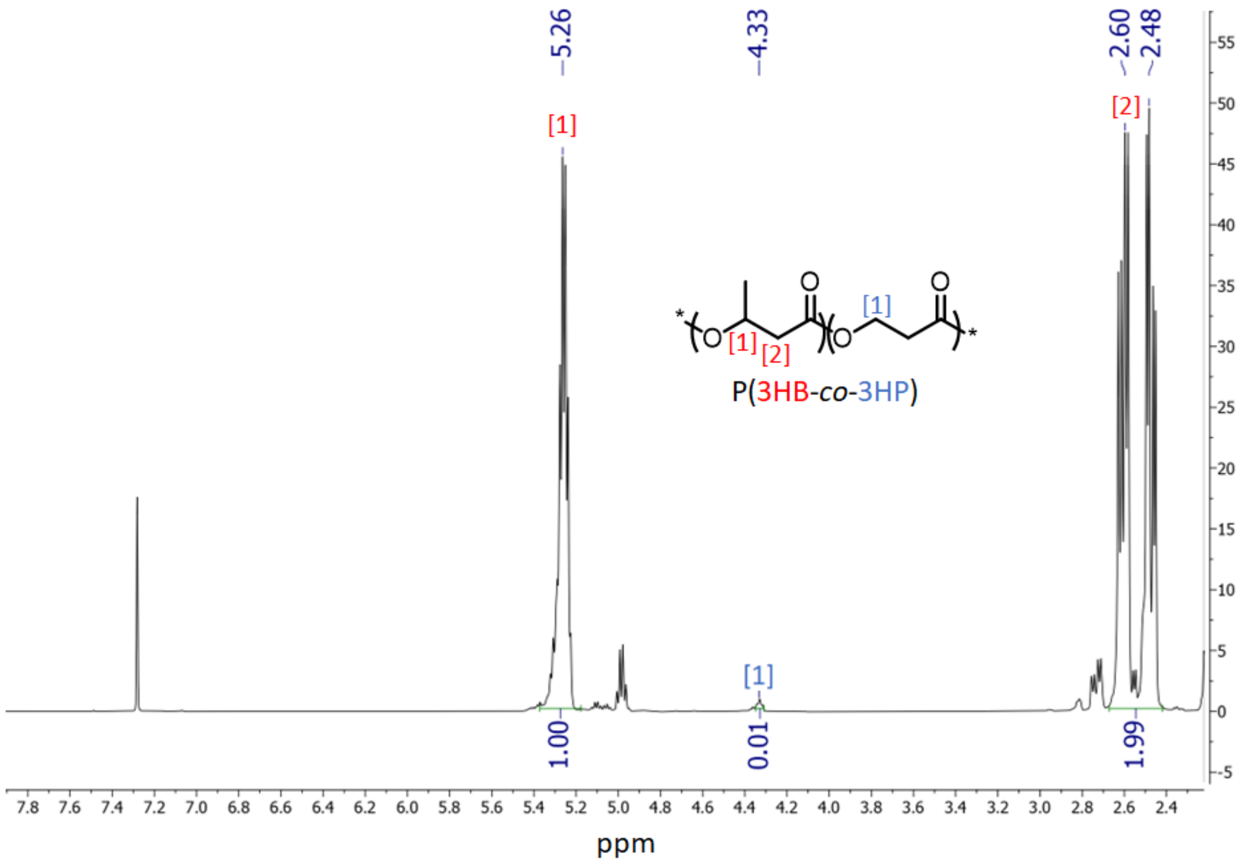


**Figure E2I: $^{1}$H-NMR in $CDCl_3$ of PHA produced by *C. necator* H16 Δ*phaC1* pCM66Tp$_{BAD}$:*hadA-phaC1437* co-fed with 3-hydroxypropionate.**

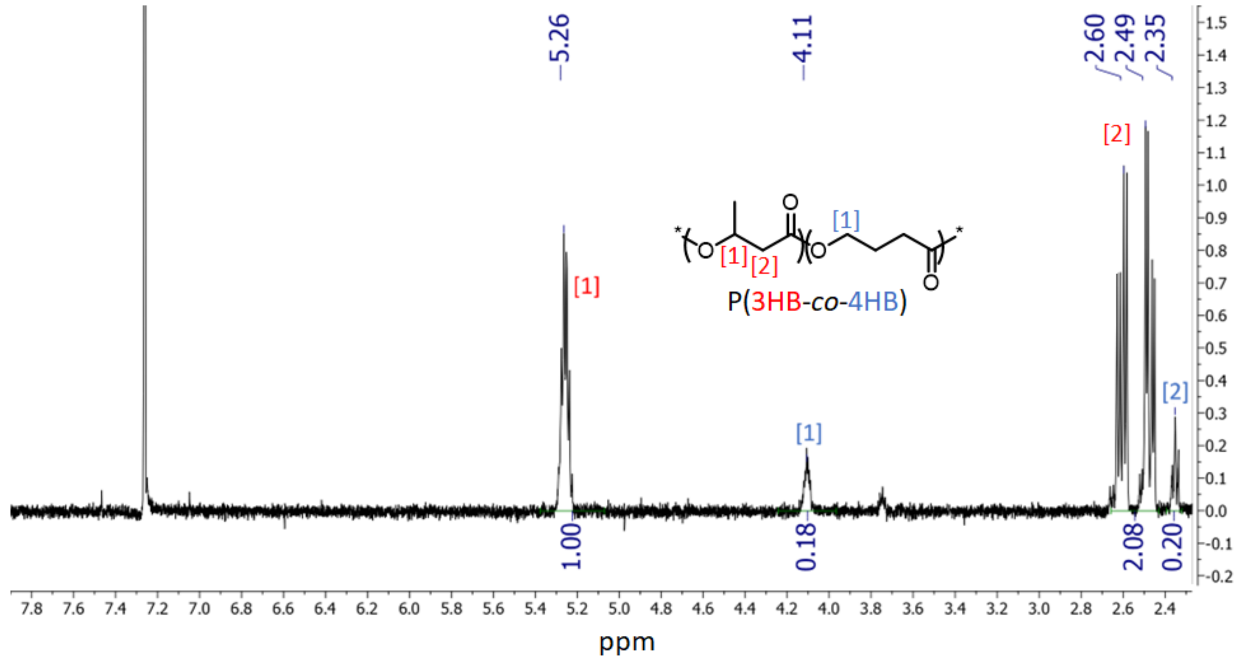


**Figure E2J: $^{1}$H-NMR in $CDCl_3$ of PHA produced by *C. necator* H16 Δ*phaC1* pCM66Tp$_{BAD}$:*hadA-phaC1437* co-fed with 4-hydroxybutyrate.**

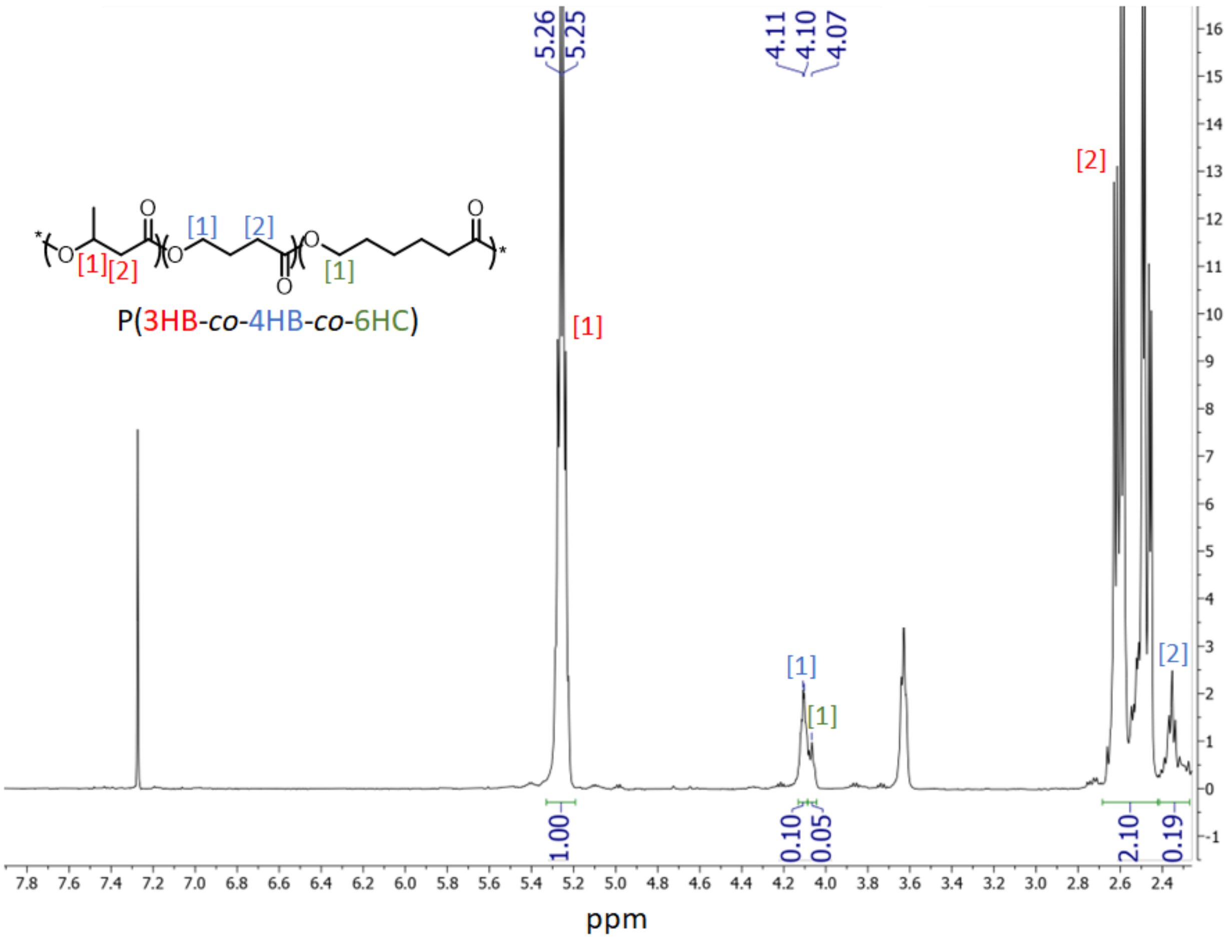


**Figure E2K: $^{1}$H-NMR in $CDCl_3$ of PHA produced by *C. necator* H16 Δ*phaC1* pCM66Tp$_{BAD}$:*hadA-phaC1437* co-fed with 6-hydroxycaproate.**

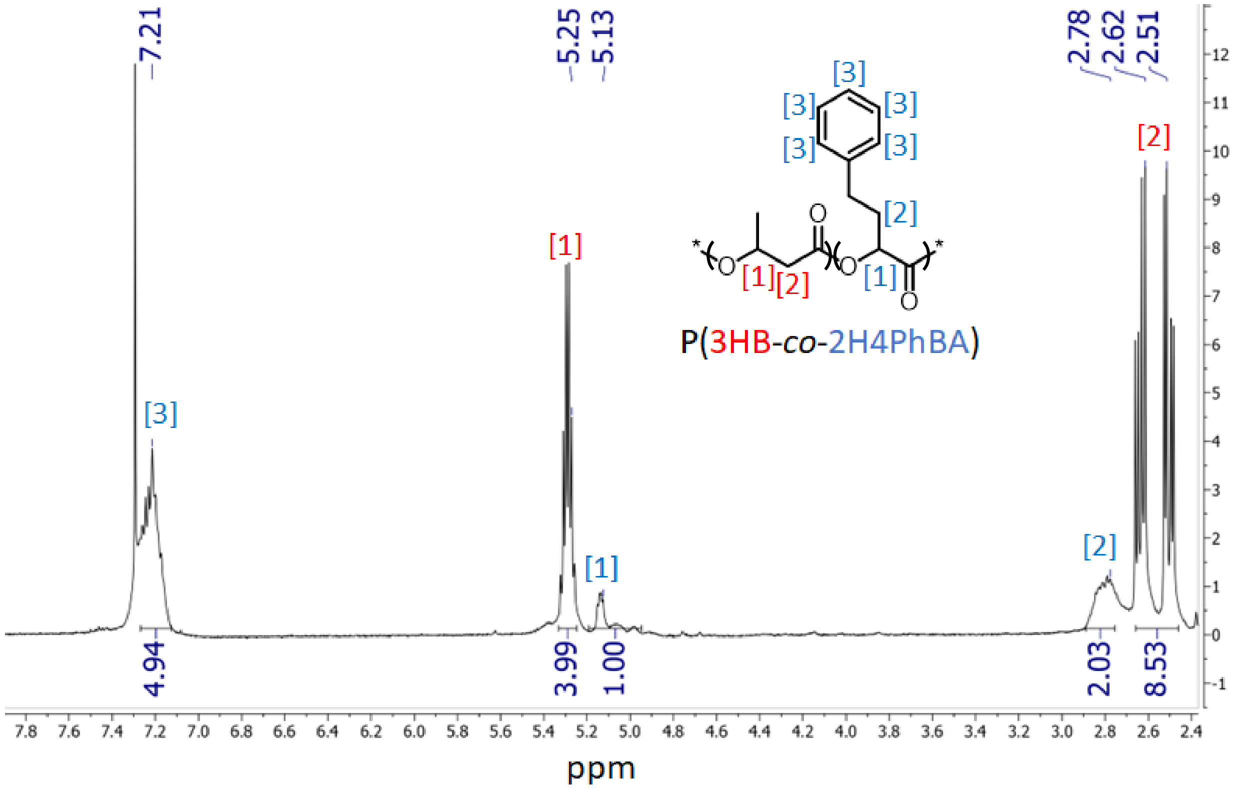


**Figure E2L: $^{1}$H-NMR in $CDCl_3$ of PHA produced by *C. necator* H16 Δ*phaC1* pCM66Tp$_{BAD}$:*hadA-phaC1437* co-fed with 2-hydroxy-4-phenylbutyrate.**

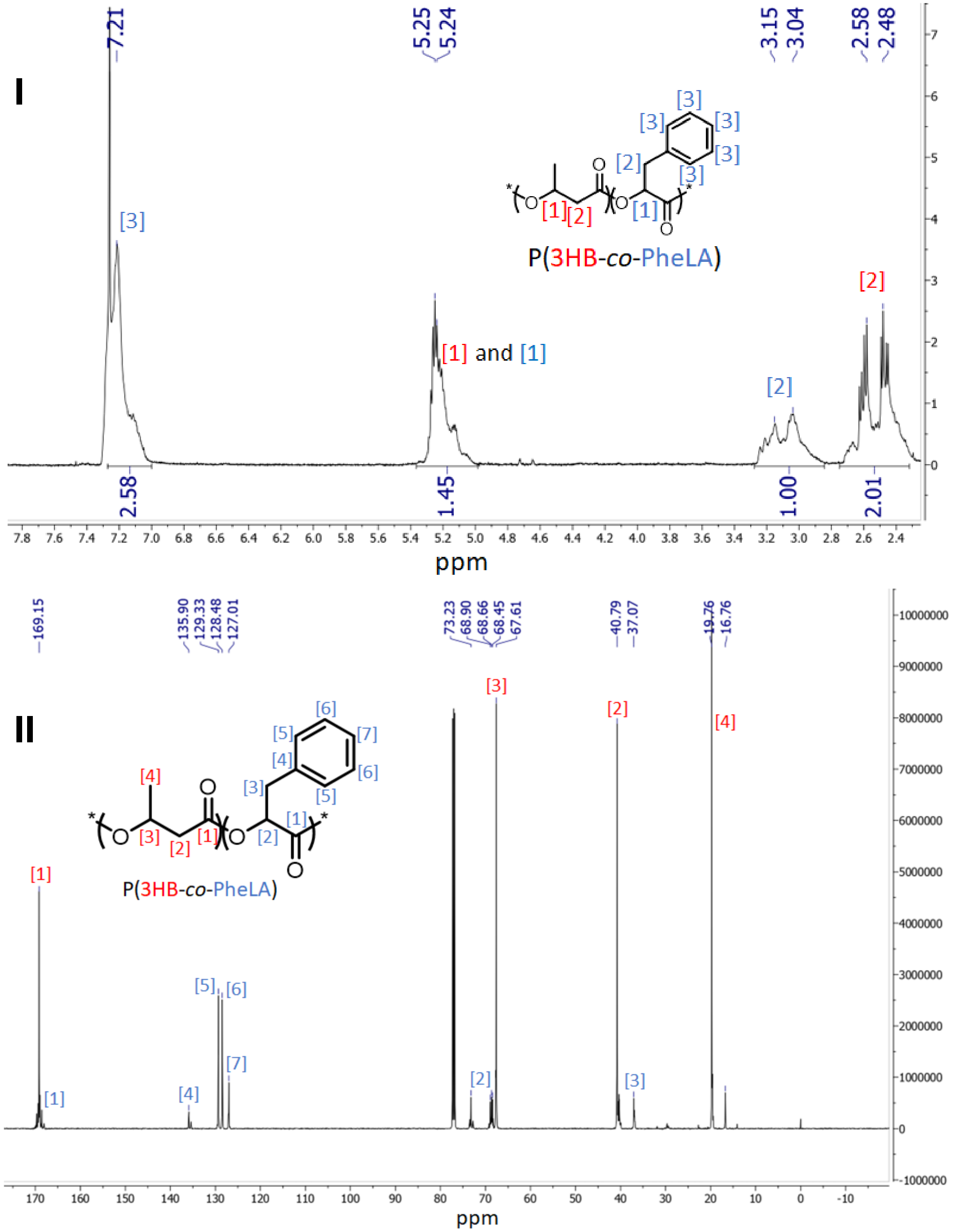


**Figure E2M: $^{1}$H-NMR (I) and $^{13}$C-NMR (II) in $CDCl_3$ of PHA produced by *C. necator* H16 Δ*phaC1* pCM66Tp$_{BAD}$:*hadA-phaC1437* co-fed with phenyllactate.**

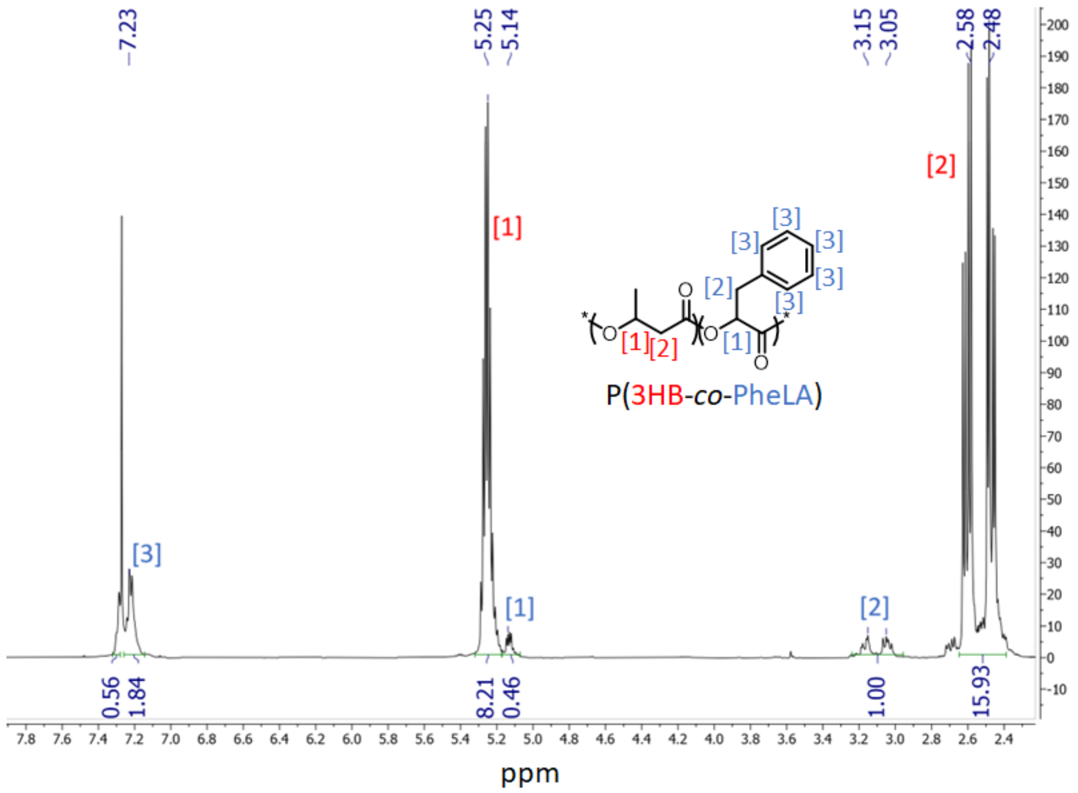


**Figure E2N: [1]H-NMR in $CDCl_3$ of PHA produced by *C. necator* H16 Δ*phaC1* pCM66Tp$_{BAD}$:*hadA-phaC1437* co-fed with phenyllactate in bio-electrochemical system.**

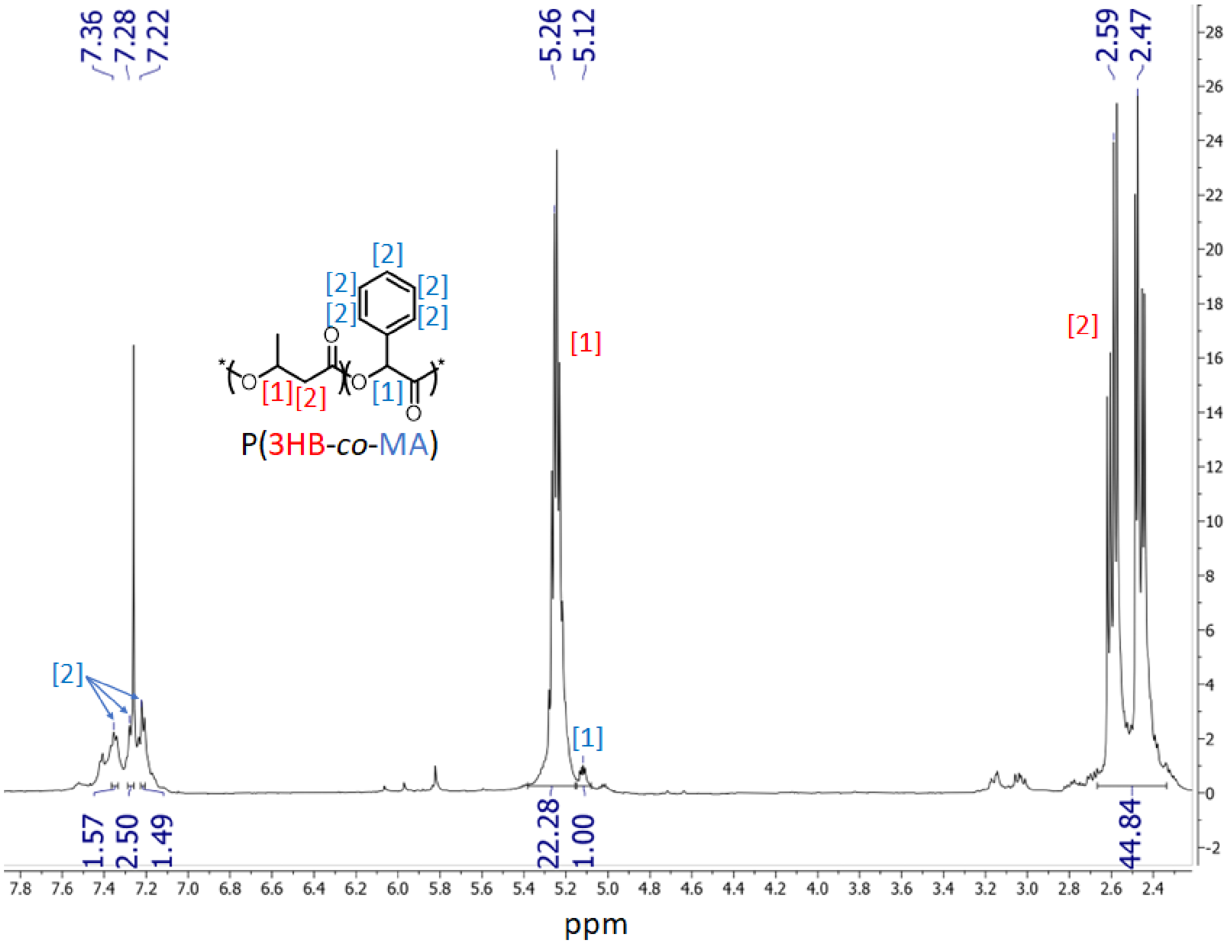


**Figure E2O: $^{1}$H-NMR in $CDCl_3$ of PHA produced by *C. necator* H16 Δ*phaC1* pCM66Tp$_{BAD}$:*hadA-phaC1437* co-fed with mandelate.**

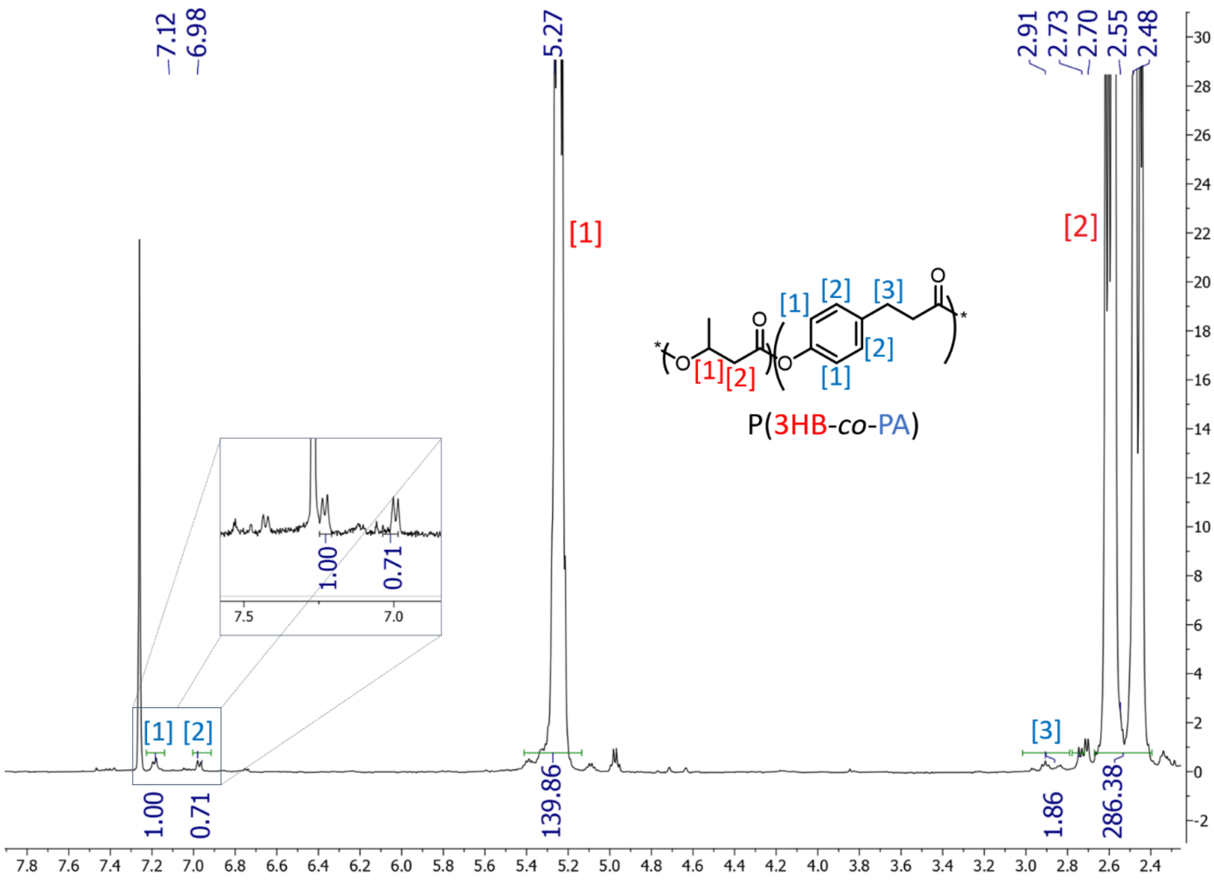


**Figure E2P: $^1$H-NMR in $CDCl_3$ of PHA produced by *C. necator* H16 Δ*phaC1* pCM66Tp$_{BAD}$:*hadA-phaC1437* co-fed with phloretate.**

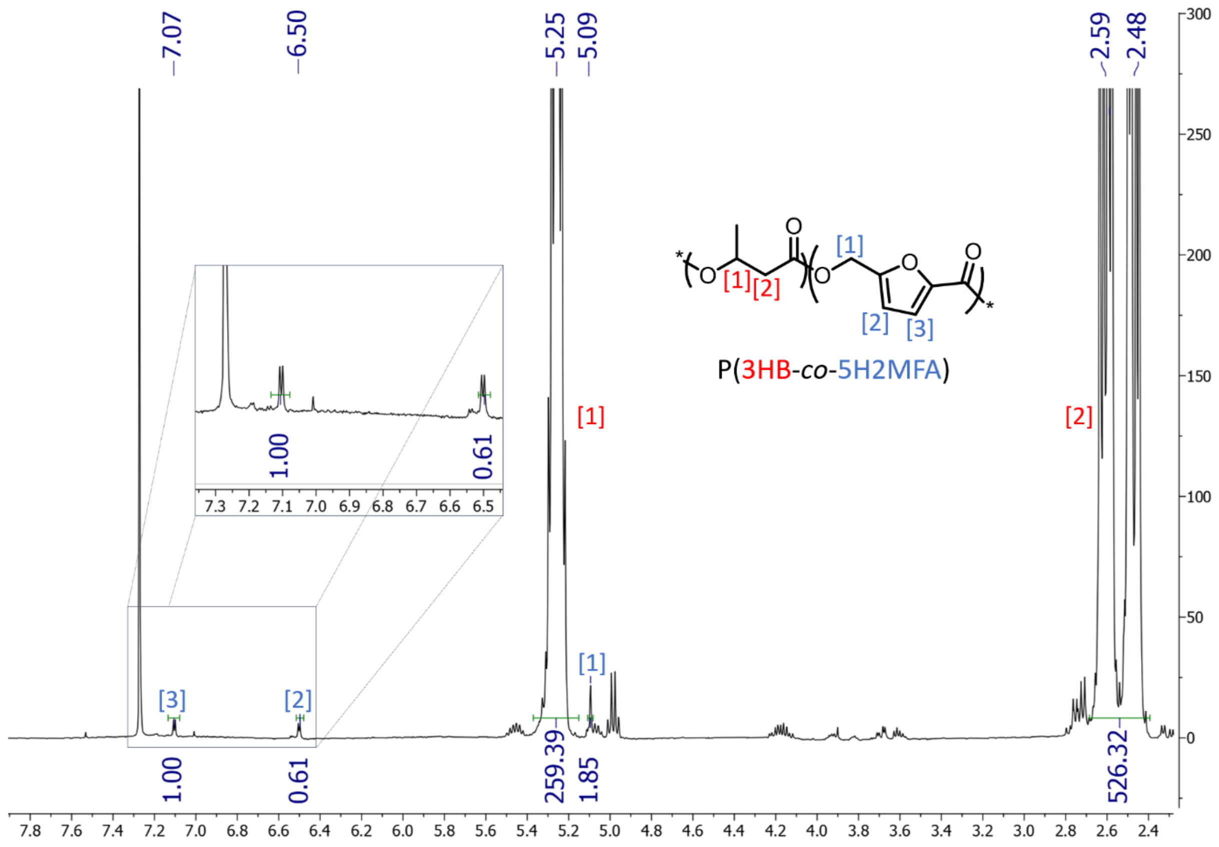


**Figure E2Q: $^1$H-NMR in $CDCl_3$ of PHA produced by *C. necator* H16 Δ*phaC1* pCM66Tp$_{BAD}$:*hadA-phaC1437* co-fed with 5-hydroxy-2-methylfuranoate.**

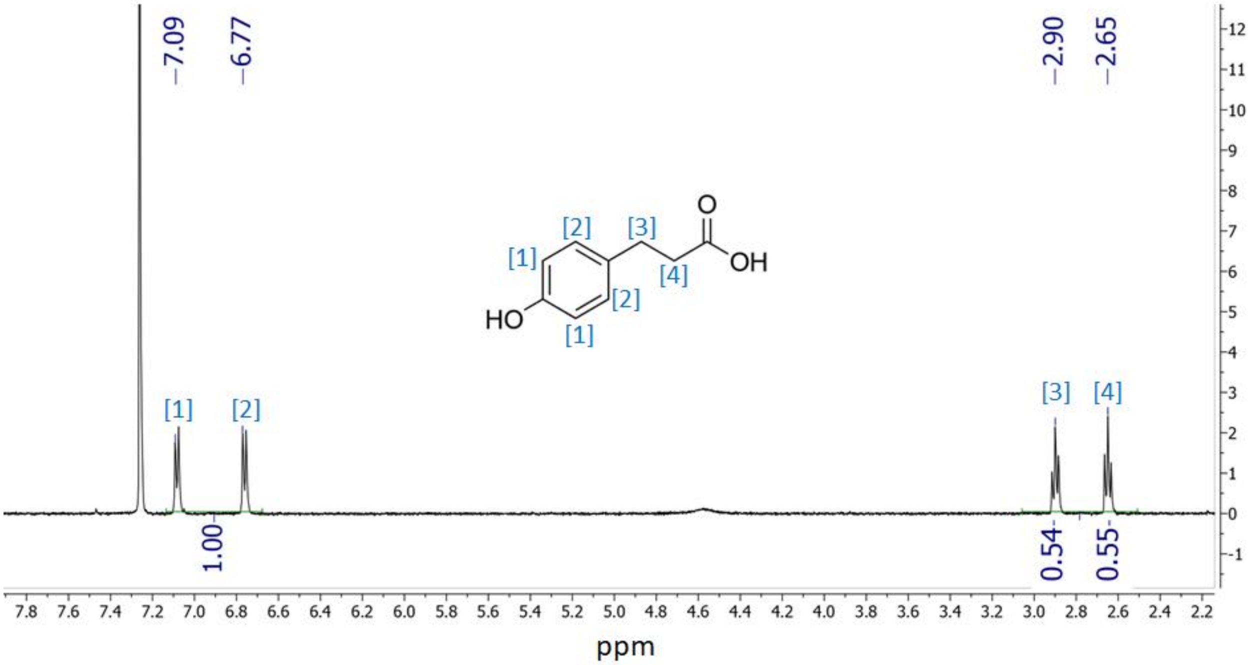


**Figure E2R: $^1$H-NMR of phloretic acid in deuterated trifluoroacetic acid.**

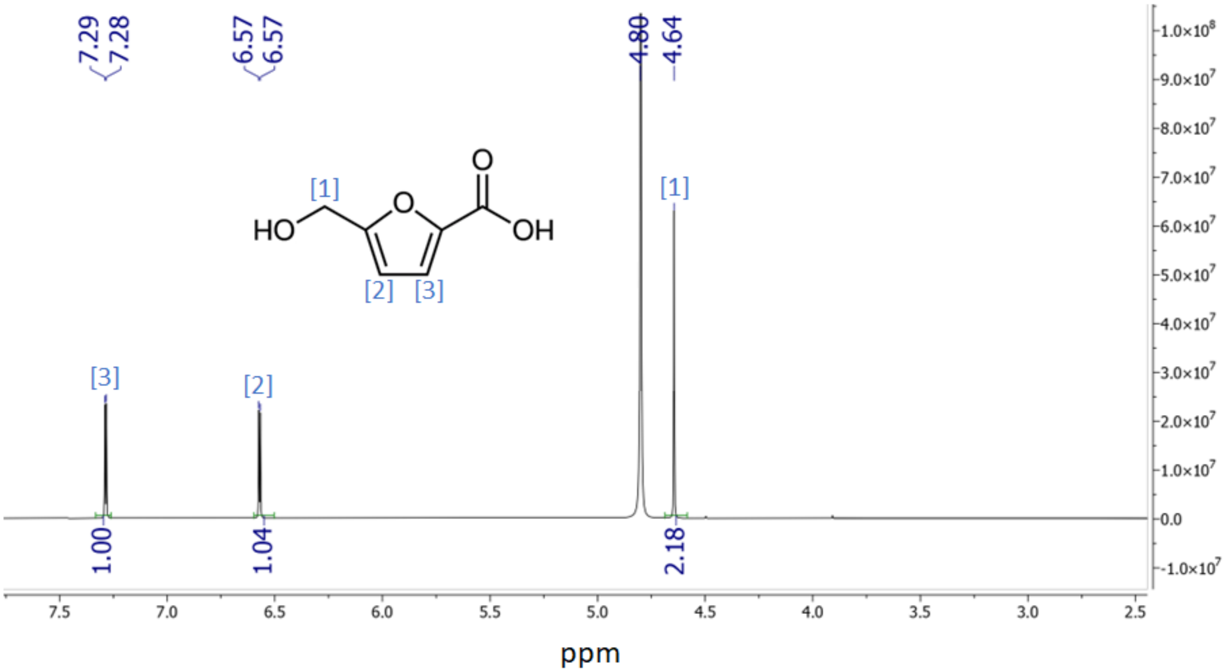


**Figure E2S: $^1$H-NMR of 5-hydroxy-2-methylfuranoic acid in deuterated water.**
