## Extended Data 3 for "Novel Aromatic Polyhydroxyalkanoates from Engineered *Cupriavidus necator* H16: Expanding Bio-Polyester Synthesis from Renewable Feedstocks"

<u>GPC traces of bio-polyesters formed by wild-type and engineered *C. necator* H16</u>

**A**

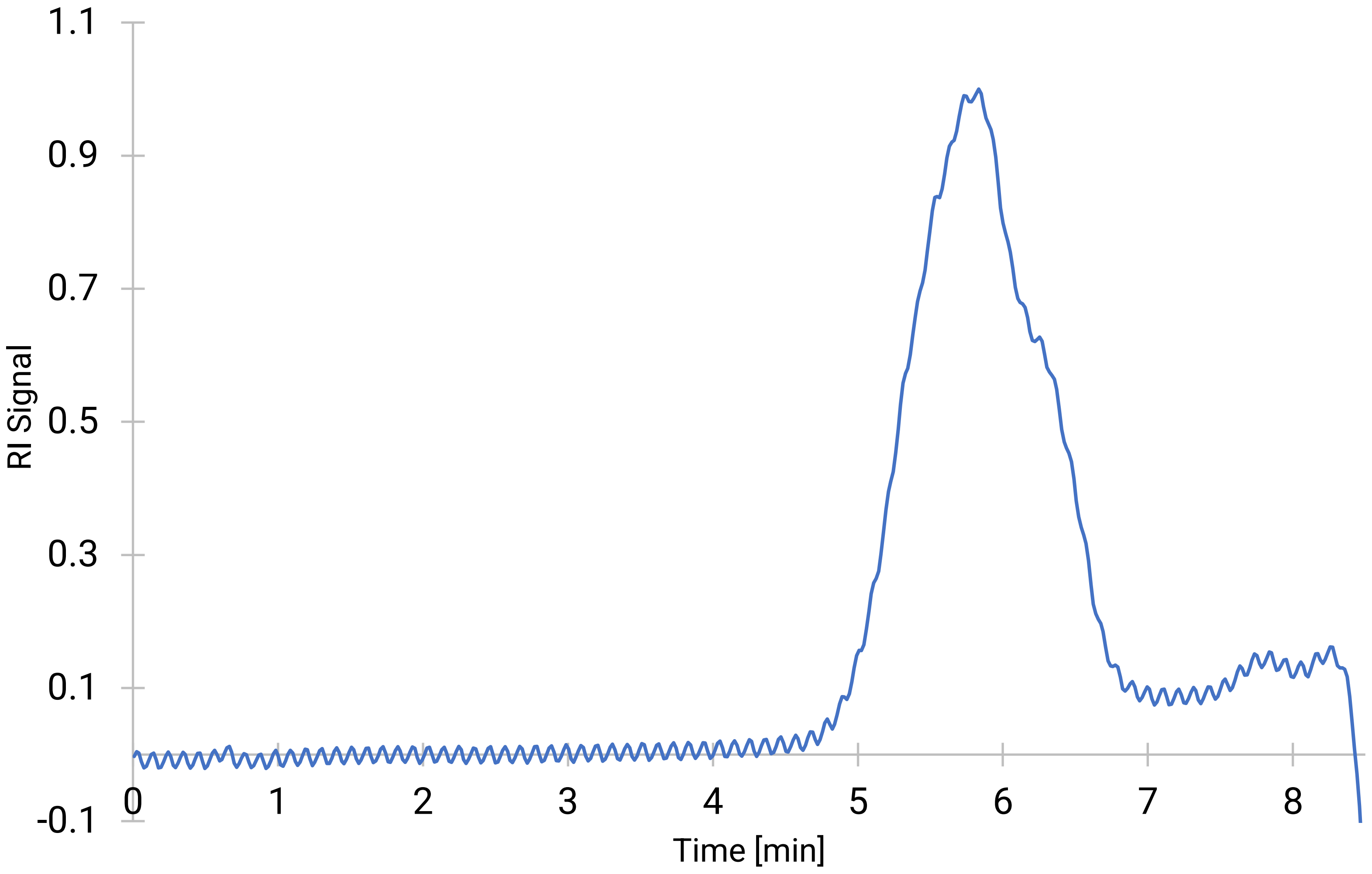


**Figure E4A: GPC plot of bio-polyester obtained from *C. necator* H16 (wild-type), as per table S2.**

**B**

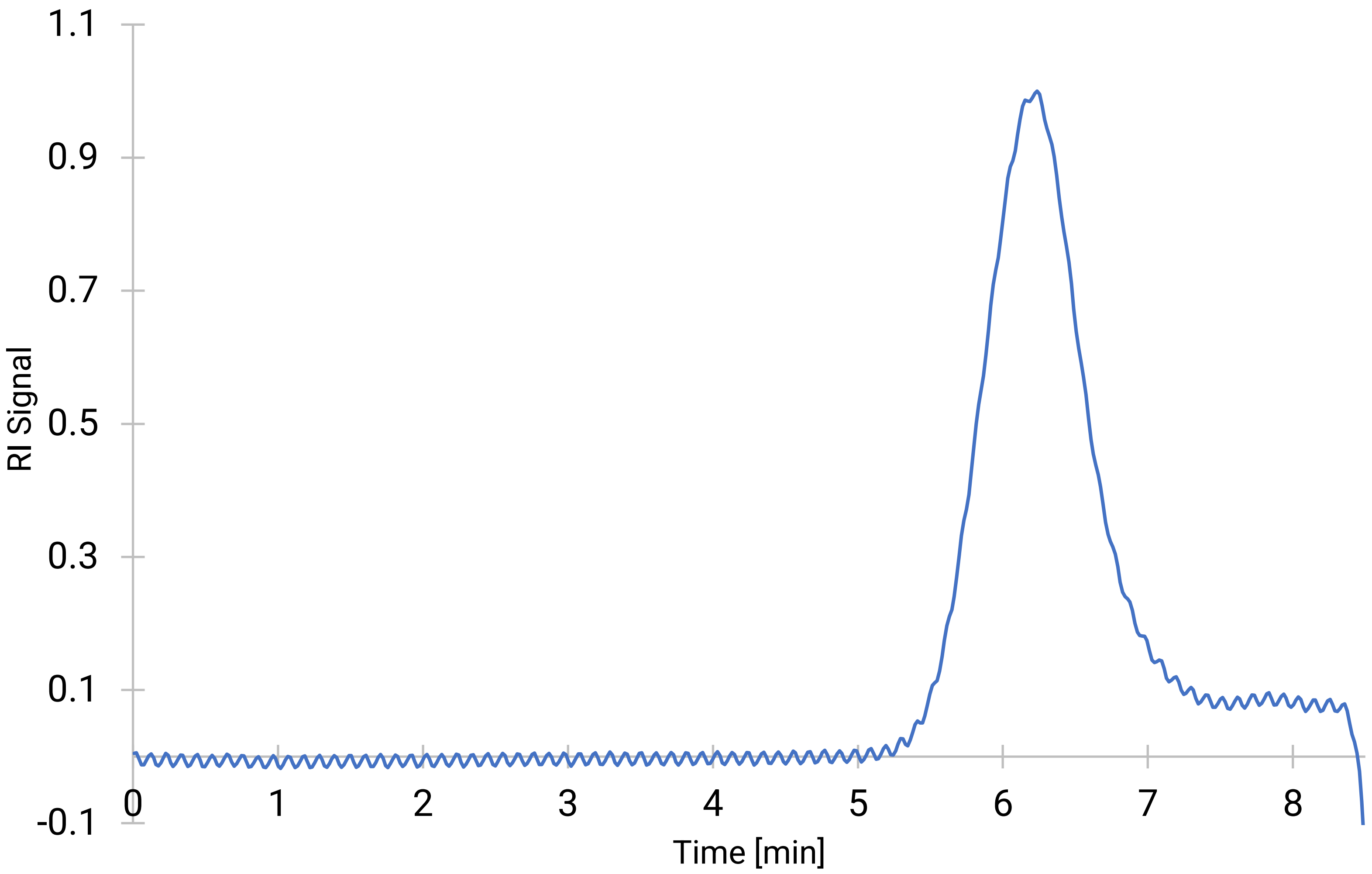


**Figure E4B: GPC plot of bio-polyester obtained from *C. necator* H16 Δ*phaC1* pCM66Tp$_{BAD}$:*pct540-phaC1437* when not induced, as per table S2.**

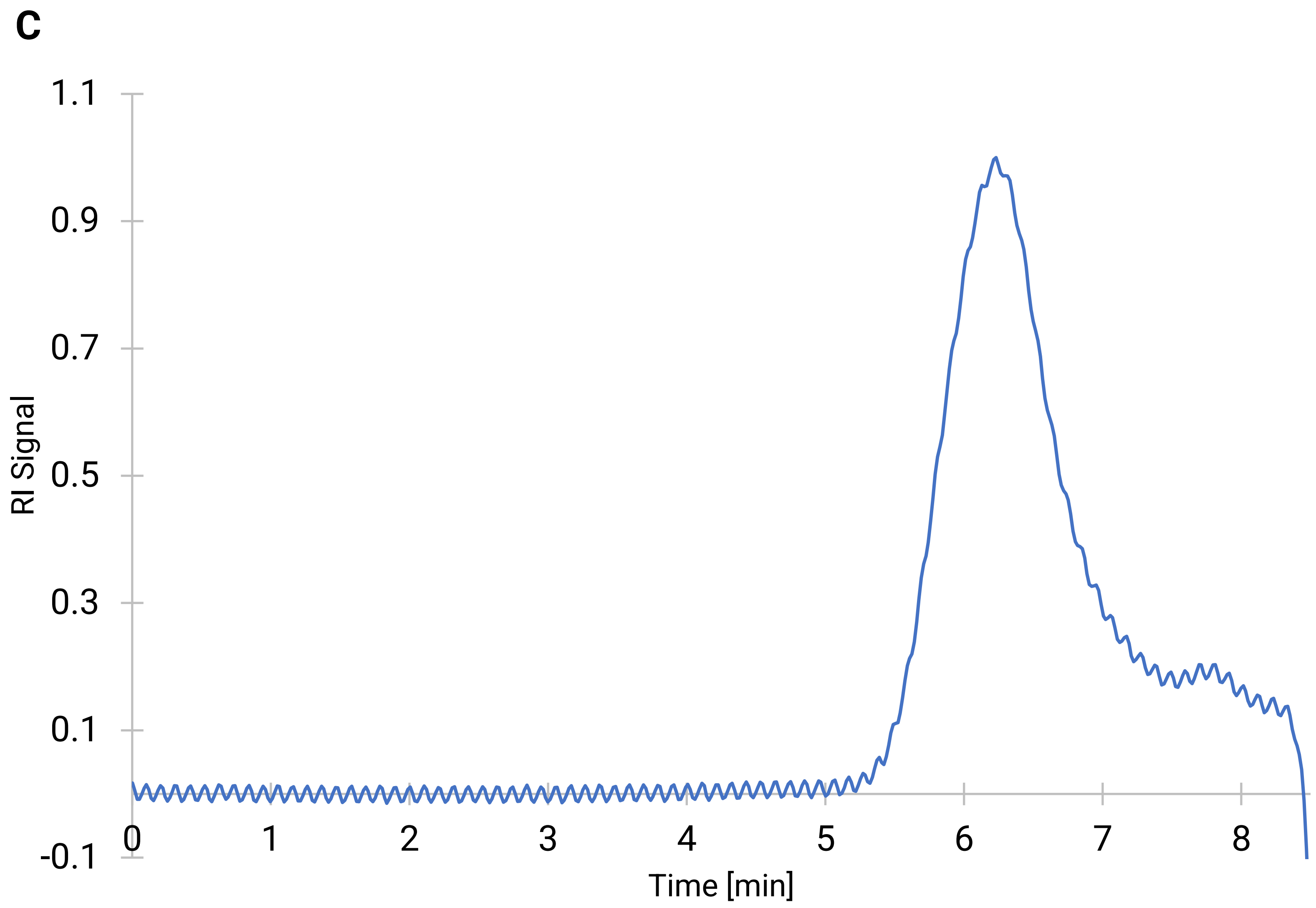


**Figure E4C: GPC plot of bio-polyester obtained from *C. necator* H16 Δ*phaC1* pCM66Tp$_{BAD}$:*pct540-phaC1437* when induced, as per table S2.**

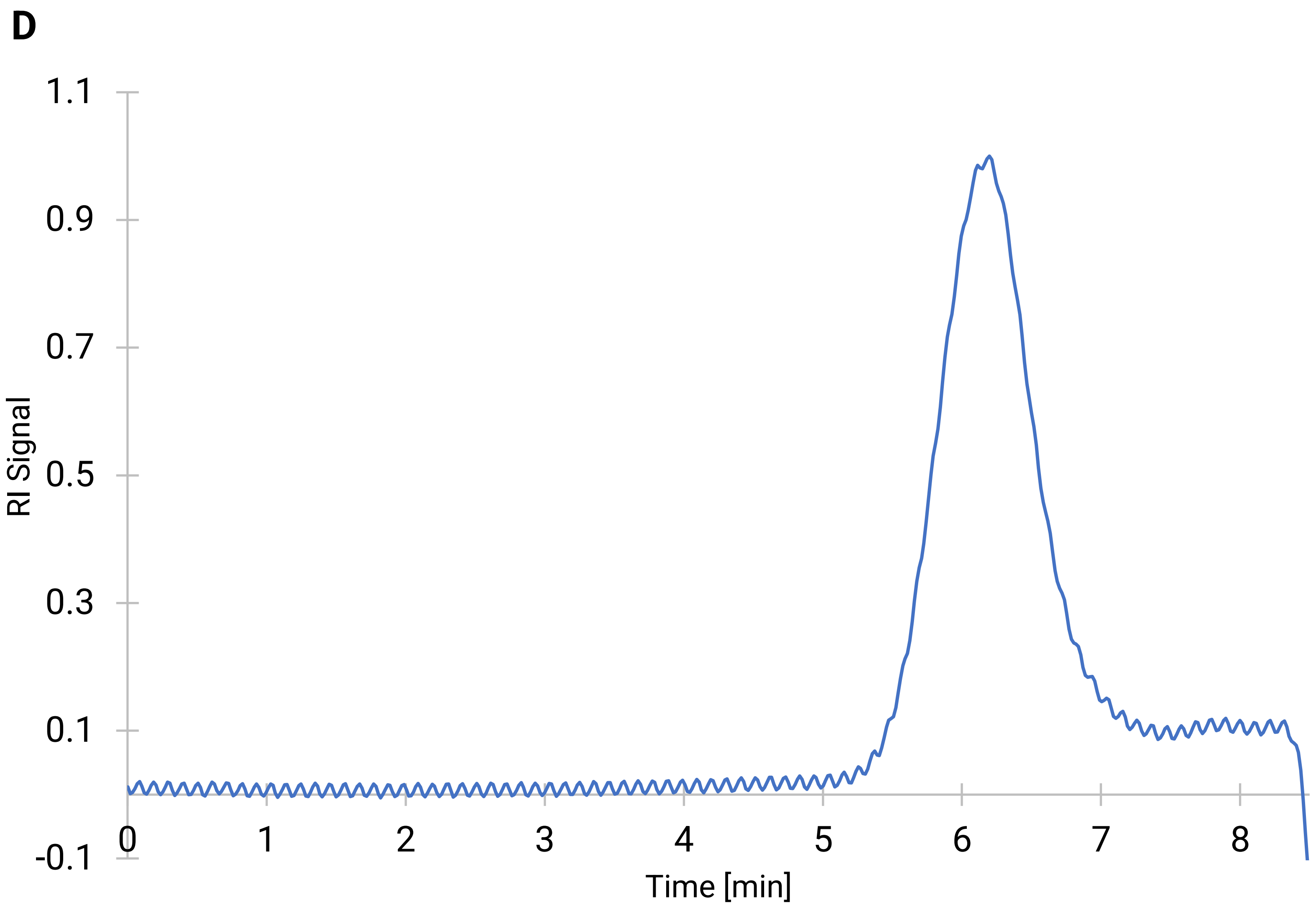


**Figure E4D: GPC plot of bio-polyester obtained from *C. necator* H16 Δ*phaC1* pCM66Tp$_{BAD}$:*hadA-phaC1437* when not induced, as per table S2.**

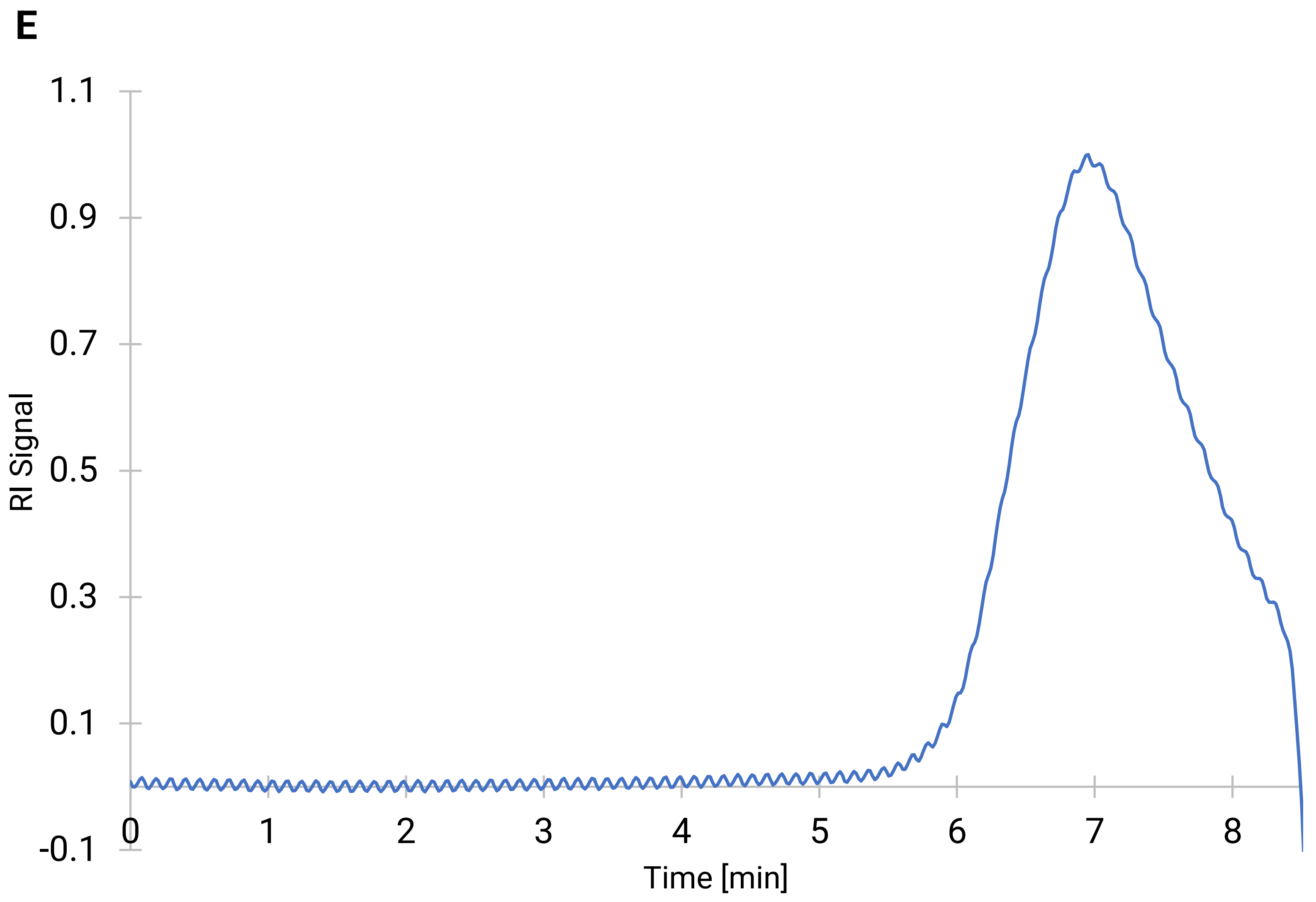


**Figure E4E: GPC plot of bio-polyester obtained from *C. necator* H16 Δ*phaC1* pCM66Tp$_{BAD}$:*hadA-phaC1437* when induced, as per table S2.**

**F**

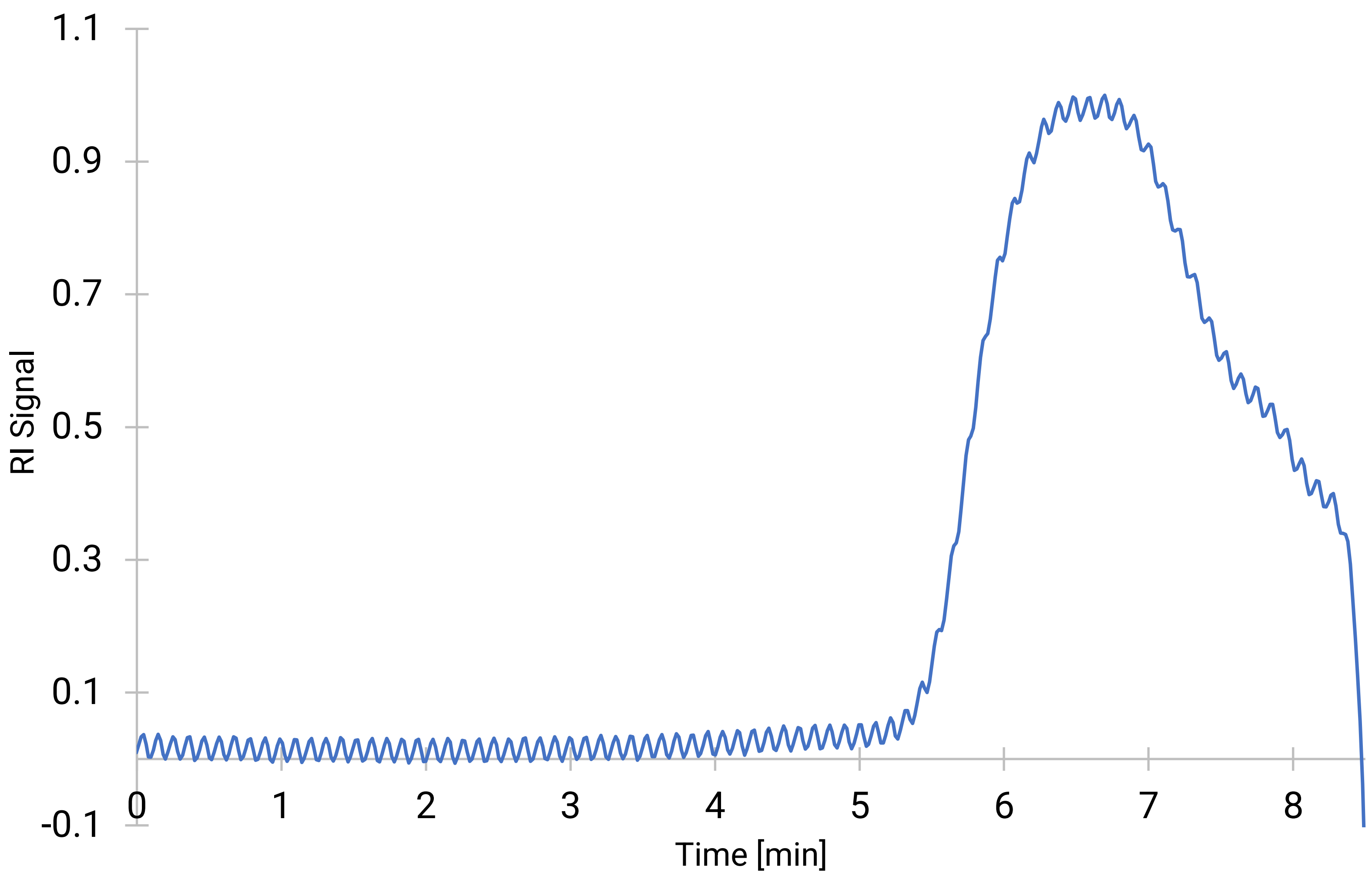


**Figure E4F: GPC plot of bio-polyester obtained from *C. necator* H16 Δ*phaC1* pCM66Tp$_{BAD}$:*pct540-phaC1437* when induced and co-fed with 4-hydroxybutyric acid, as per table S2.**

**G**

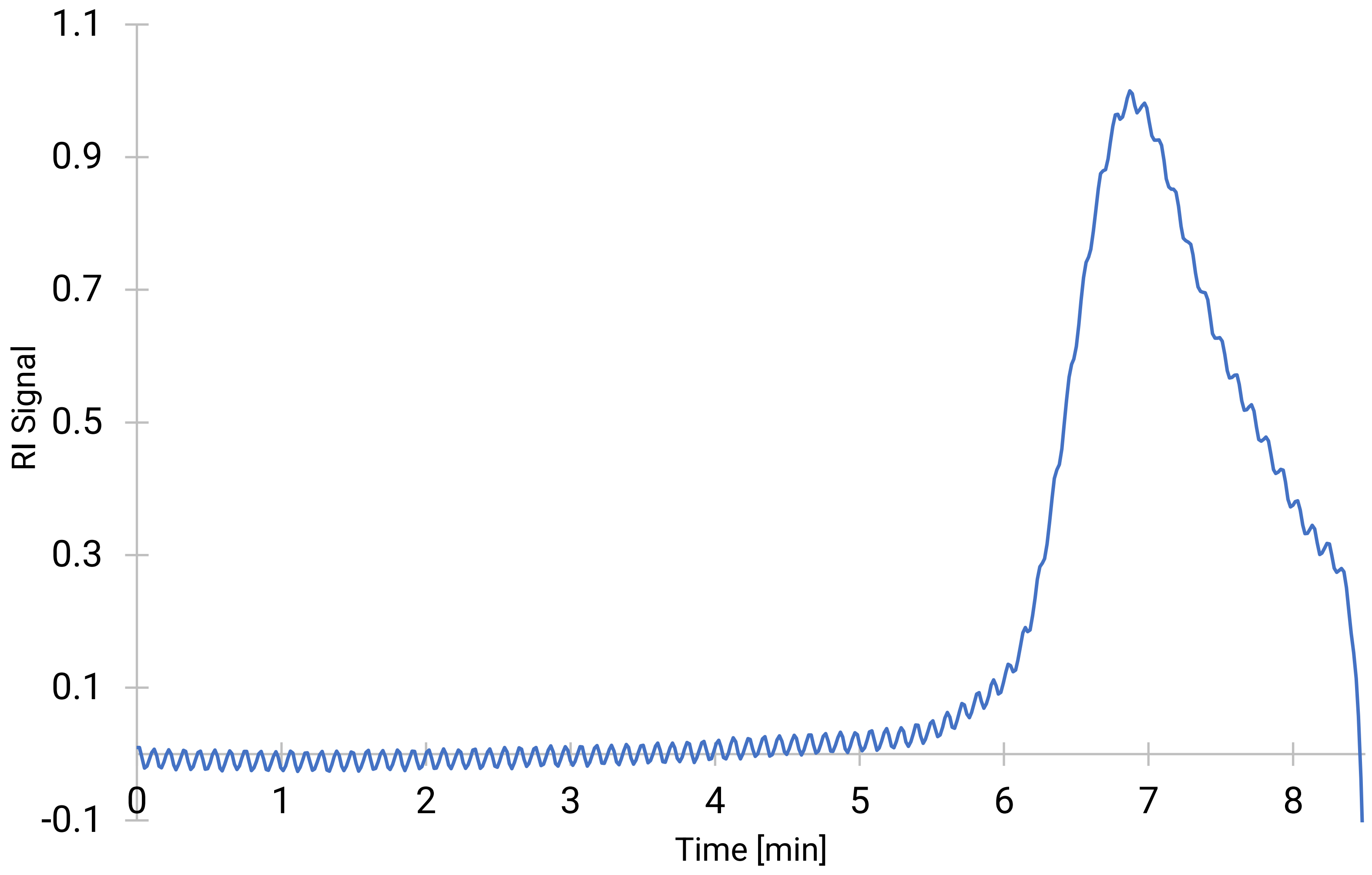


**Figure E4G: GPC plot of bio-polyester obtained from *C. necator* H16 Δ*phaC1* pCM66Tp$_{BAD}$:*hadA-phaC1437* when induced and co-fed with 6-hydroxycaproic acid, as per table S2.**

**H**

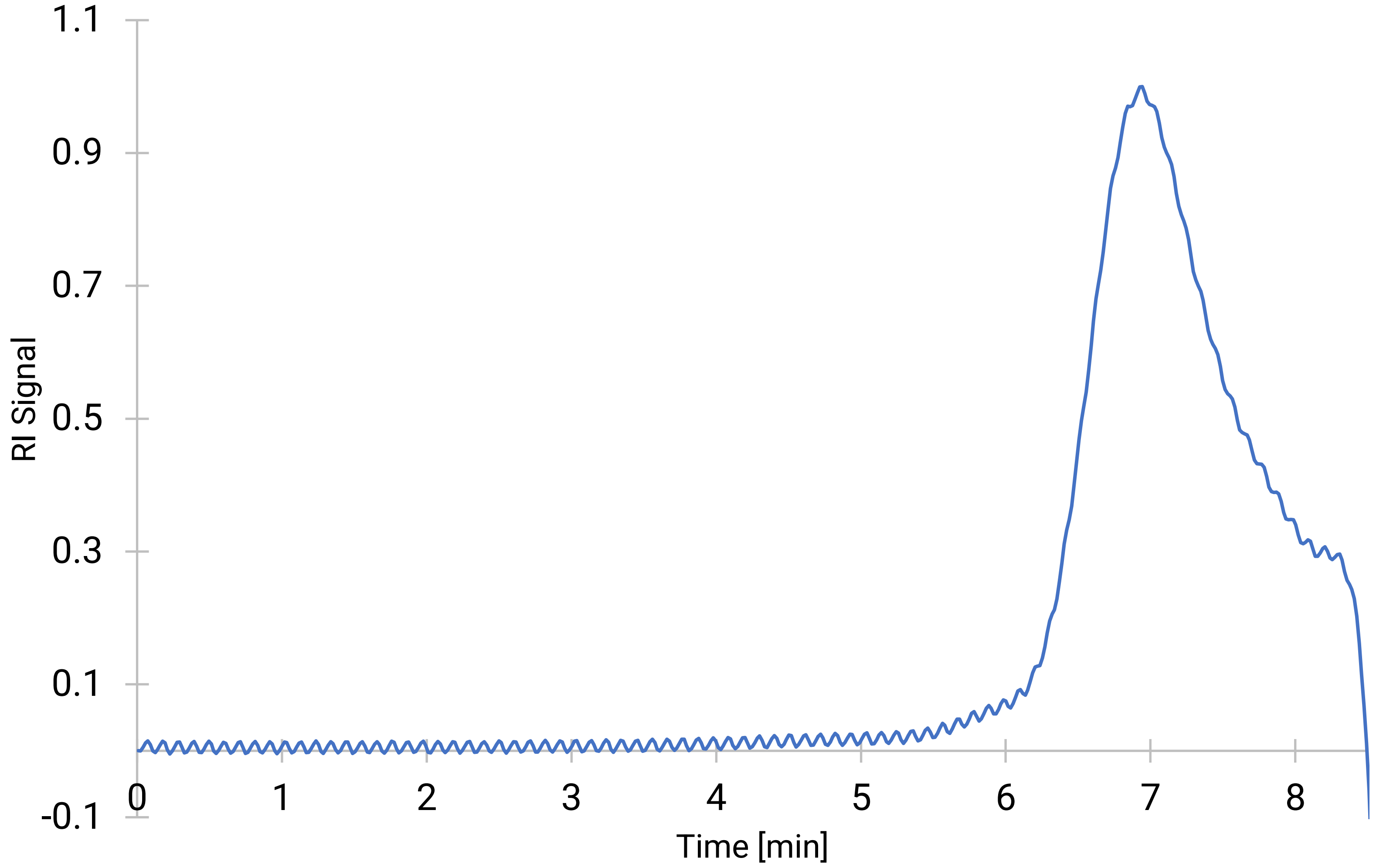


**Figure E4H: GPC plot of bio-polyester obtained from *C. necator* H16 Δ*phaC1* pCM66Tp$_{BAD}$:*hadA-phaC1437* when induced and co-fed with phenyllactic acid, as per table S2.**

**I**

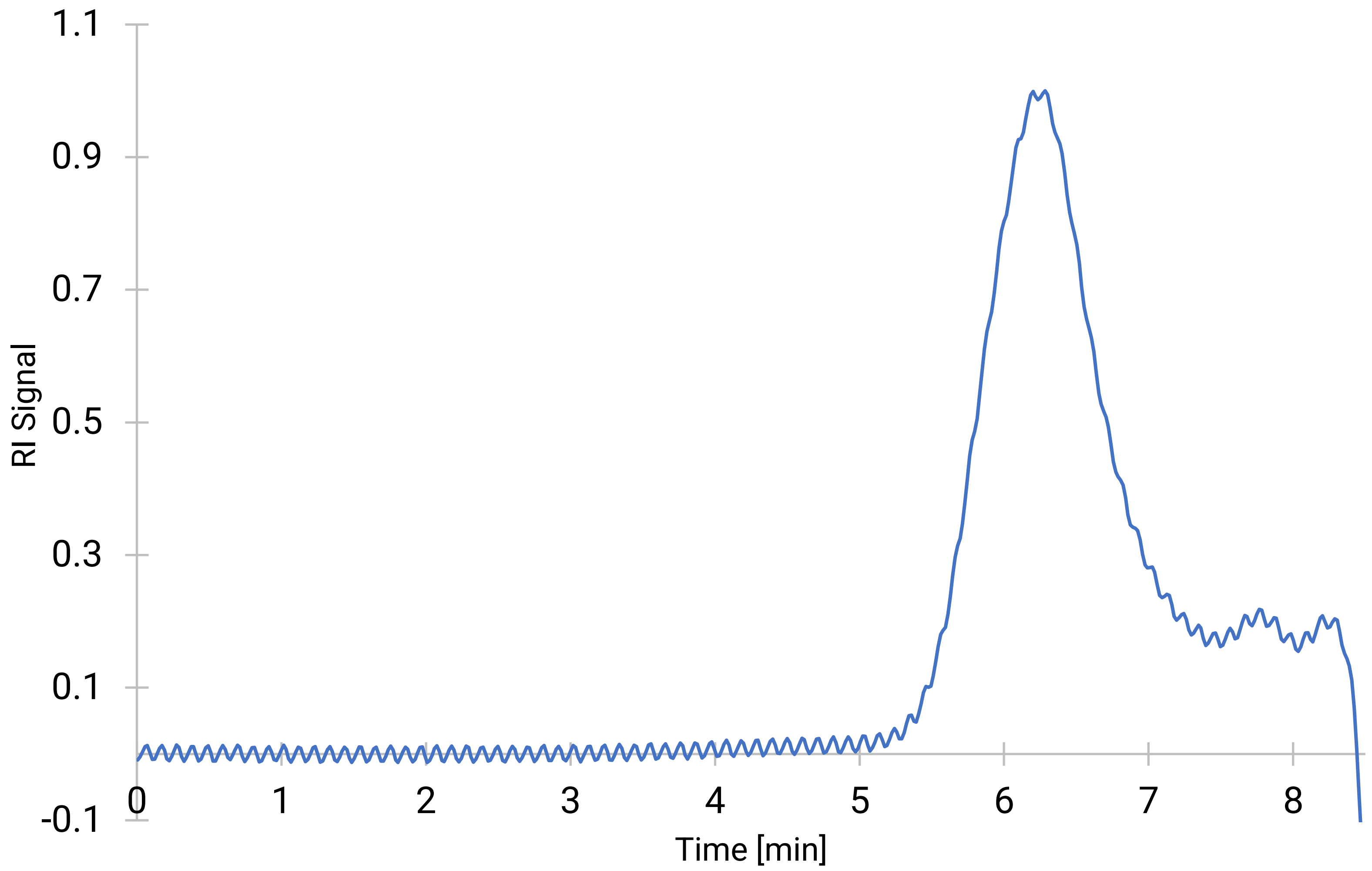


**Figure E4I: GPC plot of bio-polyester obtained from *C. necator* H16 Δ*phaC1* pCM66Tp$_{BAD}$:*hadA-phaC1437* non-induced and co-fed with phloretic acid, as per table S2.**

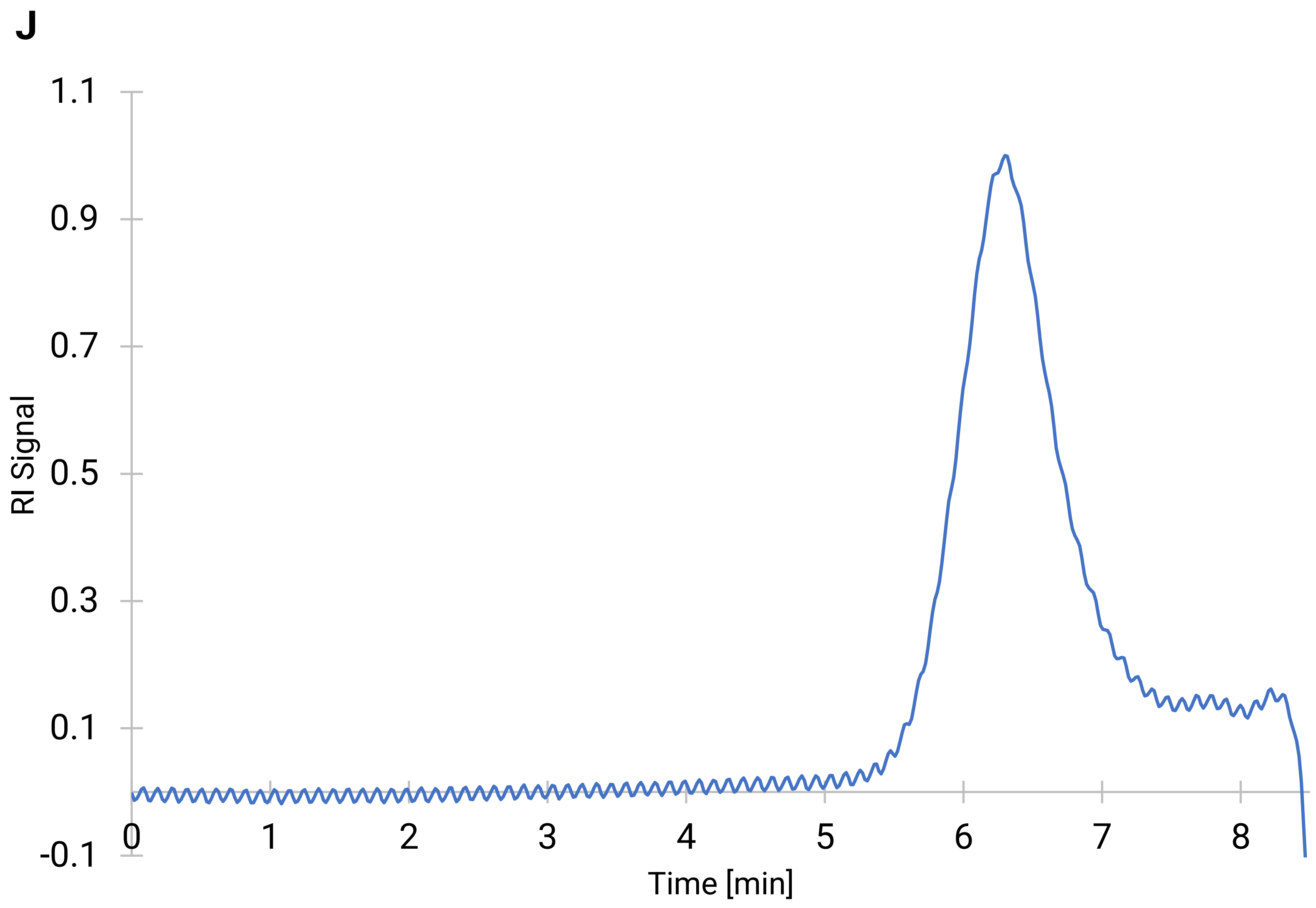


**Figure E4J: GPC plot of bio-polyester obtained from *C. necator* H16 Δ*phaC1* pCM66Tp$_{BAD}$:*hadA-phaC1437* non-induced and co-fed with phloretic acid, as per table S2.**

**K**

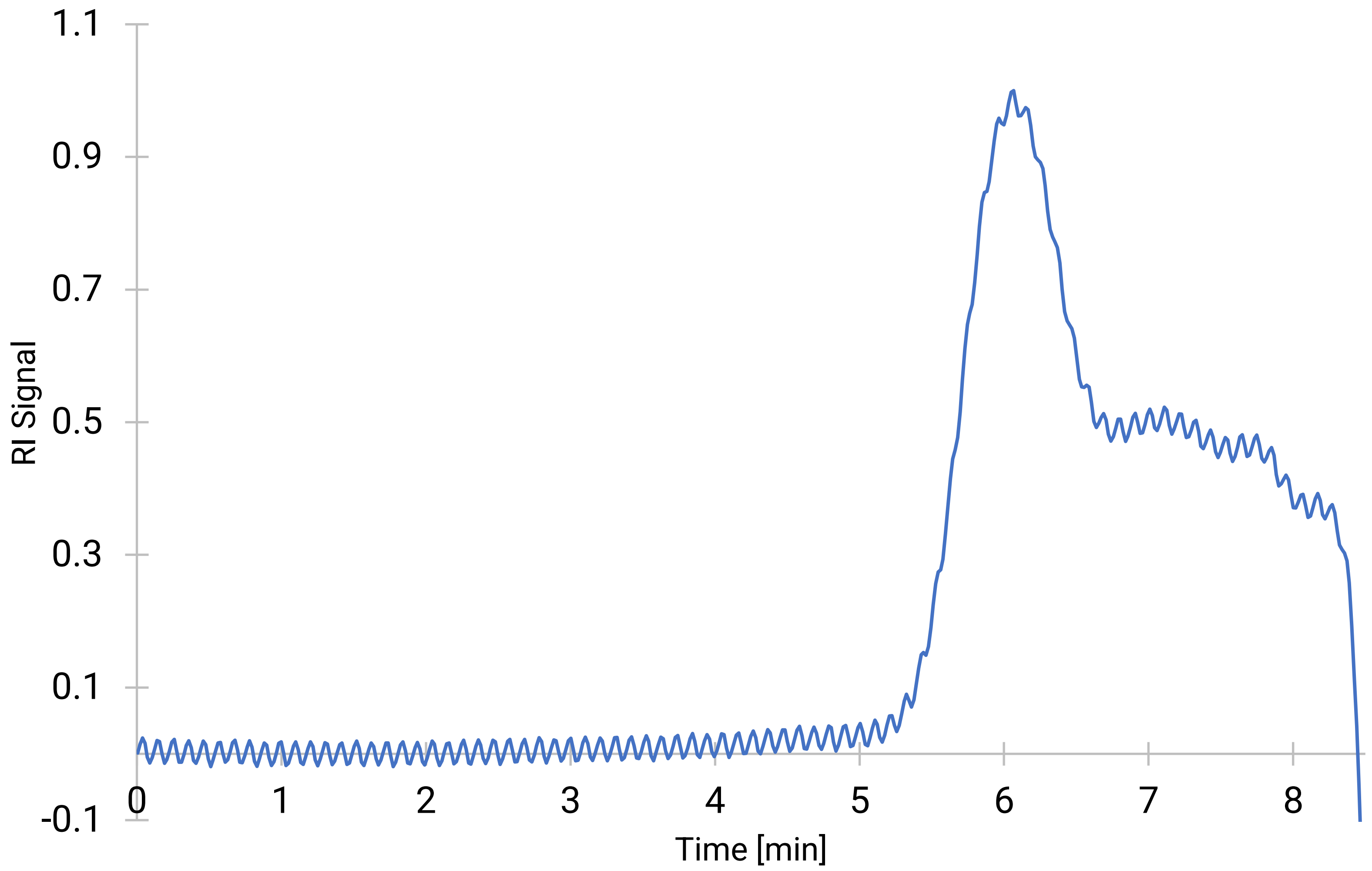


**Figure E4K: GPC plot of bio-polyester obtained from *C. necator* H16 Δ*phaC1* pCM66Tp$_{BAD}$:*hadA-phaC1437* when induced and co-fed with phloretic acid, harvested early-on in cultivation, as per table S2.**

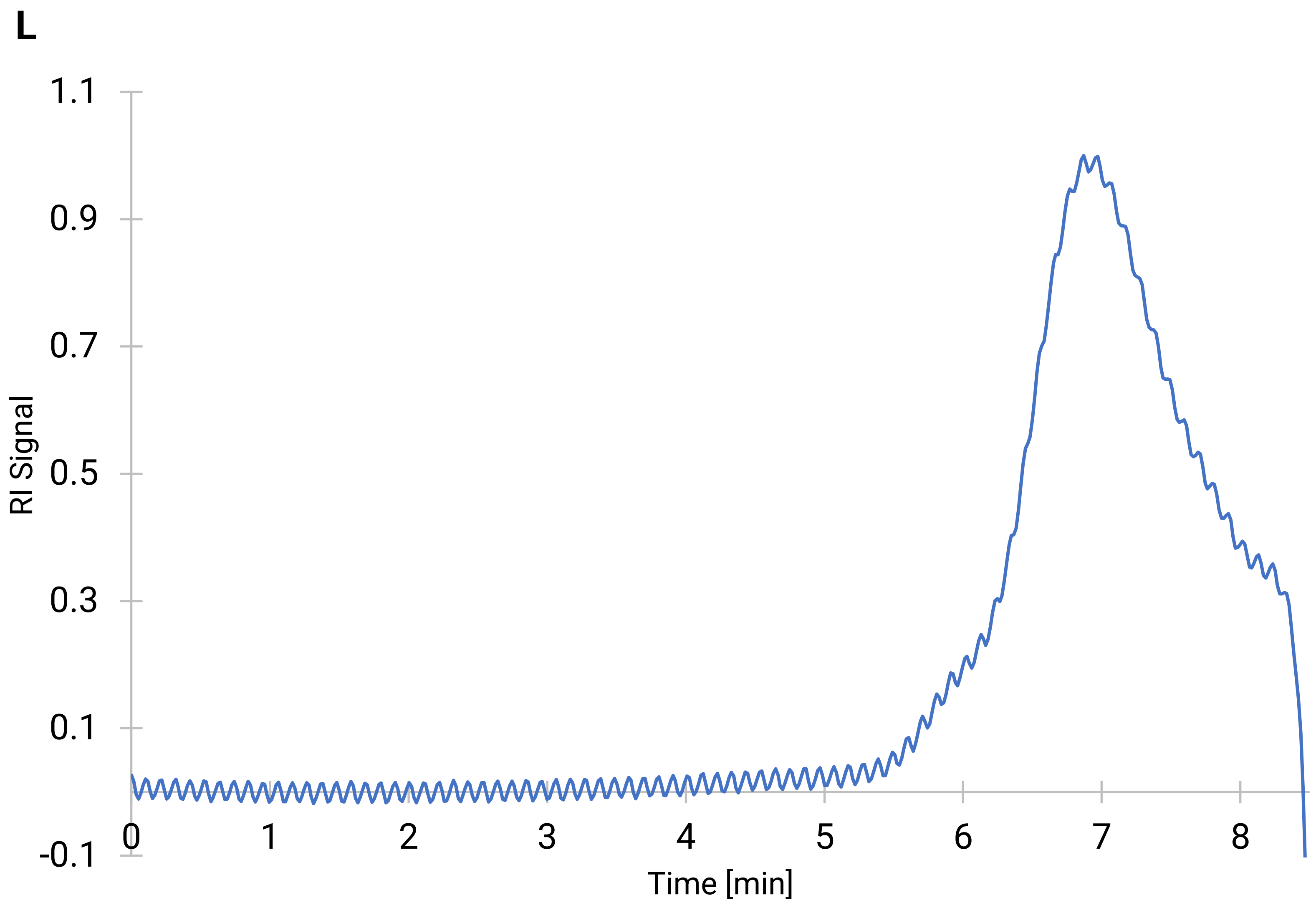


**Figure E4L: GPC plot of bio-polyester obtained from *C. necator* H16 Δ*phaC1* pCM66Tp$_{BAD}$:*hadA-phaC1437* when induced and co-fed with phloretic acid, harvested in late stage of cultivation, as per table S2.**

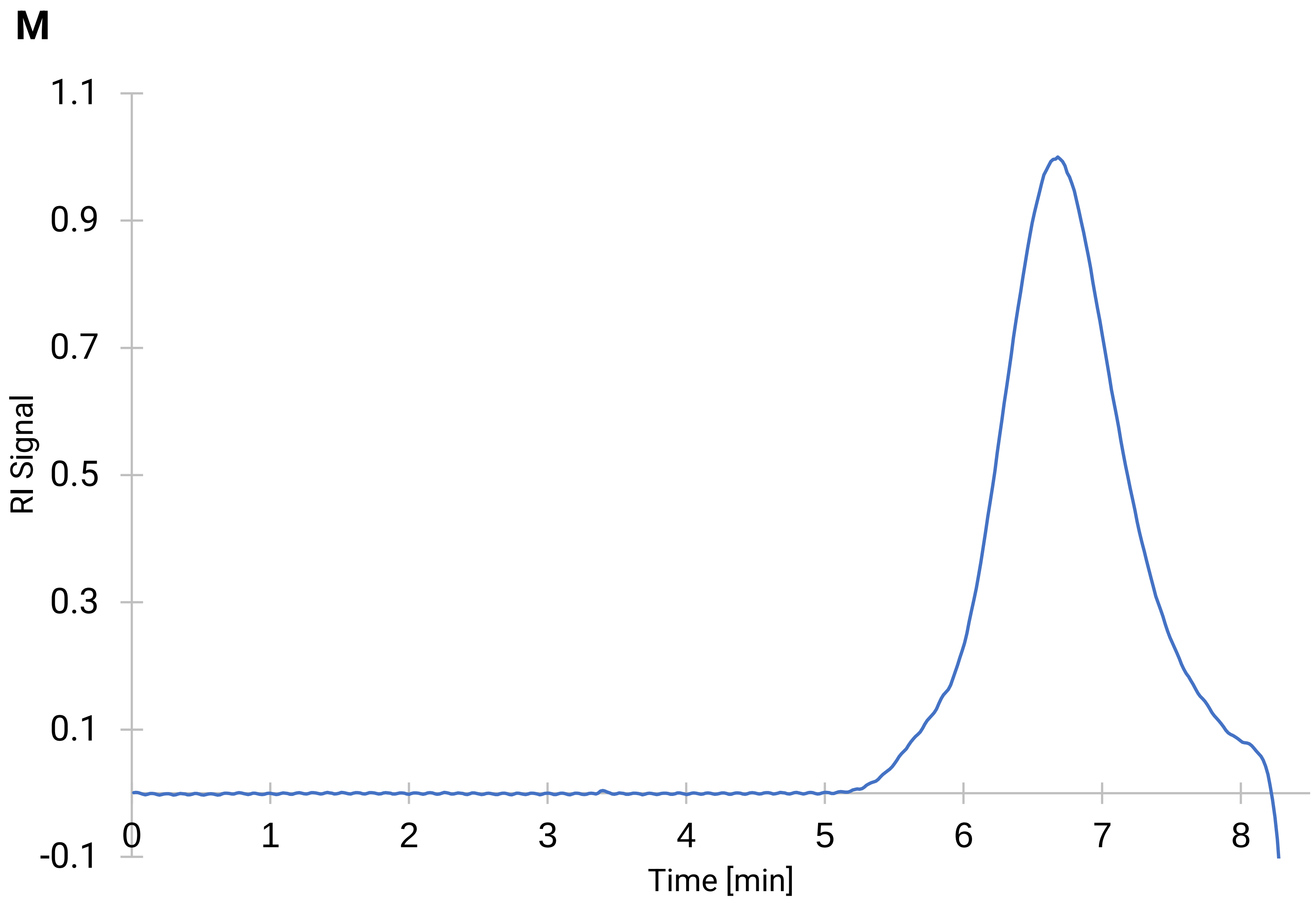


**Figure E4M: GPC plot of bio-polyester obtained from *C. necator* H16 Δ*phaC1* pCM66Tp$_{BAD}$:*hadA-phaC1437* when induced and co-fed with 5-hydroxymethyl-2-furancarboxylic acid, harvested early-on in cultivation, as per table S2.**

**N**

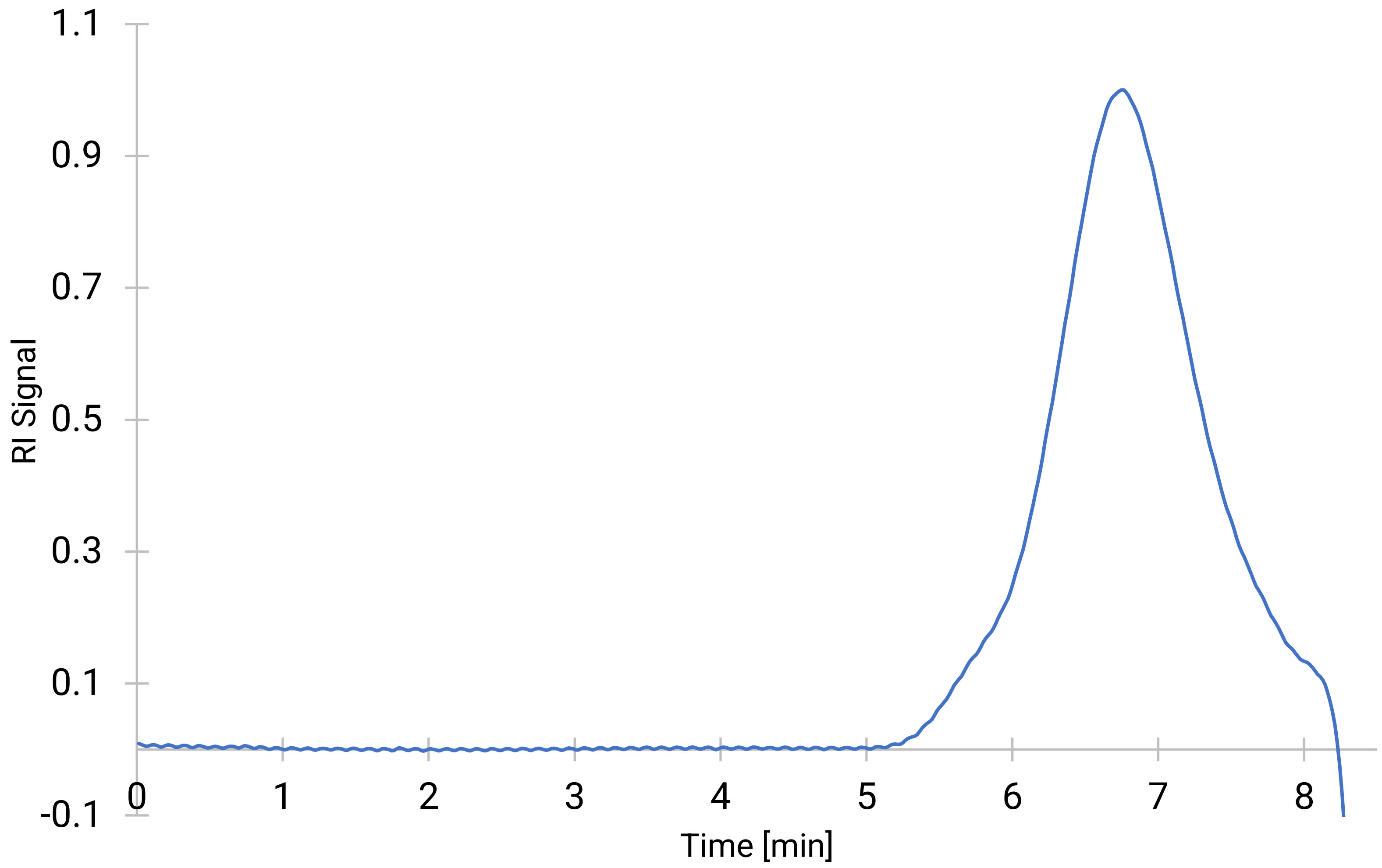


**Figure E4N: GPC plot of bio-polyester obtained from *C. necator* H16 Δ*phaC1* pCM66Tp$_{BAD}$:*hadA-phaC1437* when induced and co-fed with 5-hydroxymethyl-2-furancarboxylic acid, harvested in late stage of cultivation, as per table S2.**
