## Extended Data 4 for "Novel Aromatic Polyhydroxyalkanoates from Engineered *Cupriavidus necator* H16: Expanding Bio-Polyester Synthesis from Renewable Feedstocks"

DSC plots of bio-polyesters formed by wild-type and engineered *C. necator* H16

**A**

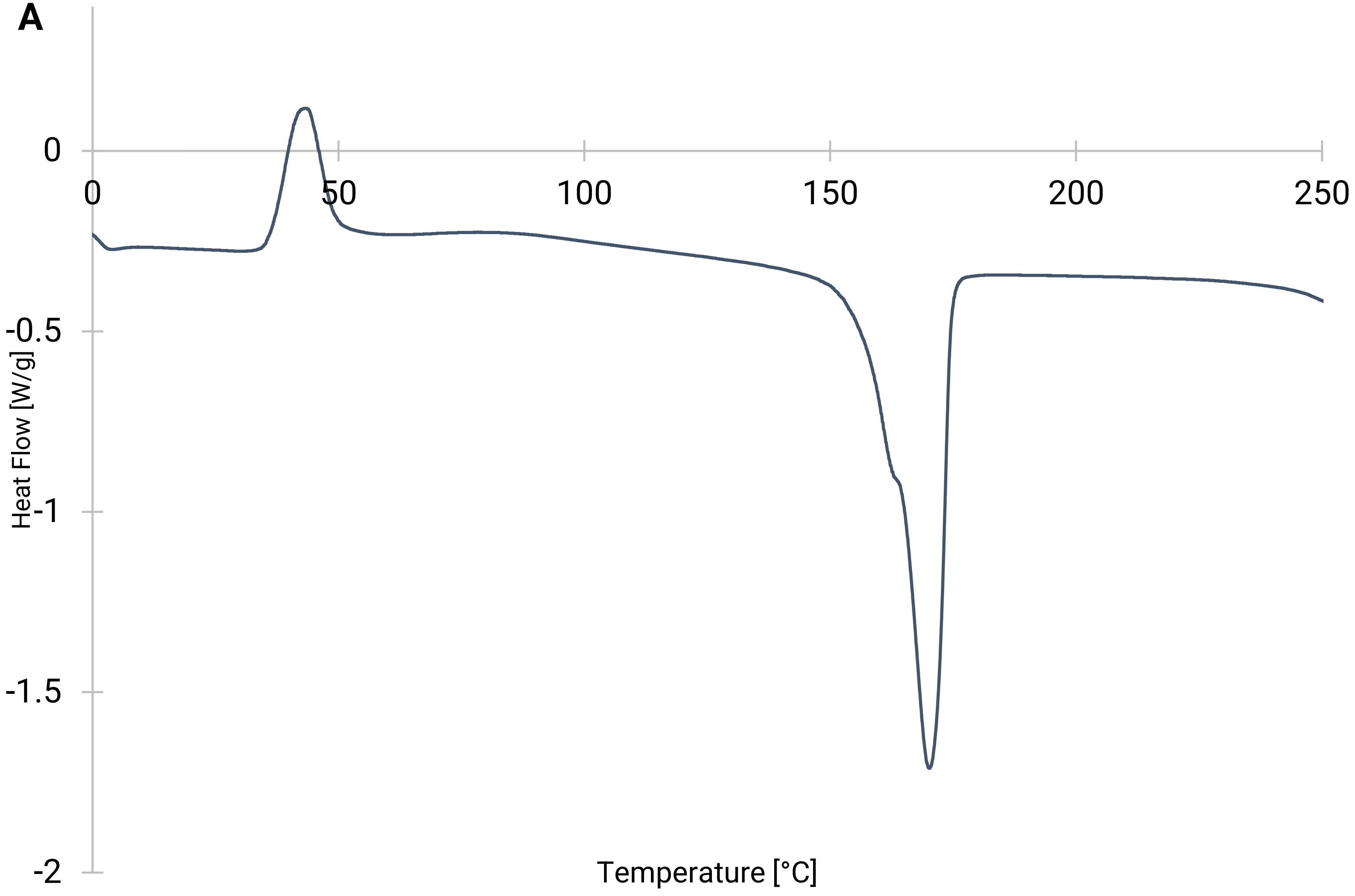


**Figure E5A: DSC plot of bio-polyester obtained from *C. necator* H16 (wild-type), as per table S2.**

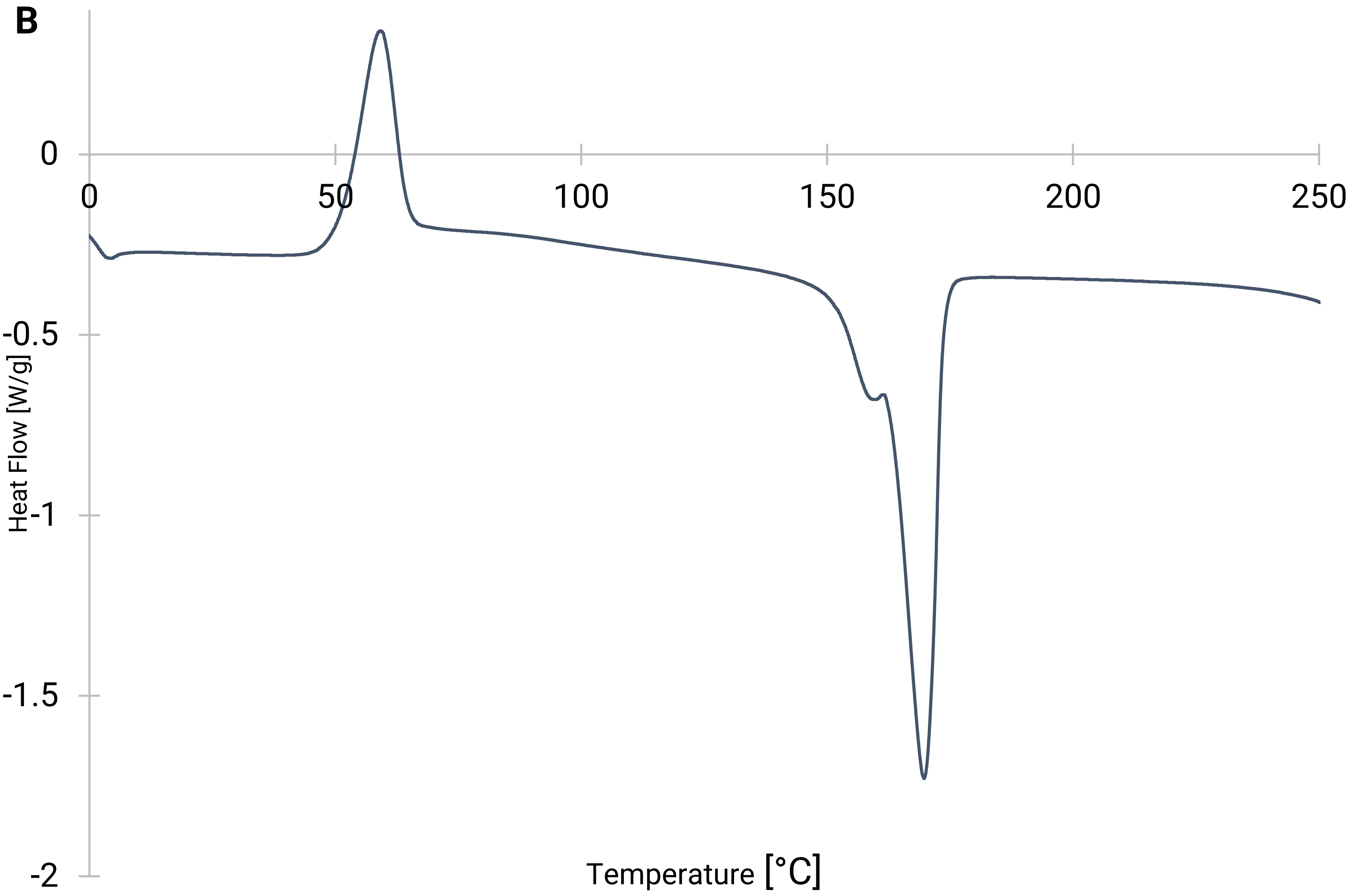


**Figure E5B: DSC plot of bio-polyester obtained from *C. necator* H16 Δ*phaC1* pCM66Tp$_{BAD}$:*pct540phaC1437* when not induced, as per table S2.**

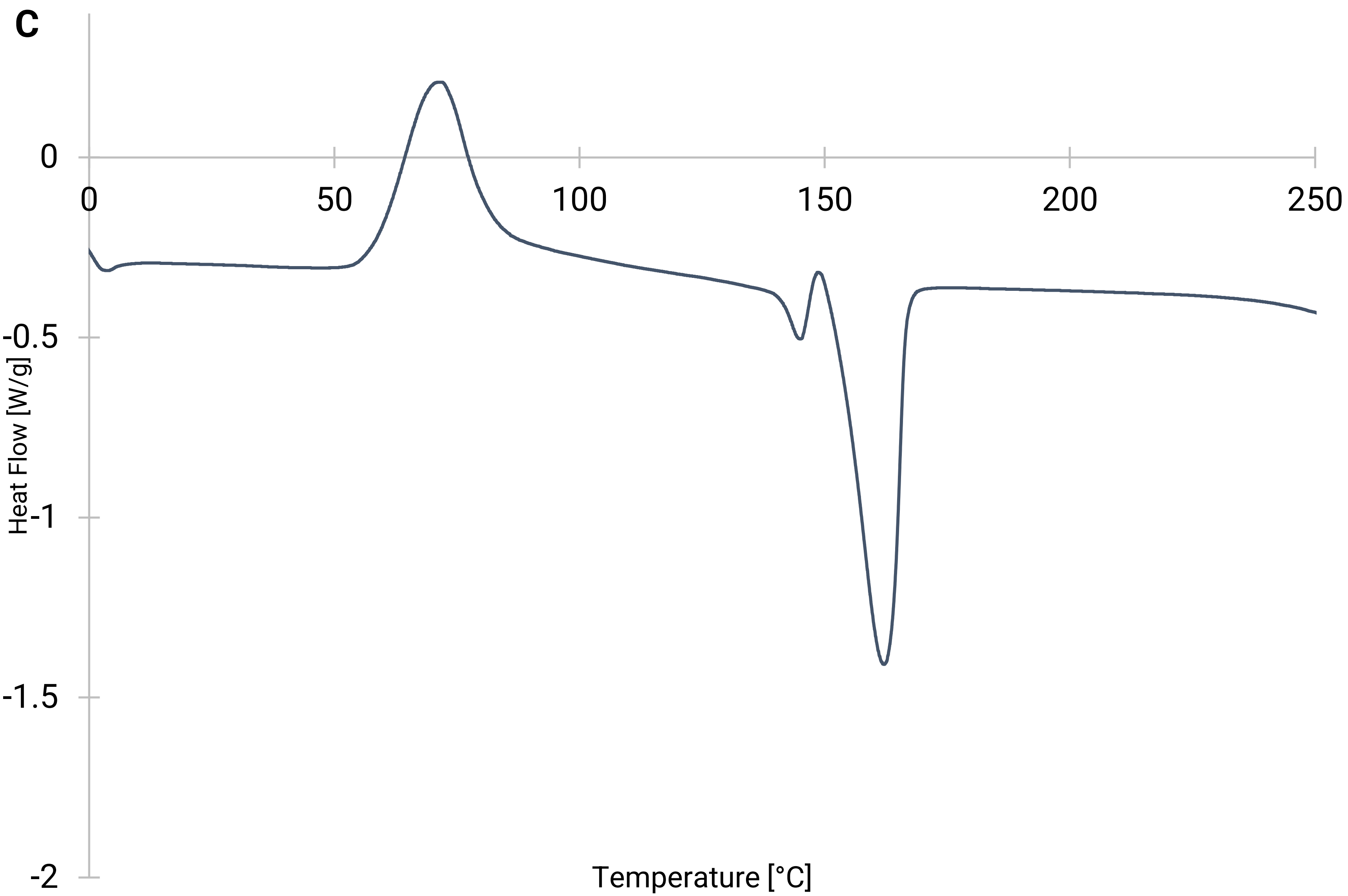


**Figure E5C: DSC plot of bio-polyester obtained from *C. necator* H16 Δ*phaC1* pCM66Tp$_{BAD}$:*pct540phaC1437* when induced, as per table S2.**

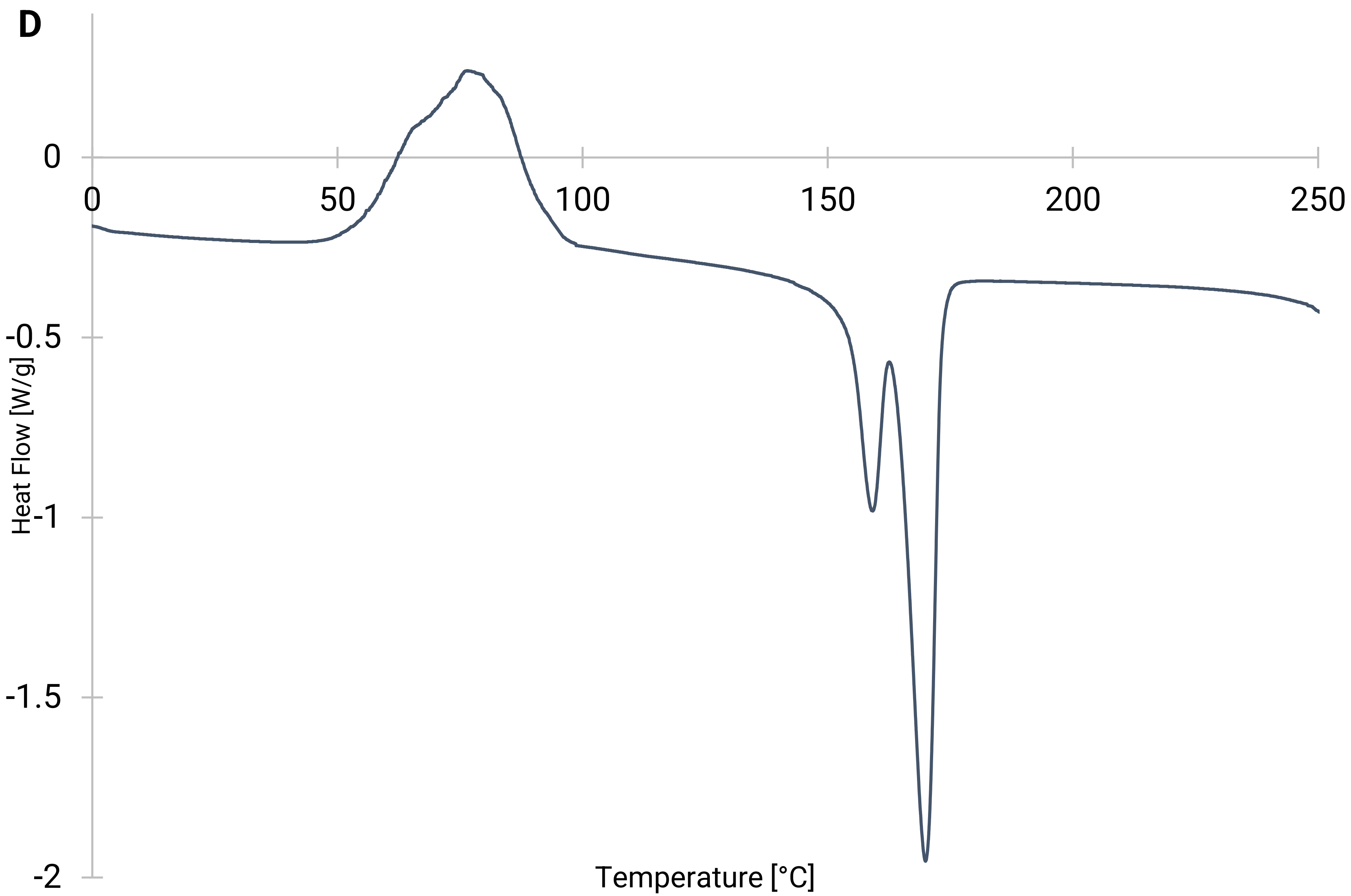


**Figure E5D: DSC plot of bio-polyester obtained from *C. necator* H16 Δ*phaC1* pCM66Tp$_{BAD}$:*hadAphaC1437* when not induced, as per table S2.**

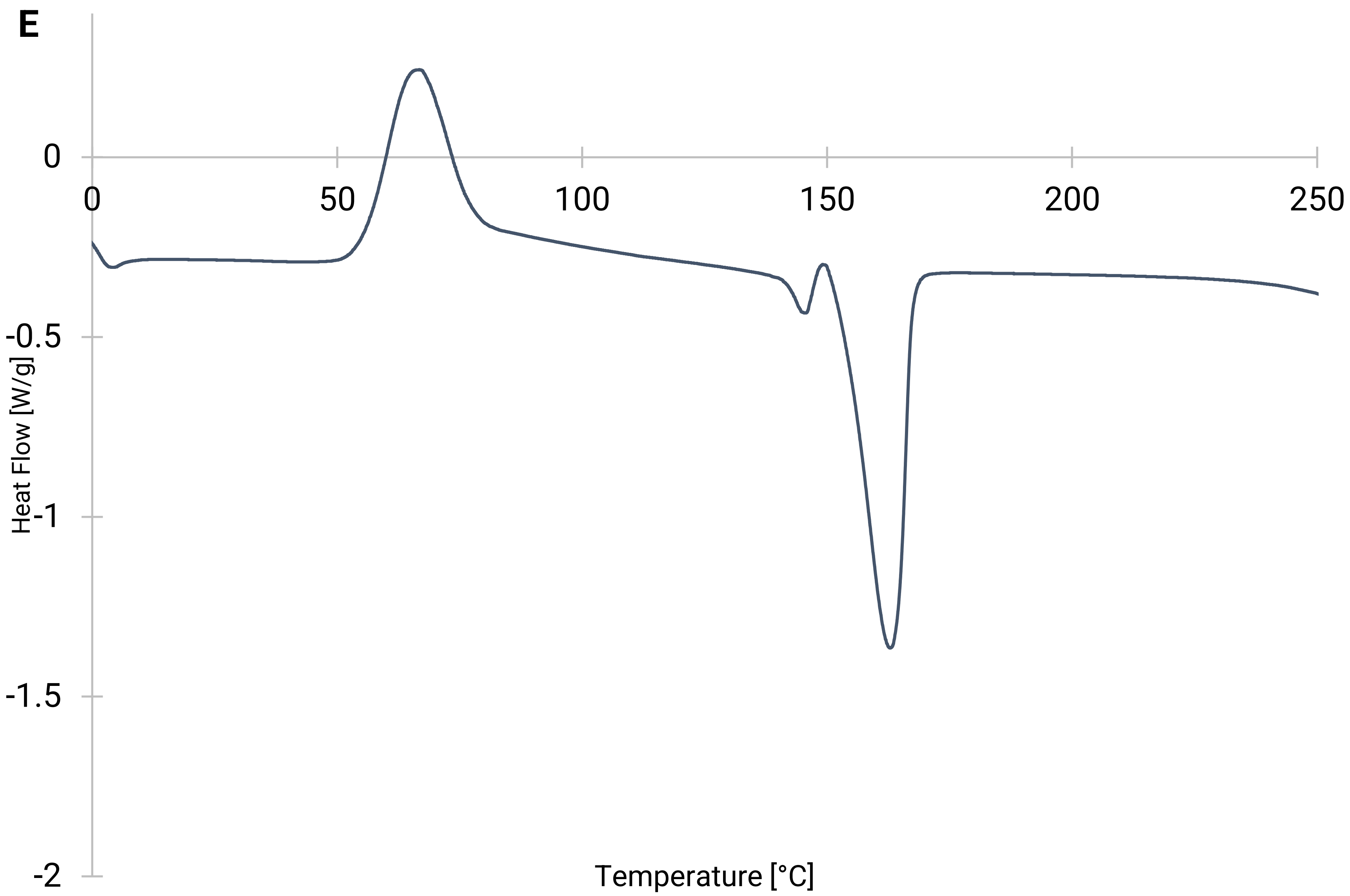


**Figure E5E: DSC plot of bio-polyester obtained from *C. necator* H16 Δ*phaC1* pCM66Tp$_{BAD}$:*hadAphaC1437* when induced, as per table S2.**

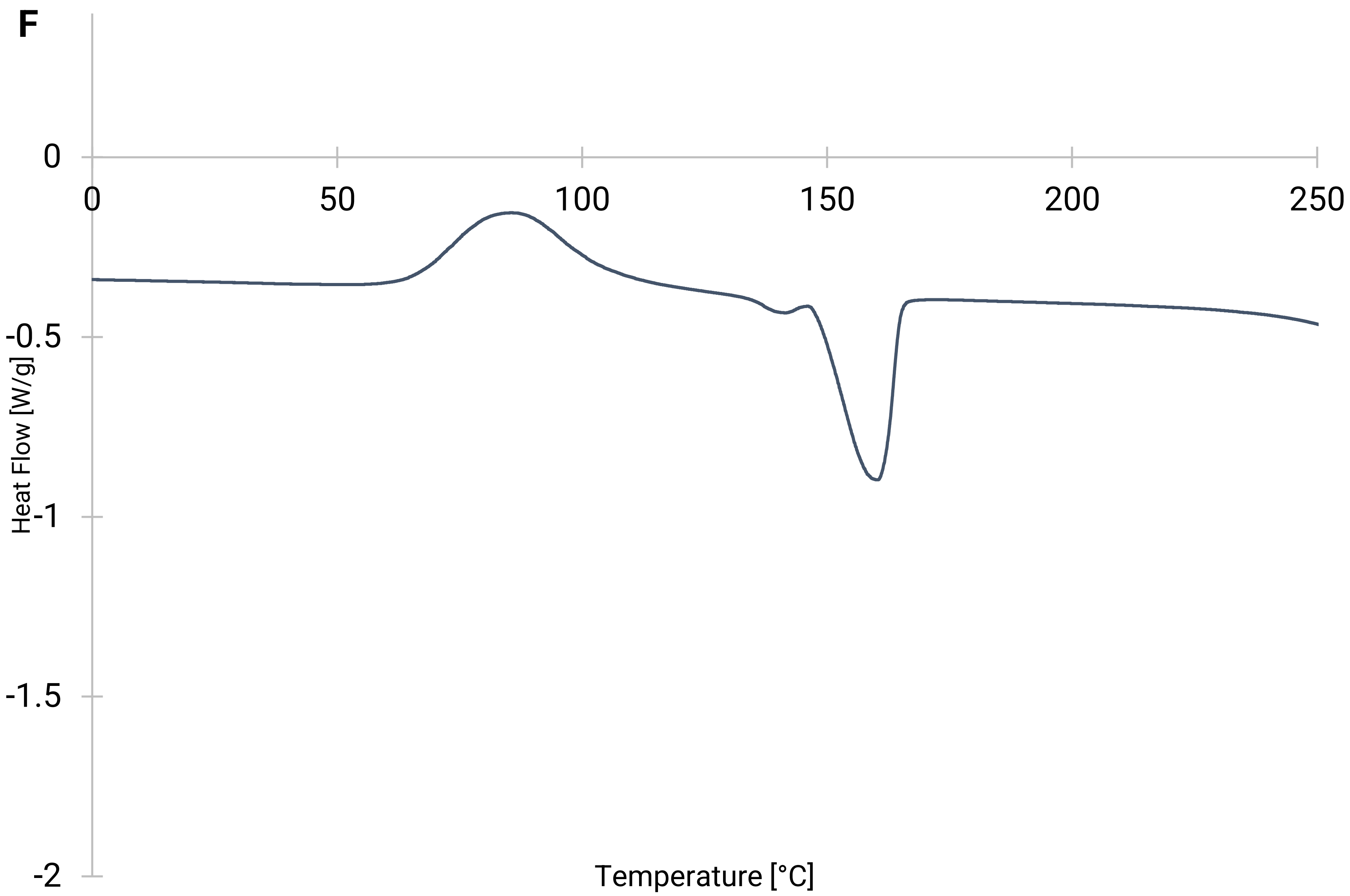


**Figure E5F: DSC plot of bio-polyester obtained from *C. necator* H16 Δ*phaC1* pCM66Tp$_{BAD}$:*pct540phaC1437* when induced and co-fed with 4-hydroxybutyric acid, as per table S2.**

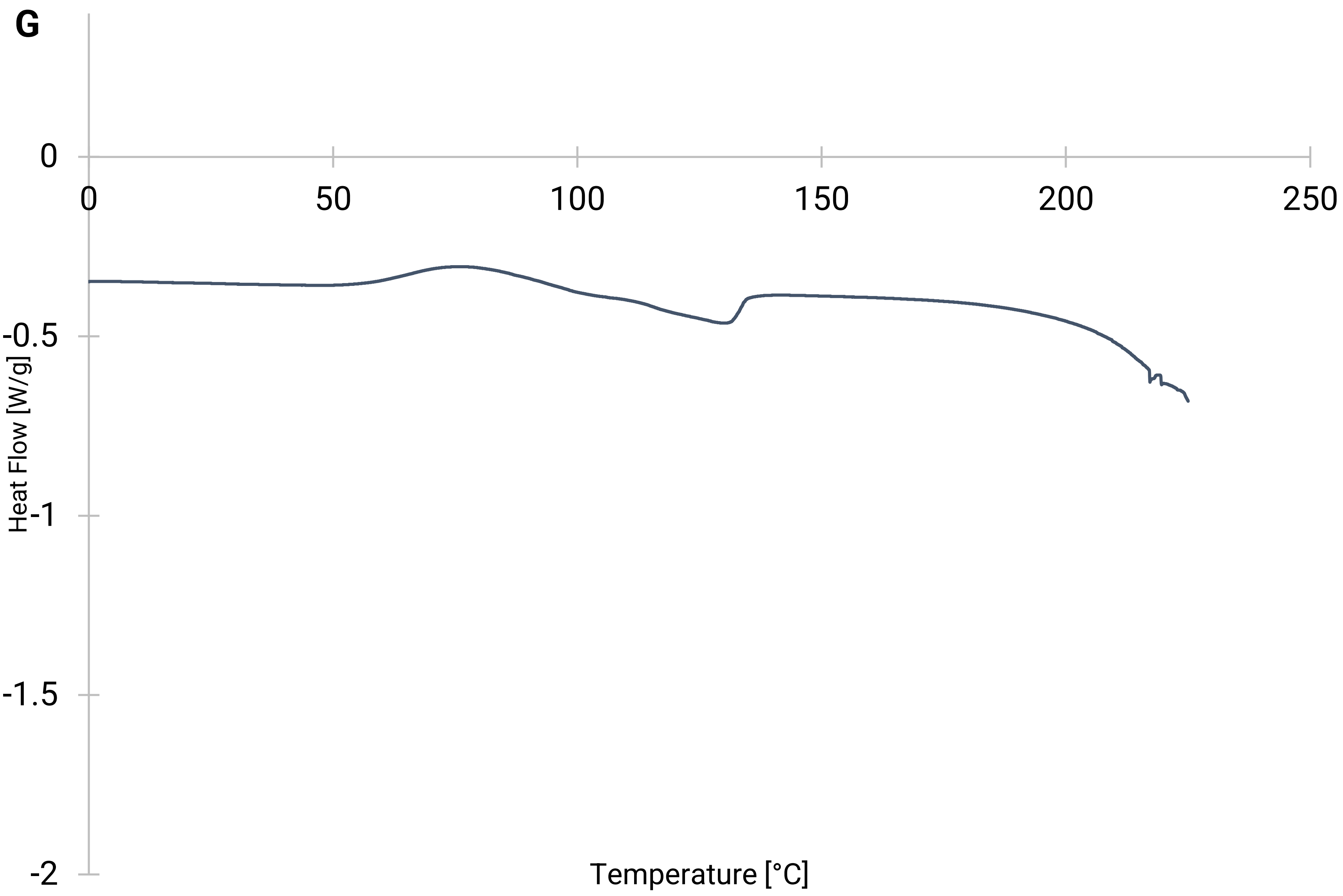


**Figure E5G: DSC plot of bio-polyester obtained from *C. necator* H16 Δ*phaC1* pCM66Tp$_{BAD}$:*hadAphaC1437* when induced and co-fed with 6-hydroxycaproic acid, as per table S2.**

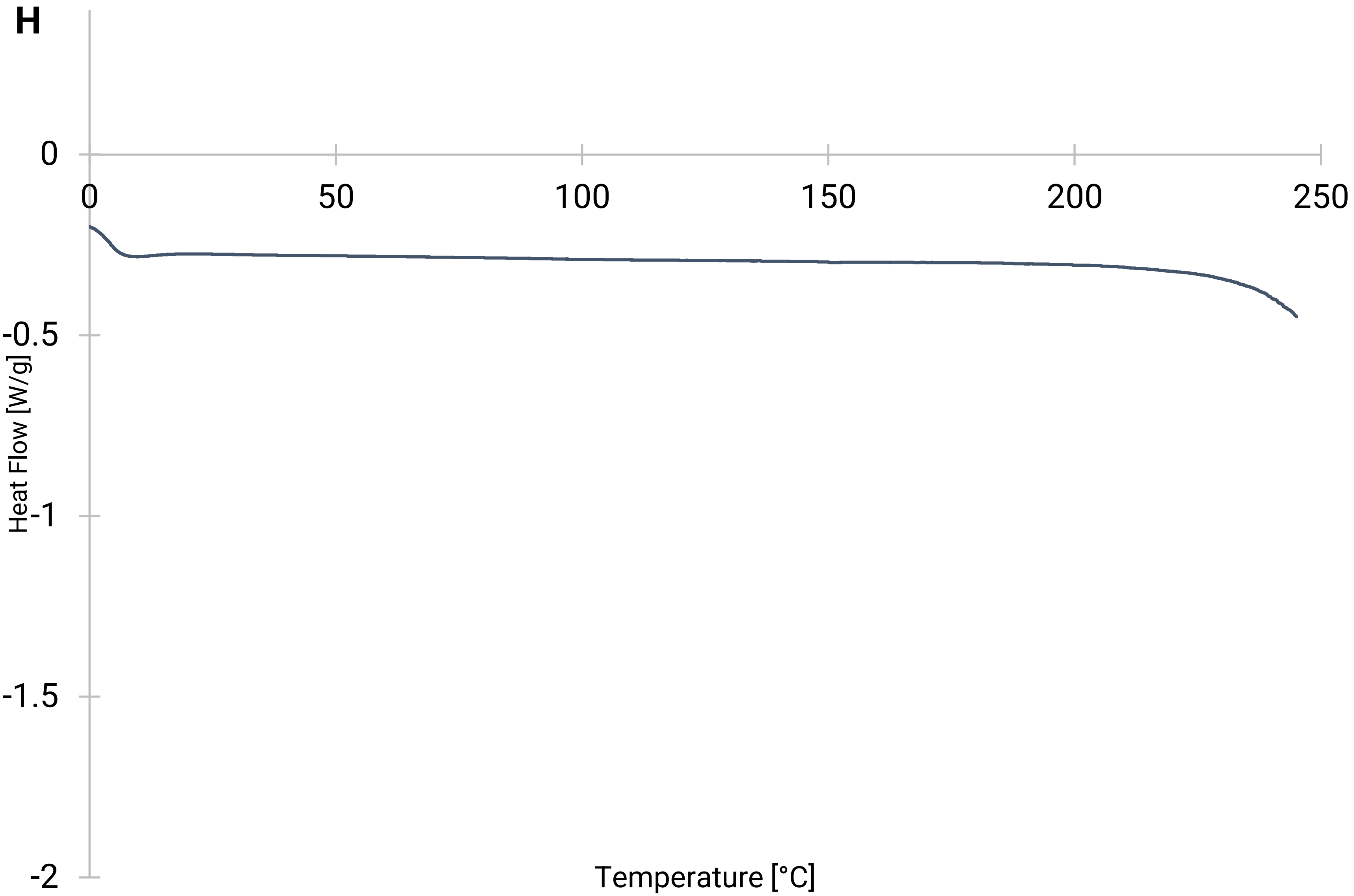


**Figure E5H: DSC plot of bio-polyester obtained from *C. necator* H16 Δ*phaC1* pCM66Tp$_{BAD}$:*hadAphaC1437* when induced and co-fed with phenyllactic acid, as per table S2.**

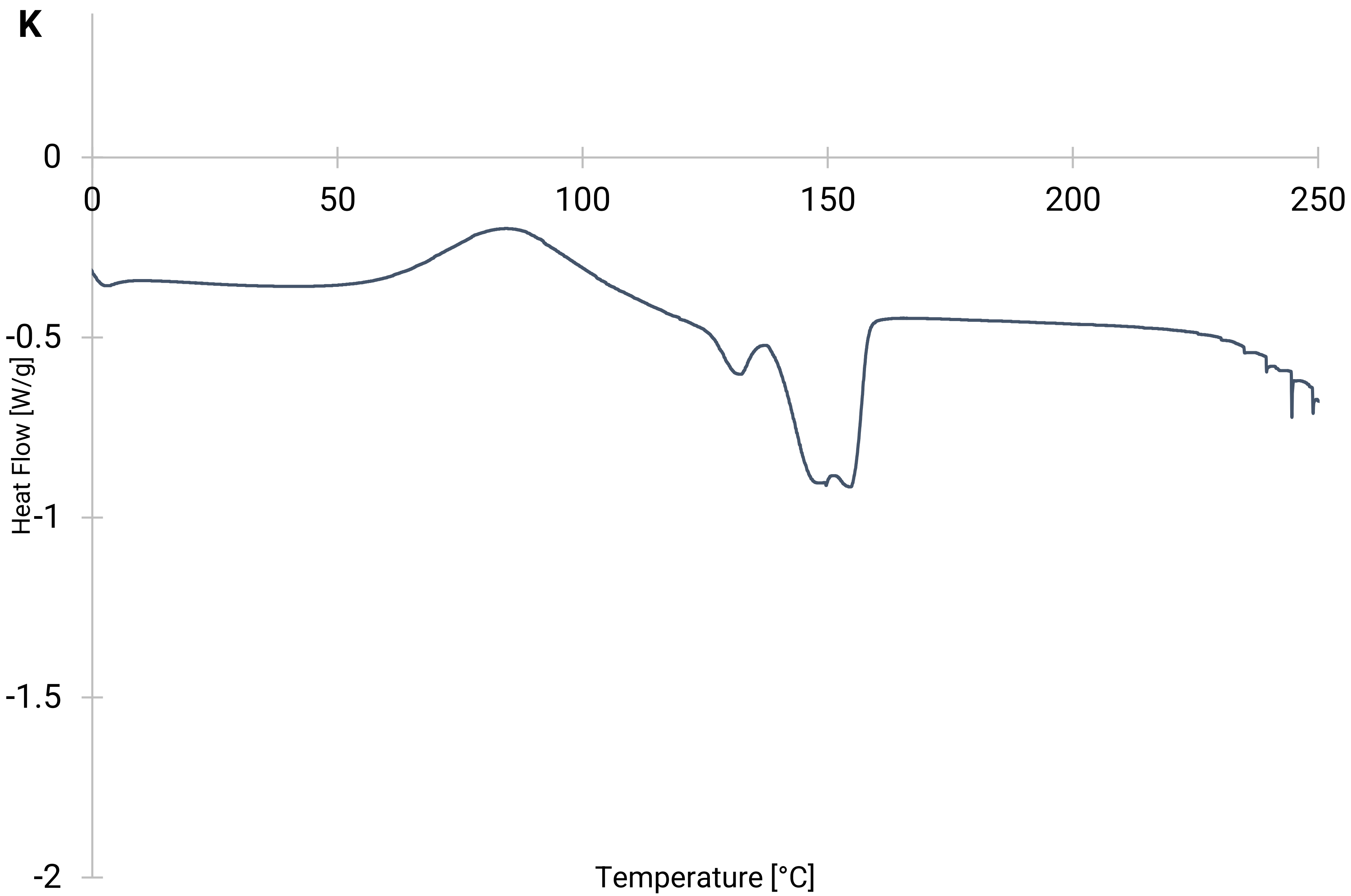


**Figure E5K: DSC plot of bio-polyester obtained from *C. necator* H16 Δ*phaC1* pCM66Tp$_{BAD}$:*hadAphaC1437* when induced and co-fed with phloretic acid, harvested early-on in cultivation, as per table S2.**

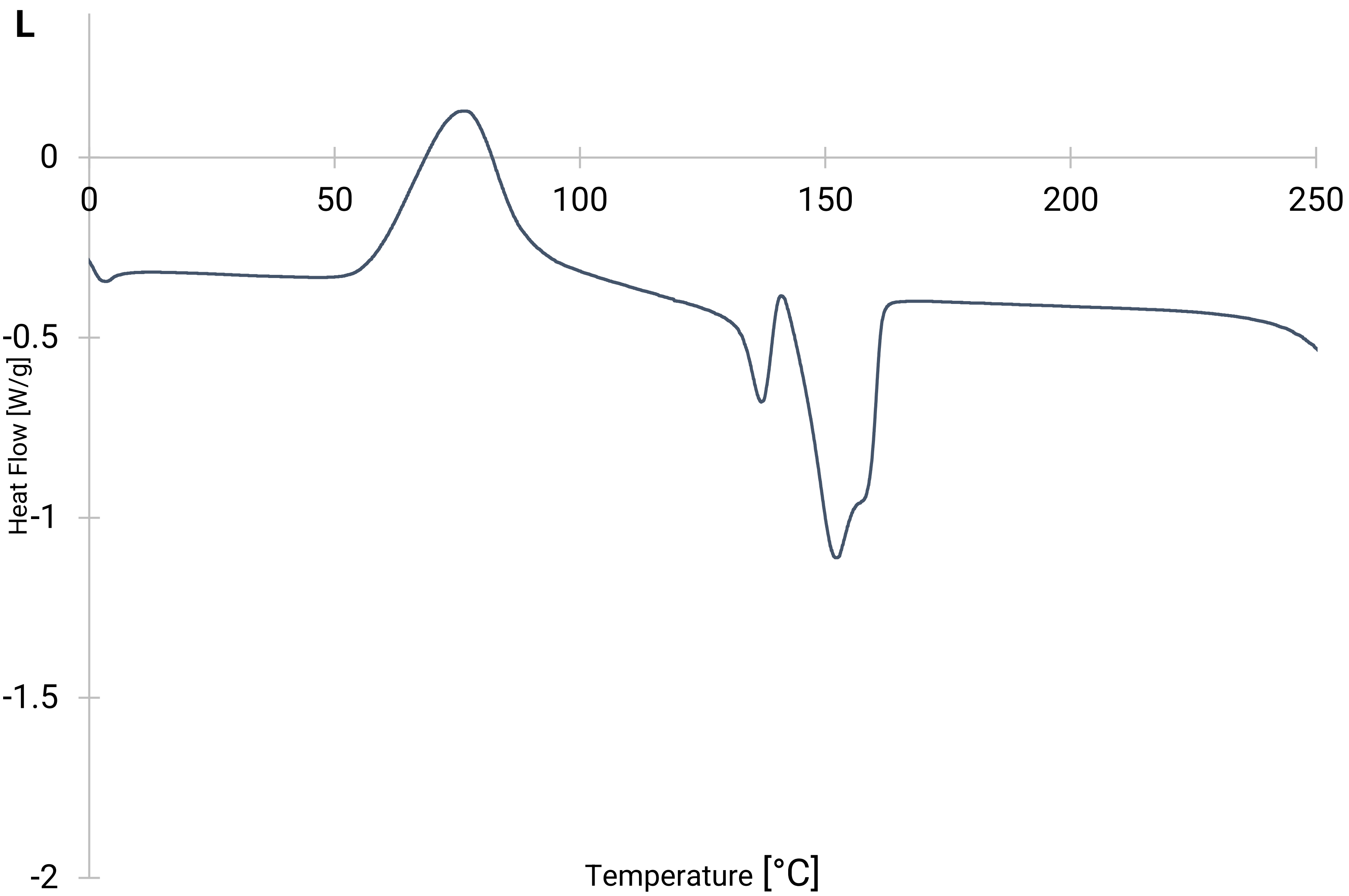


**Figure E5L: DSC plot of bio-polyester obtained from *C. necator* H16 Δ*phaC1* pCM66Tp$_{BAD}$:*hadAphaC1437* when induced and co-fed with phloretic acid, harvested in late stage of cultivation, as per table S2.**

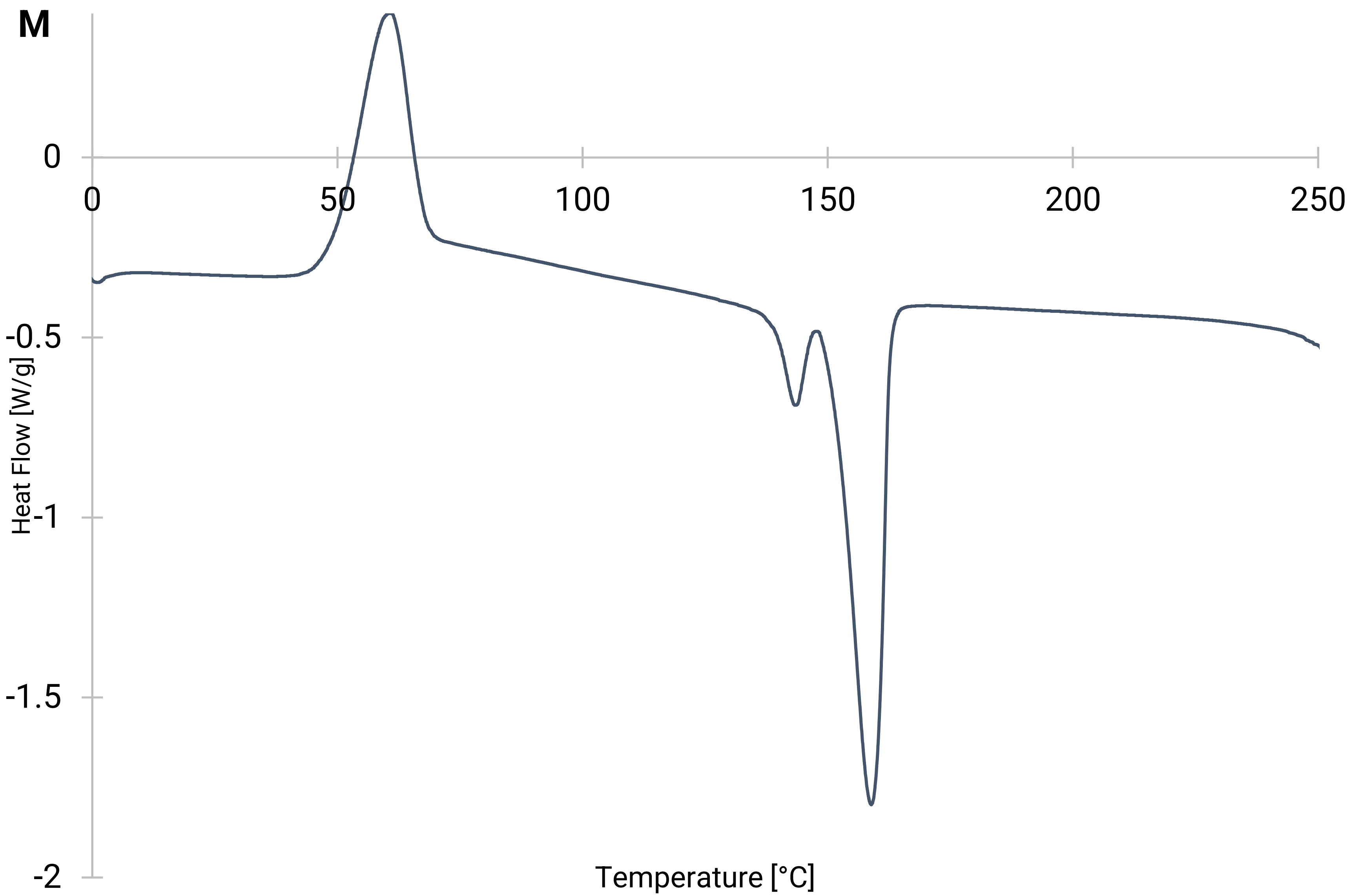


**Figure E5M: DSC plot of bio-polyester obtained from *C. necator* H16 Δ*phaC1* pCM66Tp$_{BAD}$:*hadAphaC1437* when induced and co-fed with 5-hydroxymethyl-2-furancarboxylic acid, harvested early-on in cultivation, as per table S2.**

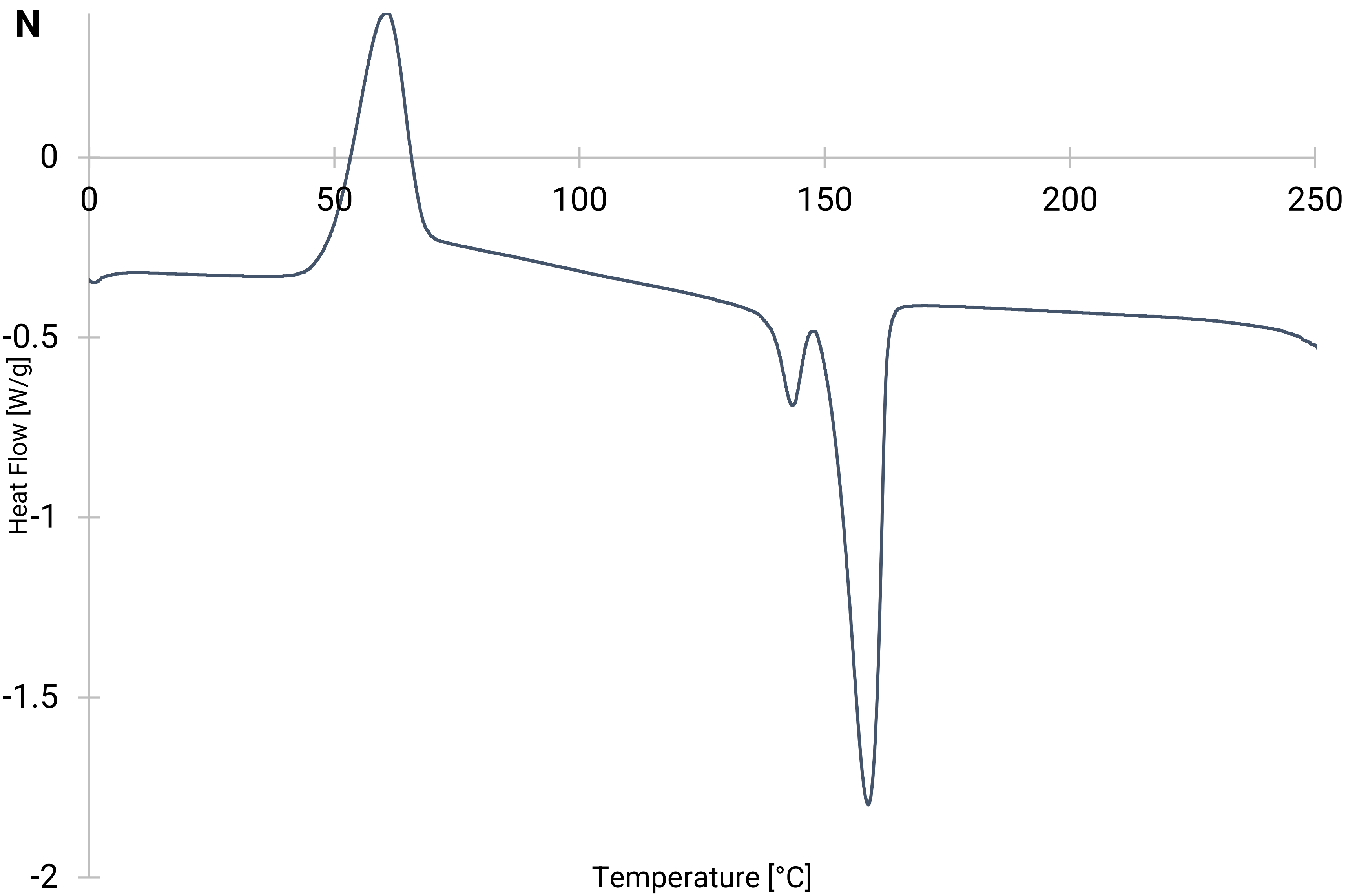


**Figure E5N: DSC plot of bio-polyester obtained from *C. necator* H16 Δ*phaC1* pCM66Tp$_{BAD}$:*hadAphaC1437* when induced and co-fed with 5-hydroxymethyl-2-furancarboxylic acid, harvested in late stage of cultivation, as per table S2.**
