## Extended Data 5 for "Novel Aromatic Polyhydroxyalkanoates from Engineered *Cupriavidus necator* H16: Expanding Bio-Polyester Synthesis from Renewable Feedstocks"

non-induced | induced

time post induction: 12 hours | 4 hours | 8 hours | 12 hours

strain: *marker* hadA pct540 | hadA pct540 | hadA pct540 | hadA pct540 *marker*

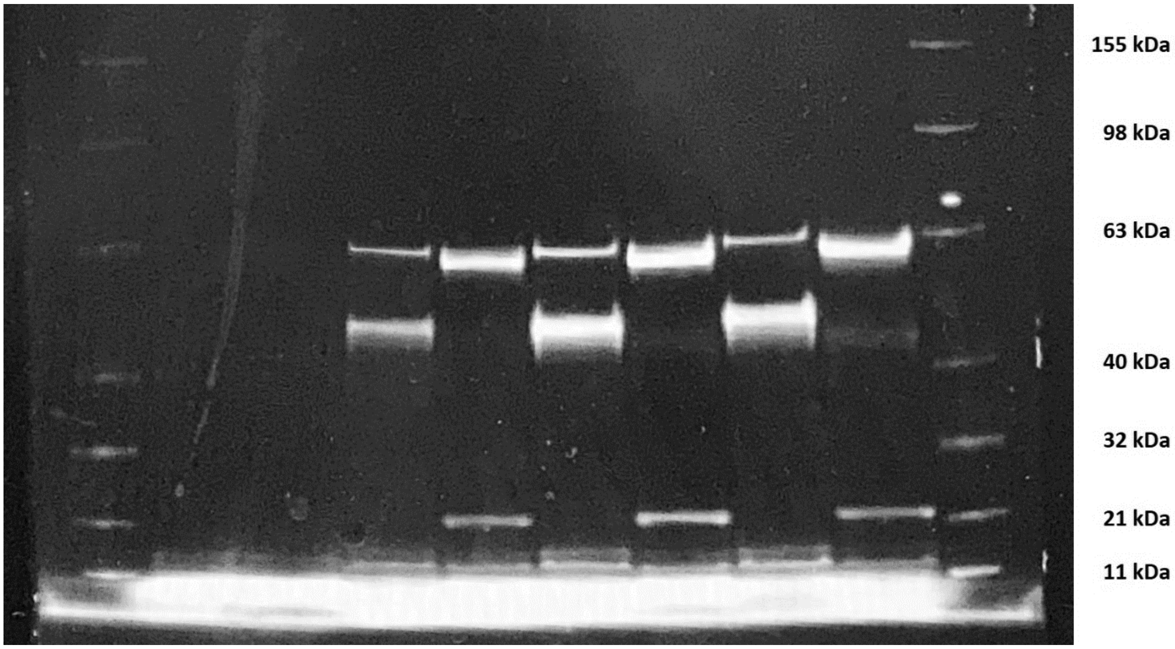


**Figure E5A: Expression of heterologous proteins by engineered *C. necator* H16 at different time-points after induction (corresponding to the cultivation profiles in figure 3). For each lane the same total protein concentration of crude-extract was loaded, except for lanes 1 & 10 which contain BenchMark™ Fluorescent Protein Standard (Thermo Fisher). Lane 2, 4, 6, 8 are from the hadA-strain; lane 3, 5, 7, 9 are from the pct540-strain. Inducer conc. = 1 g/L arabinose. Expected bands: *pct540* = 57 kDa, *hadA* = 44 kDa, *phaC1437* = 62 kDa.**

strain: pct540 | hadA

time post-induction: 6 hours | 9 hours | 4 hours | 9 hours

inducer conc. [g/L]: 0 0.1 1 | 0 0.1 1 | 0 0.1 1 | 0 0.1 1

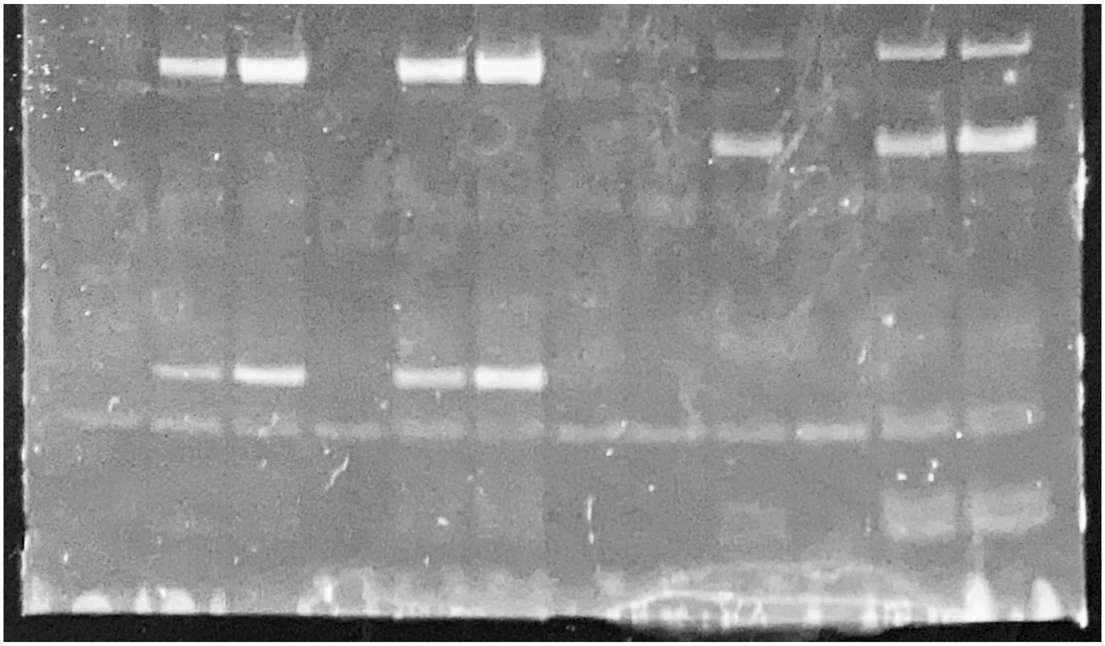

**Figure E5B: Expression of heterologous proteins by engineered *C. necator* H16 at different inducer (arabinose) concentrations and time-points after induction (corresponding to the cultivation profiles in figure 3). Lane 1 – 6: pct540-strain, lane 7 – 12: hadA-strain (same total protein concentration of crude-extract per lane).**
