## Supplementary Information 1 for "Novel Aromatic Polyhydroxyalkanoates from Engineered *Cupriavidus necator* H16: Expanding Bio-Polyester Synthesis from Renewable Feedstocks"

### 1 Growth and substrate uptake of engineered strains in dependence of heterologous enzyme expression

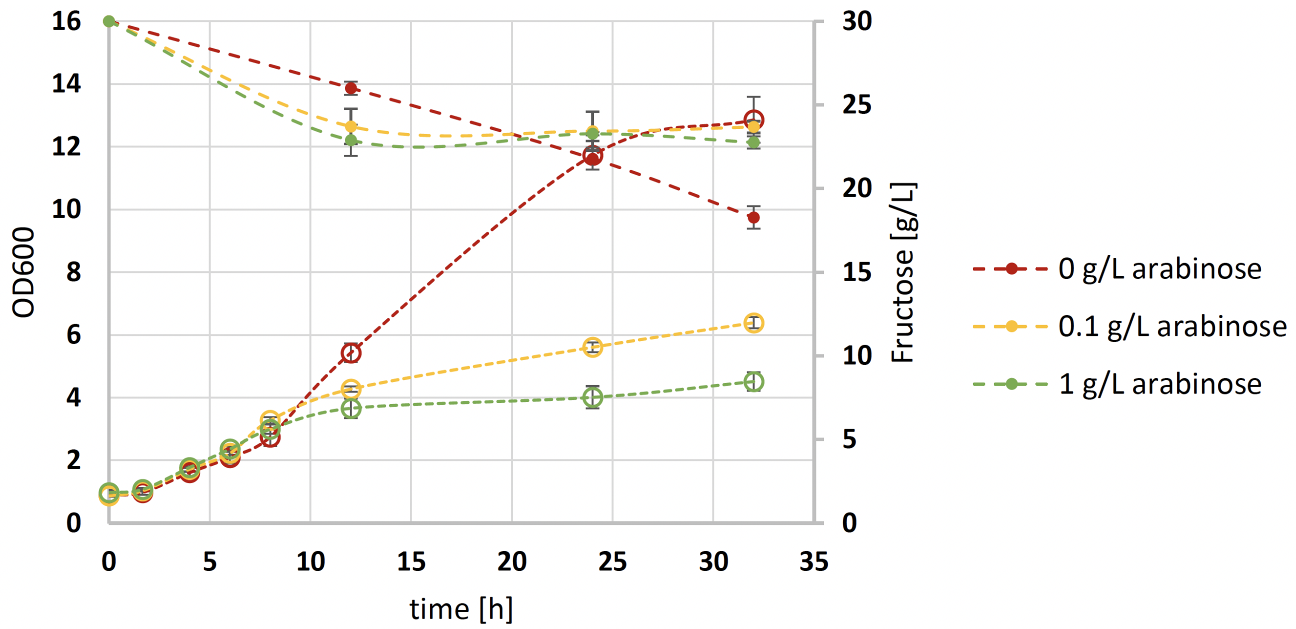


**Figure 1. Typical cultivation profile of *C. necator* H16 Δ*phaC1* pCM66Tp$_{BAD}$:*pct540-phaC1437* (*pct540*-strain) on minimal medium (MSM) with fructose as main carbon- and energy-source.** Growth is displayed by means of cell density (empty circles, absorption at 600 nm) and fructose consumption (full circles, concentration in supernatant per HPLC) over time at different levels of induction (arabinose; red = 0 g/L, yellow = 0.1 g/L, green = 1 g/L).

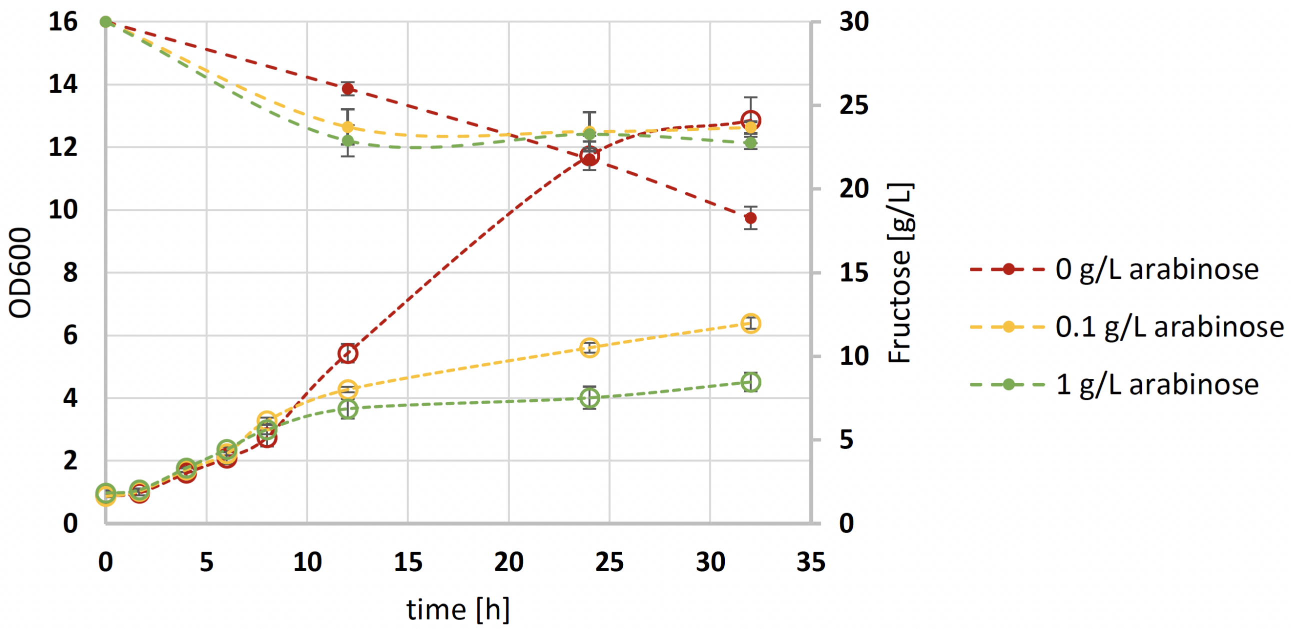


**Figure 2. Typical cultivation profile of *C. necator* H16 Δ*phaC1* pCM66Tp$_{BAD}$:*hadA-phaC1437* (*hadA*-strain) on minimal medium (MSM) with fructose as main carbon- and energy-source.** Growth is displayed by means of cell density (empty circles, absorption at 600 nm) and fructose consumption (full circles, concentration in supernatant per HPLC) over time at different levels of induction (arabinose; red = 0 g/L, yellow = 0.1 g/L, green = 1 g/L).

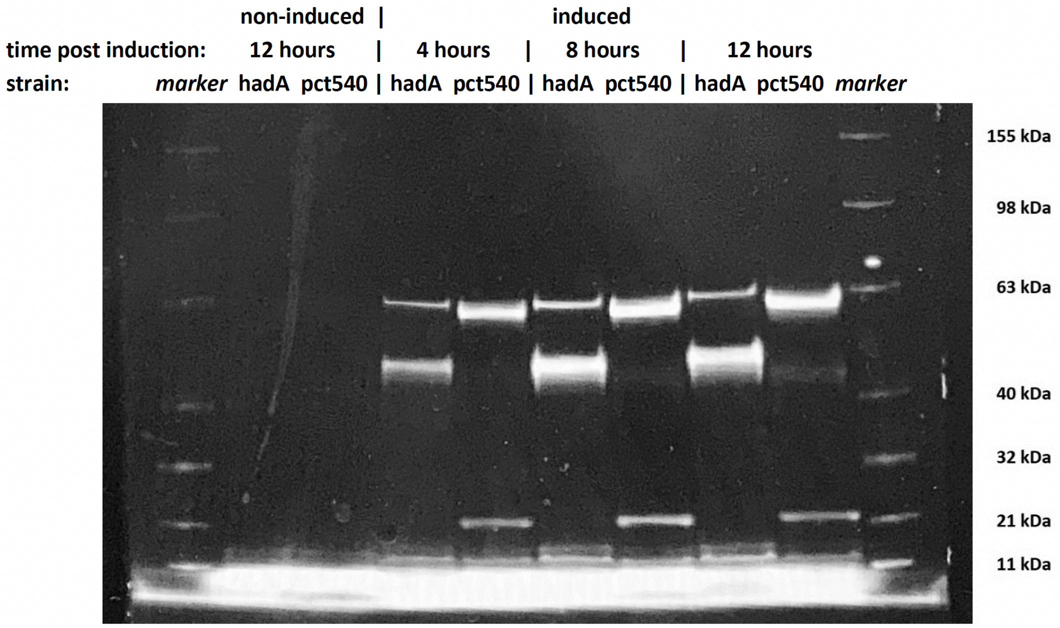


**Figure 3.** **Expression of heterologous proteins by engineered *C. necator* H16 at different time points after induction (corresponding to the cultivation profiles in Figures S1 & S2).** For each lane the same total protein concentration of crude-extract was loaded, except for lanes 1 & 10 which contain BenchMark® Fluorescent Protein Standard (Thermo Fisher). Lane 2, 4, 6, and 8 are from the hadA-strain; lane 3, 5, 7, and 9 are from the *pct540*-strain. Inducer conc. = 1 g/L arabinose. Expected bands: Pct540 = 57 kDa, HadA = 44 kDa, PhaC1437 = 62 kDa.

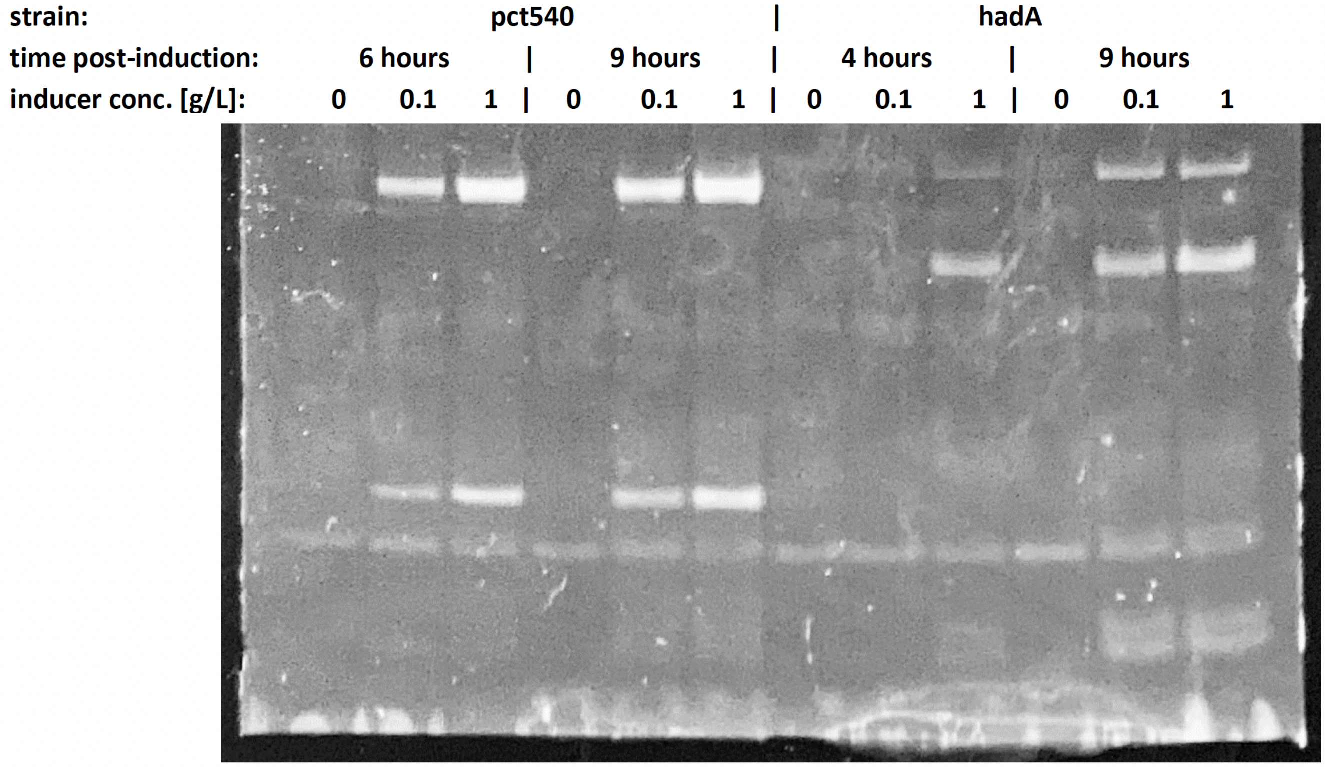


**Figure 4.** **Expression of heterologous proteins by engineered *C. necator* H16 at different inducer (arabinose) concentrations and time-points after induction (corresponding to the cultivation profiles in Figures S1 & S2).** Lane 1 to 6: *pct540*-strain; lane 7 to 12: *hadA*-strain (same total protein concentration of crude-extract per lane).

### 2 Extended discussion on correlation of growth and enzyme expression

Transgenic strains of *C. necator* H16 were cultivated on minimal medium (MSM) to the middle of the exponential growth phase, then washed and transferred to fresh medium, including inducer as applicable, such that the time-points of induction and inoculation coincided ($t_0$ = 0 h). Hence, the growth profiles in Figures S1 and S2 directly correspond to Figures S3 and S4 and the respective sampling time points used for the protein work. Detection of the heterologous enzymes was performed directly on the crude protein extract, taking advantage of fluorescence-based staining of the Lumio®-tags at the C-terminus of the respective proteins.

Without induction of the arabinose promoter ($P_{BAD}$), no expression of the proteins from the heterologous genes under the control of $P_{BAD}$ was evident, albeit catalytic PHA synthase activity existed (PHA was also formed independently from induction). Protein expression levels at inducer (arabinose) concentrations of 0.1 and 1 g/L were only differential in the early stage of the cultivations, equalizing after about 6 h. Expression of the isocaprenoyl-CoA:2-hydroxyisocaproate CoA-transferase "*hadA*" (44 kDa) was seemingly even stronger than expression of the propionate CoA-transferase "*pct540*" (57 kDa); hence, poor expression of the CoA transferase was likely not an issue, which in reverse supports the conclusion that the PHA synthase is highly active. Relative to the strain carrying the isocaprenoyl-CoA:2-hydroxyisocaproate CoA-transferase, expression of the PHA synthase "*phaC1437*" (62 kDa) appeared much higher in the strain carrying the propionate CoA-transferase. Expression of the propionate CoA-transferase seemed to be problematic: a truncated protein is present at $\sim$22 kDa, the complete protein is hard to identify on the protein gels – the respective band potentially overlaps with that of the PHA synthase (a faint band just below the PHA synthase in Figure S4 may be indicative of the propionate CoA-transferase). This could also explain the seemingly higher expression of the PHA synthase in the strain carrying the propionate CoA-transferase in Figure S3.

### 3 Identification of co-polymer compositions through $^1$H- and $^{13}$C-NMR spectroscopy

For all $^1$H-NMR spectra (see Extended Data 2) the double quadruplet resonances at 2.55 and 2.48 ppm and the sextet resonance at 5.27 ppm corresponded to the two protons on the C2 and the single proton on the C3 of the 3-hydroxybutyric acid repeat-unit, respectively. The incorporation of 3-hydroxypropionic acid repeat-unit into the PHA, indicated by the triplet resonance at 4.33 ppm of the C3-methylene, was in good accordance with literature[1]. The incorporation of 4-hydroxybutyric and 6-hydroxyhexanoic acid was identified by shifts of 4.11 ppm and 4.07 ppm, representative of the protons on the respective $\omega$-carbons of the originating hydroxyacyl, respectively.

The aryl-aliphatic, as well as aromatic PHAs, were identified by distinct chemical shifts commonly associated with protons on the aromatic ring. For phenyllactic acid, the broad peak at 7.21 ppm falls into the area expected for aromatic protons, also the broad peak overlapping the sextet resonance of 3-hydroxybutyric acid at 5.27 ppm is consistent with $^1$H-NMR spectra of aryl-aliphatic co polyesters in the literature[2]. Further, the expected number of aromatic and aliphatic carbons for phenyllactic acid could be deducted from the $^{13}$C-NMR spectrum of the respective PHA. By extension, polymers obtained from the other two aryl-alkyl hydroxy carbonates, mandelic and 2-hydroxy-4-phenylbutyric acid, show analogous split patterns at 7.28 and 7.21 ppm respectively, and narrow methine peaks at 5.12 and 5.13 ppm respectively.

Comparing $^1$H-NMR spectra of phloretic and of 5-hydroxy-2-methylfuranoic acid monomers, the two doublet resonances at 7.12 and 6.98 ppm and at 7.07 and 6.5 ppm in the co-polymers formed from these compounds were indicative of the respective repeat units. Areas below the peaks, weighted by the number of protons they represent, were indicative of the fractions of the respective repeat units.

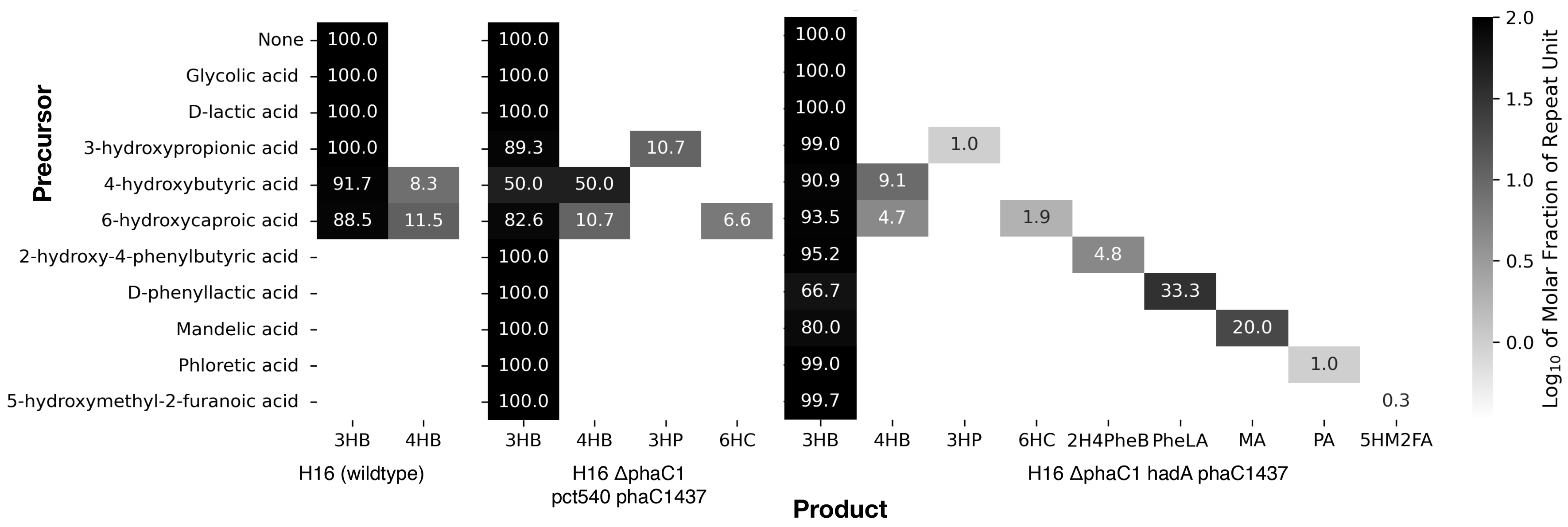


**Figure 5.** Spectrum of co-polyesters obtained from engineered strains of *C. necator* H16 (wildtype and engineered strains) provided with exogenous precursors. Fractions are molar [%] of the respective repeat unit derived from ratios as determined by $^{1}$H-NMR spectroscopy (see Extended Data 2).
Abbreviations: 3HB = 3-hydroxybutyrate, 4HB = 4-hydroxybutyrate, 3HP = 3-hyroxypropionate, 6HC = 6-hydroxycaproate, 2H4PheB = 2-hydroxy-4-phenylbutyrate, PheLA = phenyllactate, MA = mandelate, PA = phloretate, 5HM2F = 5-hydroxymethyl-2-furanoate.

| Strain | Induced | Polymer Composition | $M_n$ [g/mol] | Dev.($M_n$) [g/mol] | Mw [g/mol] | Dev.($M_w$) [g/mol] | PDI | # |
|---|---|---|---|---|---|---|---|---|
| H16 (wildtype) | n.a. | P3HB[a] | 397400 | 8639 | 680800 | 7278 | 1.71 | A |
| H16 Δ*phaC1 pct540-phaC1437* | no | P3HB | 69190 | 2649 | 134300 | 1542 | 1.94 | B |
| H16 Δ*phaC1 pct540-phaC1437* | yes | P(3HB-co-xHA)[b] | 45300 | 2057 | 123400 | 1199 | 2.72 | C |
| H16 Δ*phaC1 hadA-phaC1437* | no | P3HB | 54820 | 2965 | 130400 | 1234 | 2.38 | D |
| H16 Δ*phaC1 hadA-phaC1437* | yes | P(3HB-co-xHA)[c] | 10900 | 377 | 27690 | 404 | 2.54 | E |
| H16 Δ*phaC1 pct540-phaC1437* | yes | P(3HB-co-4HB) | 25220 | 897 | 74120 | 873 | 2.94 | F |
| H16 Δ*phaC1 hadA-phaC1437* | yes | P(3HB-co-4HB-co-6HC) | 11810 | 725 | 25120 | 518 | 2.13 | G |
| H16 Δ*phaC1 hadA-phaC1437* | yes | P(3HB-co-PheLA) | 13170 | 329 | 21750 | 330 | 1.65 | H |
| H16 Δ*phaC1 hadA-phaC1437* | no | P3HB* | 62940 | 1492 | 137700 | 1527 | 2.19 | I |
| H16 Δ*phaC1 hadA-phaC1437* | no | P3HB† | 50010 | 1209 | 98250 | 1093 | 1.96 | J |
| H16 Δ*phaC1 hadA-phaC1437* | yes | P(3HB-co-PA)* | 61270 | 1468 | 180700 | 1964 | 2.95 | K |
| H16 Δ*phaC1 hadA-phaC1437* | yes | P(3HB-co-PA)† | 14450 | 689 | 33700 | 649 | 2.33 | L |
| H16 Δ*phaC1 hadA-phaC1437* | yes | P(3HB-co-5HM2F)* | 25620 | 4445 | 56160 | 8637 | 2.19 | M |
| H16 Δ*phaC1 hadA-phaC1437* | yes | P(3HB-co-5HM2F)† | 13670 | 2224 | 44820 | 1973 | 3.28 | N |

**Table 1. Molecular weights and polydispersity indexes of natural PHB, non-natural PHB, and select aliphatic & aromatic PHAs, as determined by GPC.** Abbreviations: Dev. = deviation, n.a. = not applicable, P = *poly*-, 3HB = 3-hydroxybutyrate, 4HB = 4-hydroxybutyrate, 6HC = 6-hydroxycaproate, PheLA = phenyllactate, PA = phloretate, 5HM2F = 5-hydroxymethyl 2-furanoate, xHA = indeterminate repeat unit, $M_n$ = number average molecular weight, $M_w$ = weight average molecular weight, PDI = polydispersity index ($M_w/M_n$), # sample identifier (corresponds to GPC traces in Extended Data 3); [a] natural PHB, [b,c] PHB with minor fraction of indeterminate repeat unit of natural origin, * harvested in early stage of cultivation, † harvested in late stage of cultivation.

| Strain | Volume [mL] | $OD_{600}$ | CDW [mg] | Biomass [mg/L] | Factor |
|---|---|---|---|---|---|
| H16 (wildtype) | 50 | 24.5 | 0.3377 | 6.754 | 0.28 |
| | 50 | 22.7 | 0.912 | 18.24 | 0.8 |
| | 50 | 21.9 | 0.2837 | 5.674 | 0.26 |
| | 50 | 16 | 0.2068 | 4.136 | 0.26 |
| | 50 | 13.4 | 0.174 | 3.48 | 0.26 |
| | | strain average | | | 0.37±0.24 |
| H16 Δ*phaC1* *pct540-phaC1437* | 50 | 4.88 | 0.0696 | 1.392 | 0.29 |
| | 50 | 5.08 | 0.093 | 1.86 | 0.37 |
| | 50 | 5.16 | 0.081 | 1.62 | 0.31 |
| | 50 | 6.32 | 0.0936 | 1.872 | 0.3 |
| | 50 | 6.32 | 0.076 | 1.52 | 0.24 |
| | | strain average | | | 0.30±0.05 |
| H16 Δ*phaC1* *hadA-phaC1437* | 50 | 18.1 | 0.2145 | 4.29 | 0.24 |
| | 50 | 22.7 | 0.235 | 4.7 | 0.21 |
| | 50 | 17.7 | 0.203 | 4.06 | 0.23 |
| | 50 | 19.7 | 0.3055 | 6.11 | 0.31 |
| | 50 | 10.7 | 0.1345 | 2.69 | 0.25 |
| | 100 | 3.76 | 0.0876 | 0.876 | 0.23 |
| | 100 | 6.32 | 0.179 | 1.79 | 0.28 |
| | 100 | 4.44 | 0.137 | 1.37 | 0.31 |
| | 100 | 4.08 | 0.109 | 1.09 | 0.27 |
| | 100 | 3.92 | 0.112 | 1.12 | 0.29 |
| | | strain average | | | 0.26±0.04 |
| | | overall average | | | 0.31±0.05 |

**Table 2.** Determination of correlation factor for optical density (OD per absorption at 600 nm) to cell dry-weight (CDW) for *Cupriavidus necator* H16 wildtype and the respective mutant strains when cultured in minimal medium.

| Strain | Precursor | CDW [g] | PHA [g] | PHA/Biomass [g/g] |
|---|---|---|---|---|
| H16 (wildtype) | none | 0.291 | 0.059 | 0.203 |
| | glycolic acid | 0.338 | 0.266 | 0.786 |
| | lactic acid | 0.912 | 0.224 | 0.246 |
| | 3-hydroxypropanoic acid | 0.284 | 0.207 | 0.730 |
| | 4-hydroxybutanoic acid | 0.207 | 0.142 | 0.687 |
| | 6-hydroxyhexanoic acid | 0.174 | 0.128 | 0.736 |
| | strain average | | | 0.56±0.27 |
| H16 Δ*phaC1 pct540-phaC1437* | none | 0.054 | 0.027 | 0.494 |
| | glycolic acid | 0.070 | 0.005 | 0.072 |
| | lactic acid | 0.093 | 0.030 | 0.323 |
| | 3-hydroxypropanoic acid | 0.081 | 0.024 | 0.296 |
| | | 0.081 | 0.058 | 0.713 |
| | 4-hydroxybutanoic acid | 0.094 | 0.035 | 0.374 |
| | | 0.090 | 0.057 | 0.631 |
| | 6-hydroxyhexanoic acid | 0.076 | 0.024 | 0.316 |
| | | 0.087 | 0.066 | 0.752 |
| | strain average | | | 0.44±0.22 |
| H16 Δ*phaC1 hadA-phaC1437* | none | 0.207 | 0.128 | 0.618 |
| | none | 0.886 | 0.855 | 0.965 |
| | glycolic acid | 0.215 | 0.146 | 0.681 |
| | lactic acid | 0.235 | 0.163 | 0.694 |
| | 3-hydroxypropanoic acid | 0.203 | 0.138 | 0.680 |
| | | 0.198 | 0.081 | 0.409 |
| | 4-hydroxybutanoic acid | 0.306 | 0.120 | 0.393 |
| | | 0.147 | 0.057 | 0.388 |
| | 6-hydroxyhexanoic acid | 0.135 | 0.077 | 0.572 |
| | | 0.170 | 0.068 | 0.401 |
| | 2-hydroxy-4-phenylbutanoic acid | 0.063 | 0.030 | 0.476 |
| | | 0.089 | 0.074 | 0.834 |
| | phenyllactic acid | 0.088 | 0.037 | 0.422 |
| | | 0.180 | 0.090 | 0.501 |
| | mandelic acid | 0.179 | 0.113 | 0.631 |
| | | 0.133 | 0.093 | 0.697 |
| | phloretic acid | 0.137 | 0.064 | 0.467 |
| | | 0.097 | 0.075 | 0.779 |
| | strain average | | | 0.58±0.17 |

**Table 3.** Comparison of PHA content per cell dry-weight (CDW) for *Cupriavidus necator* H16 wildtype and the respective mutant strains cultured in minimal medium with or without supplemented precursor.

### 4 Operation of Bio-Electrochemical System for autotrophic cultivation of *Cupriavidus necator* H16

Cultivated in serum-bottles on a mixture of $H_2/O_2/CO_2$ with an over-pressure of 22 psi, growth of *C. necator* H16 was limited to a maximum $OD_{600}$ of 3.5 (Figure S6). Further, since H16 only forms polyhydroxyalkanoates under strict N-limitation when metabolism is autotrophic[3], the concentration of ammonium in the medium was reduced by 4-fold for cultures on gas (based on the average maximum biomass obtained in shake-flask experiments when fructose was the substrate).

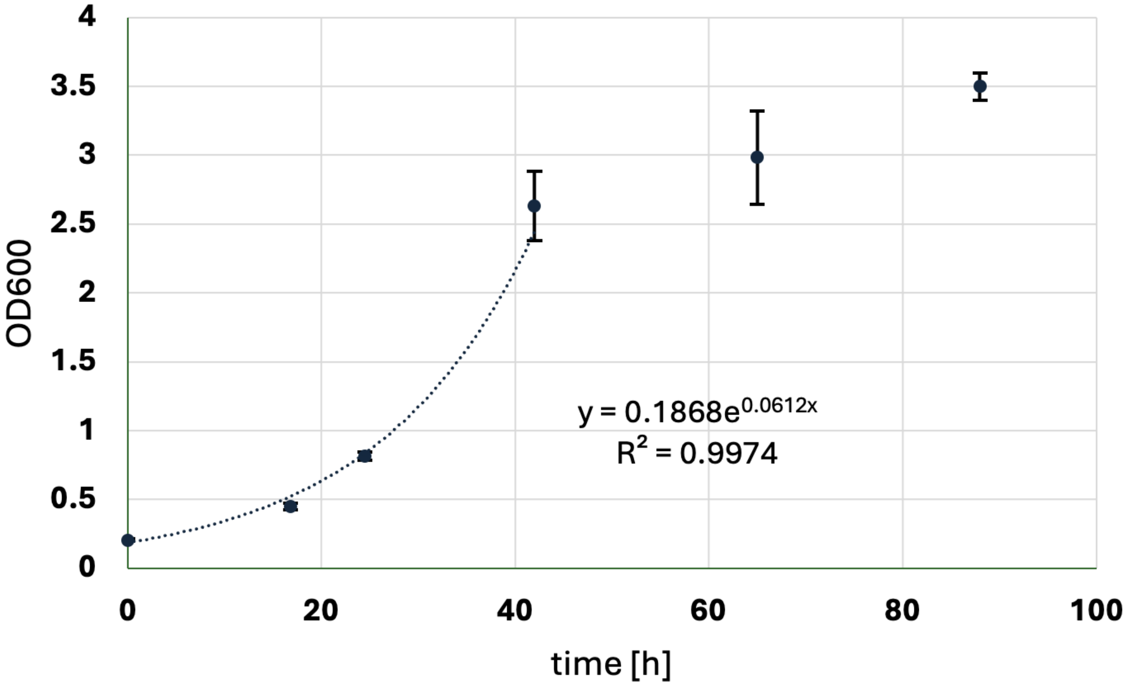


**Figure 6.** Autotrophic growth of *Cupriavidus necator* H16 Δ*phaC1* pCM66Tp$_{BAD}$:*hadA-phaC1437* in serum bottles (averaged data based on triplicates, $H_2$:$O_2$:$CO_2$ = 64:16:20 [v/v/v], 22 psi).

Neither the growth rate nor the maximum biomass of the autotrophic cultures were affected by the reduction of the nitrogen source. Further, while biosynthesis of PHB became active in the late growth stage of these cultures, polymer yield could not be quantified with confidence, due to the low volume (of the total 125 mL the culture occupied only 20% to maintain sufficient headspace for the substrate) and biomass density in the serum-bottles. Table S4 comprises key data associated with autotrophic cultivation trials of *C. necator* H16 on gas in serum-bottles.

| Substrate (ratio at 22 psi) | Ammonium | $\mu$ [1/h] | max. $OD_{600}$ | PHB |
|---|---|---|---|---|
| $H_2/O_2/CO_2$ (64:16:20) [v/v/v] | 20 mM | 0.06±0.01 | 3.6 | no |
| $H_2/O_2/CO_2$(64:16:20) [v/v/v] | 5 mM | 0.06±0.01 | 3.5 | yes |

**Table 4.** Key-Data associated with autotrophic cultivation of *Cupriavidus necator* H16 in serum-bottles.

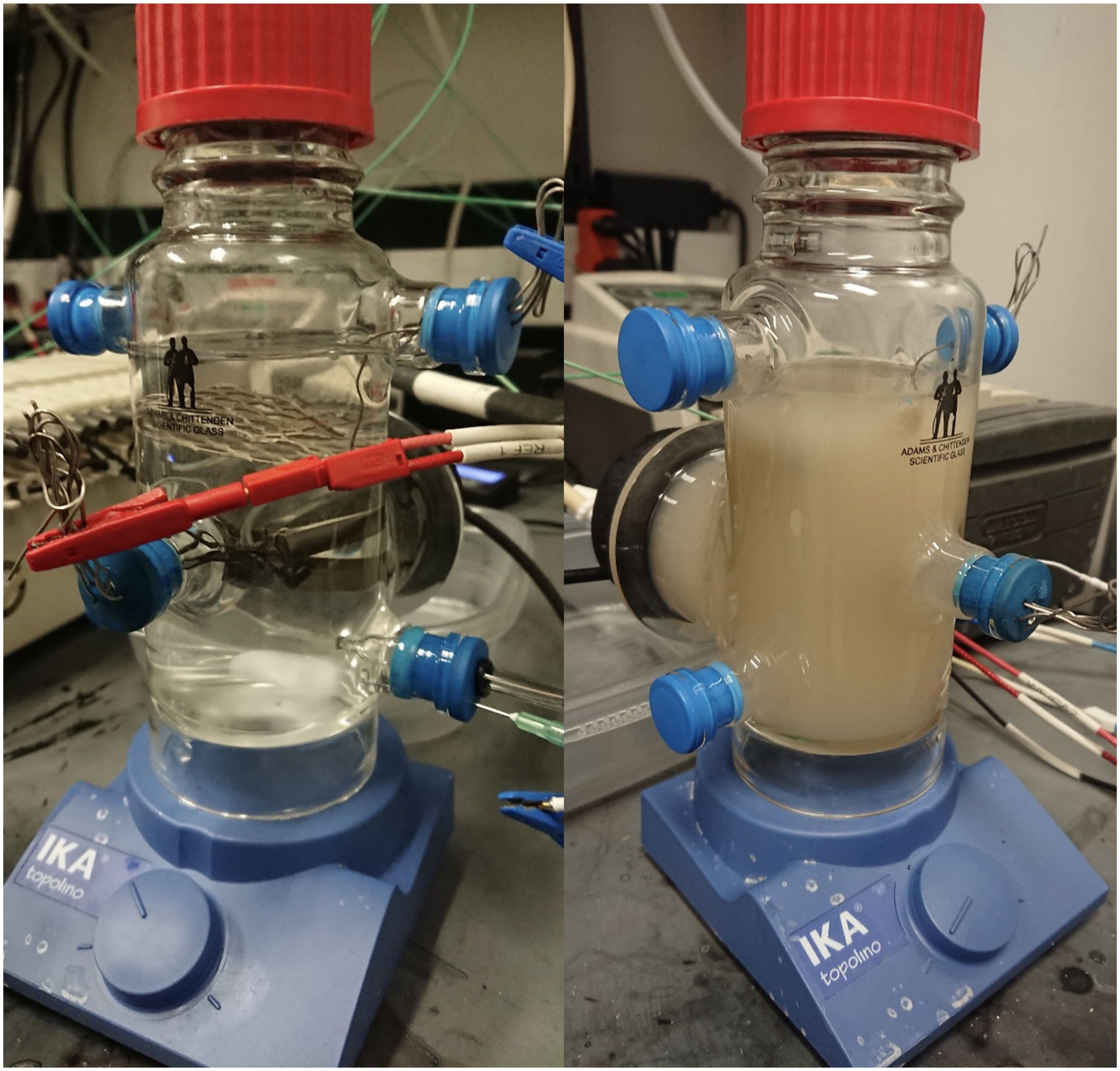


**Figure 7.** Cultivation vessel allowing autotrophic growth of *Cupriavidus necator* H16 on carbon dioxide and electrochemically formed oxyhydrogen. Left: abiotic system before inoculation. Right: operational bioreactor with fully grown *C. necator* H16 Δ*phaC1 hadA-phaC1437* culture.

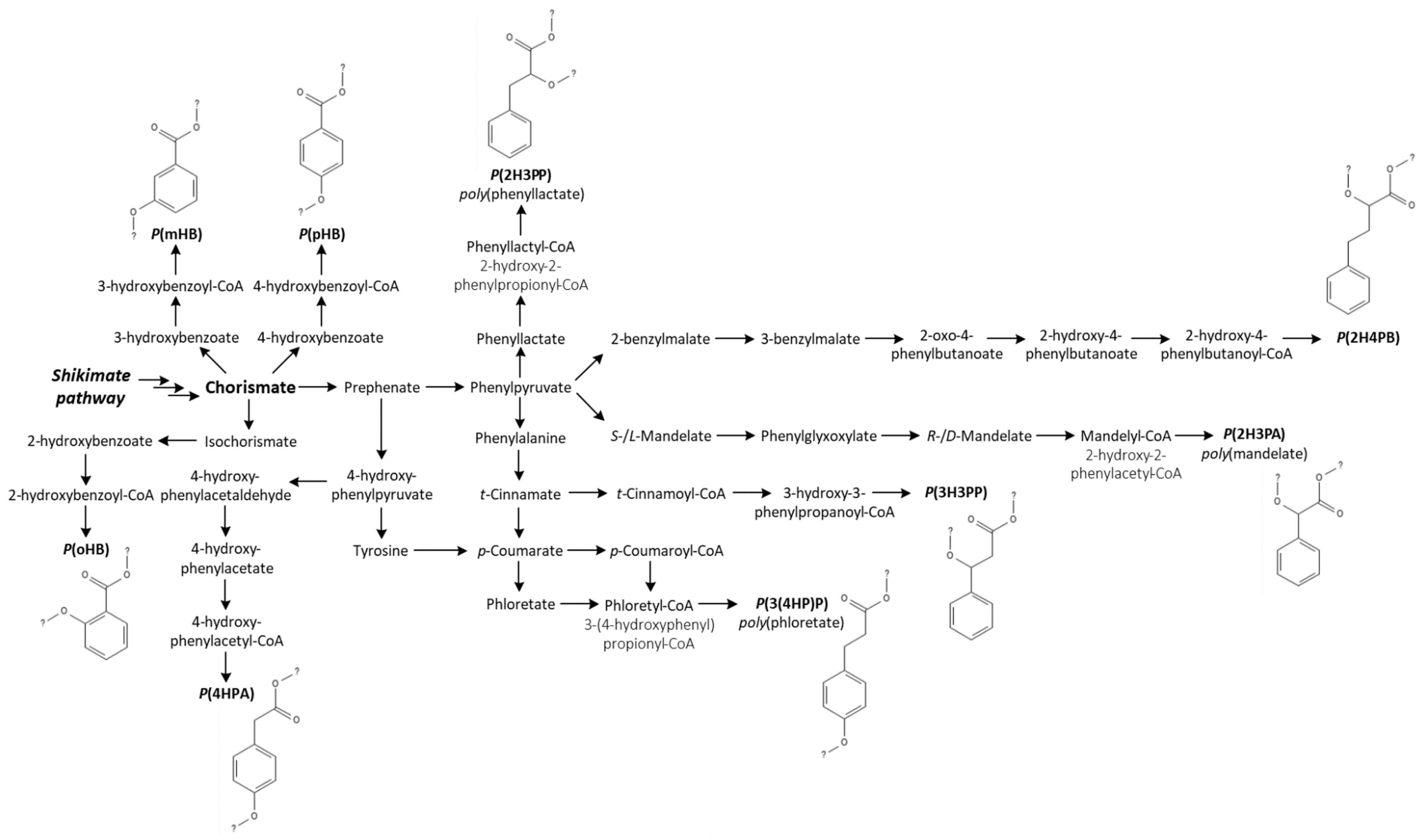


**Figure 8.** Pathways for production of aromatic PHAs from the shikimate pathway (simplified, non-stoichiometric).
Abbreviations: P3H2PP = poly(3-hydroxy-3-phenylpropanoate), P2H4PB = poly(2-hydroxy-4-phenylbutanoate), P2H3PP = poly(2-hydroxy-3-phenylpropanoate) [poly(phenyllactate)], P2H3PE = poly(2-hydroxy-3-phenylethanoate) [poly(mandelate)], P3(4HP)P = poly(3-(4-hydroxyphenyl)propanoate) [poly(phloretate)], P4HPA = poly(4-hydroxyphenylacetate), PoHB = poly(*ortho*-hydroxybenzoate), PmHB = poly(*meta*-hydroxybenzoate), PpHB = poly(*para*-hydroxybenzoate).

### 5 Maximum theoretical carbon-yields of bio-polyester production as per from metabolic network modeling

| Substrate → | **Fructose** | | **Glycerol** | | **Acetate** | | **Formate** | | **$CO_2$+$H_2$** | |
|---|---|---|---|---|---|---|---|---|---|---|
| Polyester ↓ | maximum theoretical carbon-yields [% Cmol] | | | | | | | | | |
| poly(2-hydroxyacetate) = poly(glycolate) | 83 | 81 | 88 | 80 | 64 | 50 | 19 | 16 | 100 | 85 |
| poly(2-hydroxypropanoate) = poly(lactate) | 89 | 76 | 100 | 99 | 62 | 51 | 18 | 11 | 100 | 74 |
| poly(3-hydroxypropanoate) | 83 | 77 | 97 | 85 | 56 | 50 | 17 | 5 | 100 | 47 |
| poly(3-hydroxybutyrate) | 78 | 74 | 82 | 75 | 63 | 25 | 14 | 12 | 100 | 90 |
| poly(4-hydroxybutyrate) | 78 | 72 | 92 | 82 | 55 | 49 | 18 | 12 | 100 | 90 |
| poly(3-hydroxyvalerate) | 83 | 77 | 91 | 83 | 60 | 51 | 15 | 9 | 100 | 70 |
| poly(5-hydroxyvalerate) | 74 | 67 | 85 | 77 | 50 | 47 | 14 | 12 | 100 | 94 |
| poly(3-hydroxycaproate) | 74 | 72 | 78 | 74 | 55 | 28 | 13 | 11 | 100 | 92 |
| poly(6-hydroxycaproate) | 69 | 68 | 74 | 72 | 48 | 38 | 13 | 6 | 100 | 69 |
| poly(3-hydroxy-3-phenylpropanoate) | 75 | 73 | 92 | 89 | 50 | 45 | 16 | 12 | 100 | 92 |
| poly(2-hydroxy-4-phenylbutanoate) | 79 | 77 | 87 | 84 | 54 | 41 | 15 | 14 | 100 | 96 |
| poly(2-hydroxy-3-phenylpropanoate) = poly(phenyllactate) | 75 | 72 | 92 | 89 | 50 | 42 | 16 | 13 | 100 | 93 |
| poly(2-hydroxy-3-phenylacetate) = poly(mandelate) | 69 | 68 | 83 | 79 | 47 | 36 | 14 | 12 | 100 | 92 |
| poly(3-(4-hydroxyphenyl)propanoate) = poly(phloretate) | 75 | 72 | 92 | 89 | 50 | 42 | 16 | 13 | 100 | 93 |
| poly(4-hydroxyphenylacetate) | 76 | 71 | 87 | 83 | 51 | 38 | 15 | 12 | 100 | 81 |
| poly(2-hydroxybenzoate) = poly(*orth*-hydroxybenzoate) | 77 | 73 | 95 | 87 | 51 | 37 | 16 | 14 | 100 | 91 |
| poly(3-hydroxybenzoate) = poly(*meta*-hydroxybenzoate) | 77 | 73 | 95 | 87 | 51 | 37 | 16 | 14 | 100 | 91 |
| poly(4-hydroxybenzoate) = poly(*para*-hydroxybenzoate) | 77 | 73 | 95 | 87 | 51 | 37 | 16 | 14 | 100 | 91 |
| Biomass (for comparison, not a polyester) | 74 | | 87 | | 49 | | 15 | | 100 | |

**Table 5.** Numeric values of all theoretical maximum theoretical carbon-yields in % [C-mol/C-mol] of bio-polyesters as per Elementary Flux-Mode Analysis.
